# Diversifications of both the three domains of life and SARS-CoV-2 possibly driven by biases between amino acid biosynthetic families

**DOI:** 10.64898/2026.06.22.733698

**Authors:** Dirson Jian Li

## Abstract

All cellular life forms fall under the three-domain classification of life, raising a fundamental evolutionary question: why does this classification feature three rather than two or four? To answer this question, a more general method, rather than the traditional one based on comparing small-subunit ribosomal RNAs, is required. The three-base periodicity in genomes is a common feature of both cellular life forms and viruses, which is species-specifically biased between amino acid biosynthetic families. Based on comparing the corresponding proteomic feature among all life forms, a global triangular diversification picture has been obtained, whose three angular regions correspond to the three domains, respectively. This mechanism of diversification of life attributes the evolutionary driving forces behind the diversification of the three domains of life to the biases between amino acid biosynthetic families. Notably, the same mechanism also applies to the contemporary diversification of SARS-CoV-2, whose reasonable results in turn corroborate the above explanation of primordial diversification of life and in addition shed light on the mechanism of speciation.

## Introduction

This paper aims to explain the driving forces behind the diversification of the three domains of life based on numerous genomes and proteomes. Carl Woese’s great expectation was to explain the primordial evolution of life based on sequences of contemporary species. He proposed the three domains of life and posed a series of challenging problems for his adherents. Here, I propose a statistical scenario, based on species-specific genome and proteome features, to reconstruct the three-domain classification of life and further reveal the profound relationship between the number of domains of life and the number of amino acid biosynthetic families, grounded on the exquisite concordance between observations and simulations. Intriguingly, this statistical method also applies to the ongoing diversification of SARS-CoV-2, which enables the validity of this primordial evolutionary theory to be validated against contemporary evolutionary events.

The evolution of life exhibits stochasticity [1] [2] [3]. Focusing on primordial relics can inject certainty into the study of early evolution of life [4] [5] [6] [7]. The common features of contemporary species are relics of their ancestors. The small-subunit ribosome (SSU) in all cellular life forms is a primordial relic, by comparing which the three domains of life have been found [8]. However, the method based on SSU rRNA has limitations [9] [10]: it is not general in that it cannot apply to viruses; it is not primordial enough in that it can hardly reveal the prebiotic evolution; and it is not complete in that it does not contain all the genome information. Hence, a deeper understanding of the evolution of life requires focusing on a more general, more primordial, and more complete common feature of life. The common feature of both cellular life forms and viruses is the three-base periodicity (TBP) in genomes (Supp. Fig. 4a) [11] [12] [13], whose origin relates to the origin of the triplet genetic code [14] [15] [16] [17] [18] [19]. And cellular life forms and viruses use same canonical amino acids. Evidence shows that amino acid biosynthetic families played an essential role in the evolution of the genetic code [20] [21]. The three-base periodicity biased between amino acid biosynthetic families (TBP Family Bias) is a more important primordial relic to be focused on. Moreover, there is an evolutionary relationship between TBP and SSU, considering the match between the size of anti-codon loop of tRNA and the size of SSU. In a global picture of diversification of life based on comparing species-specific TBP Family Bias, bacteria, archaea, eukaryotes, and viruses are distributed separately [22]. Although the three domains of life have been found based on SSU rRNA, the mechanism of their diversification should be revealed alternatively based on TBP Family Bias. The TBP Family Bias method provides an opportunity to clarify the driving force behind the emergence of each domain of life, to explain why the number of domains is three, and to elucidate the inherent diversification at different taxonomic levels as well as the speciation.

Advantages of the TBP Family Bias method to understand the diversification of life are as follows. TBP Family Bias at the genome level provides more comprehensive evolutionary information than SSU rRNA and the like at the gene level. To be specific, the spatial correspondence between codons in gene and amino acid residues in protein can be reconceptualized as the spatiotemporal correspondence between trinucleotide (codon) interval in genome and amino acid residue interval in proteome (Supp. Fig. 4dA, 4dB). The temporality involved here is reflected in the fact that, from a genomic perspective, the codon spacing determines the time interval for tRNA reuse, while, from a proteomic perspective, the efficiency of amino acid synthesis has to keep up with the demands of aminoacylation. Such a more important spatiotemporal correspondence had been overlooked. The discovery of TBP just refers to the preferential spacing of n-tuples, such as trinucleotides, by distances of multiples of three bases [11]. The distribution of the intervals between trinucleotides in a genome contains abundant evolutionary information [22]. TBP Family Bias is species-specific, which remains conservative during the genome evolution due to its additivity in sequence duplications and the consequent similarity among closely related species. Such statistical features of genomes of diverse species can be passed to features of the corresponding proteomes in the evolution of life. A successful pass of the features relies on translation efficiency. Without proper matching between sequence features and metabolic features, such efficiency can hardly be achieved. The highest-level diversification of life, identifiable through both sequence and metabolic features, is molded by biases between amino acid biosynthetic families. There exists, accordingly, a definite association between the three domains of life and amino acid biosynthetic families.

In this new scenario, genomes and proteomes exhibit species-specific features associated with amino acid biosynthetic families. Accordingly, a global picture of diversification of life can be obtained by comparing TBP Family Bias among numerous representative bacteria, archaea, eukaryotes, and viruses. Only in the global picture can we gain insight into the mechanism of the highest level diversification of life, and identify the constraint relationship between the number of domains of life and the number of amino acid biosynthetic families. The global picture based on species-specific TBP Family Bias is approximately a tetrahedron, whose four angular regions correspond to four of the five biosynthetic families, respectively (Fig. 1a; Supp. Fig. 4dD; Tab. 1). In this tetrahedron, high bias in the four angular regions brings about sufficiently high translation efficiency, while low bias in the central region brings about insufficient translation efficiency. Namely, archaea and eukaryotes are distributed in two angular regions, respectively, bacteria in the other two angular regions, and viruses in the central region. After passing genome features to proteome features, the tetrahedron can be compressed into an approximate triangle, whose three angular regions correspond to the three domains of life, respectively. Such a general theory on diversification of life attributes the driving forces for diversification of the three domains of life to the biases between amino acid biosynthetic families. This mechanism also comes with the ability to explain the diversification at different taxonomic levels.

**Table 1:**
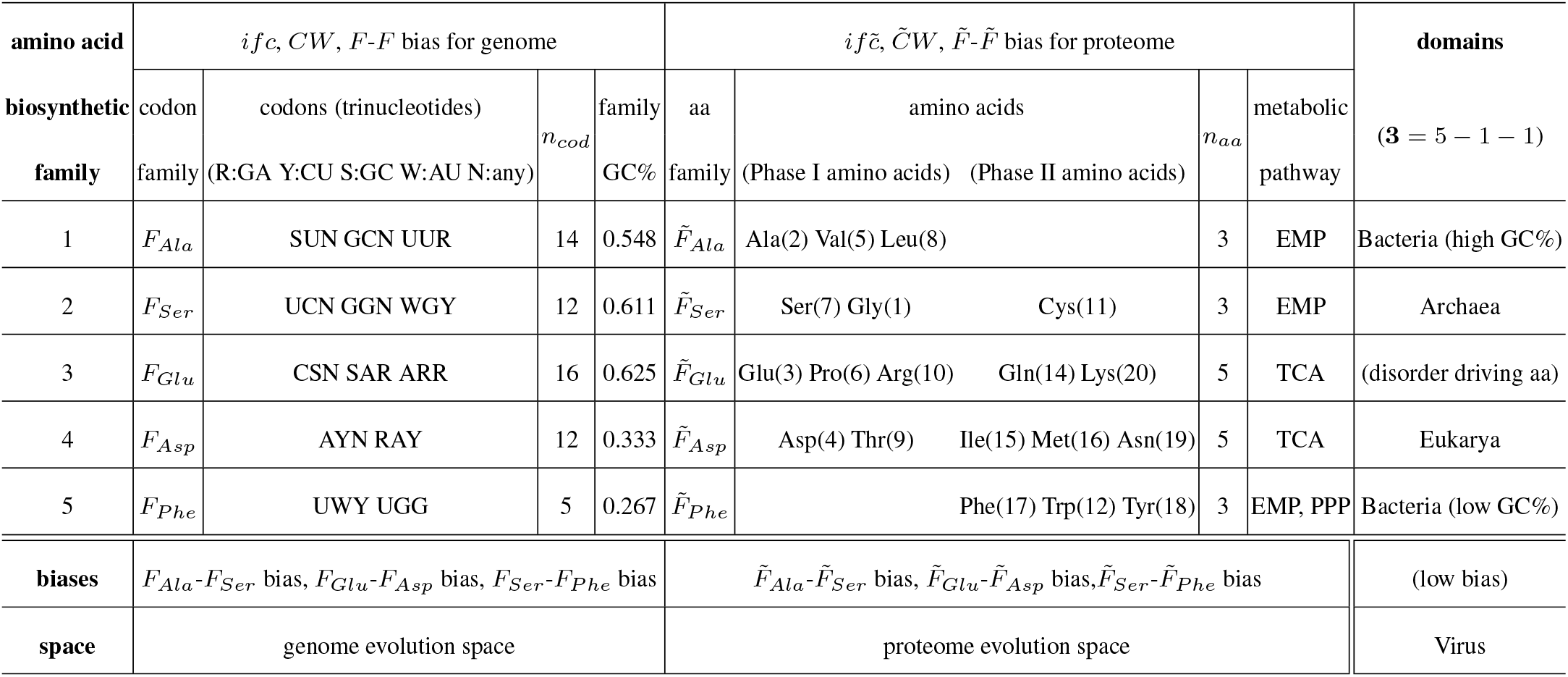
Diversification of the three domains of life driven by biases between amino acid biosynthetic families.

**Fig. 1.**
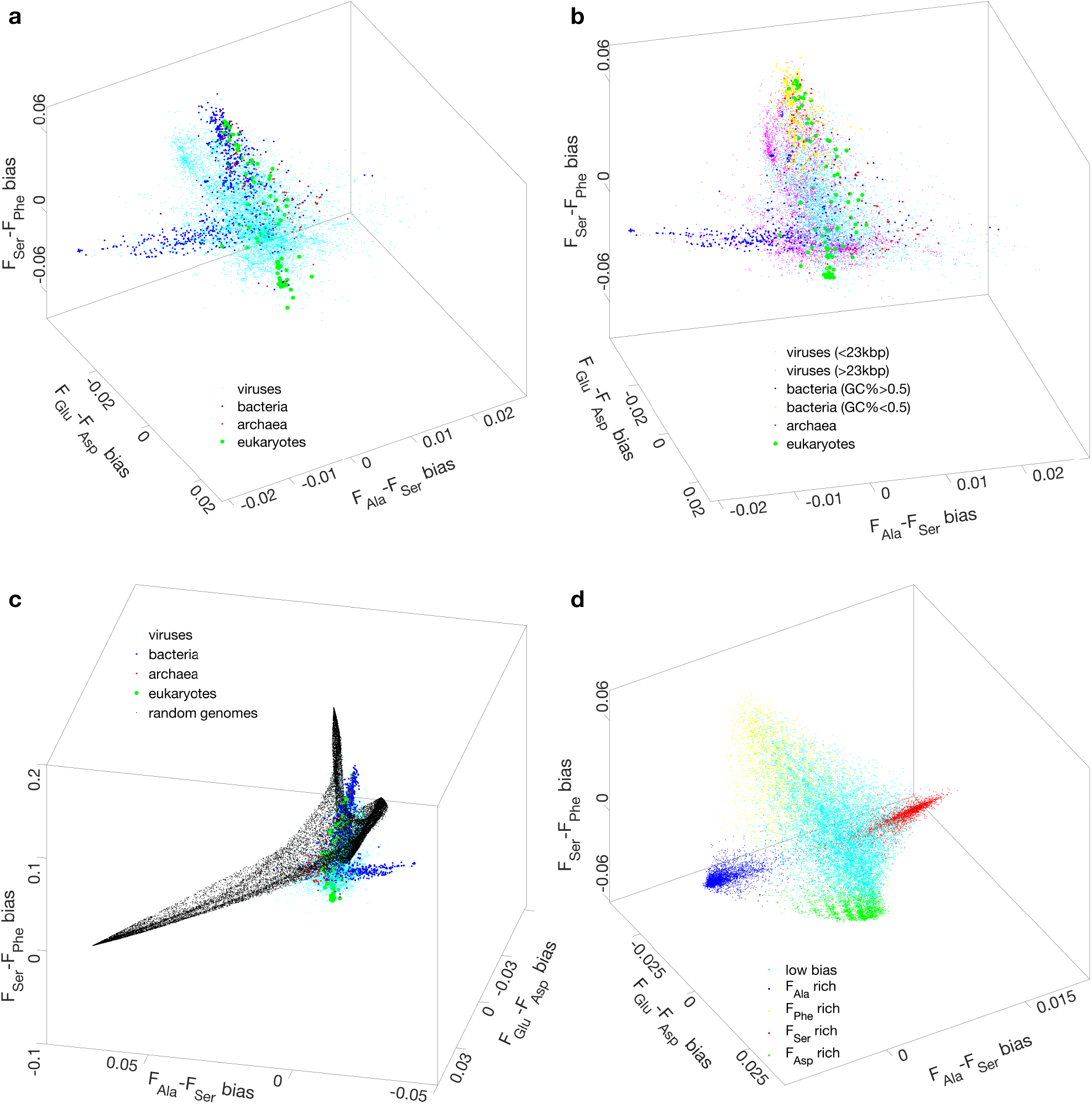
Observations and simulations of the diversification of viruses, bacteria, archaea, and eukaryotes in GenEvoSp based on TBP Family Biases in genomes. **a** The three domains of life and viruses in GenEvoSp. **b** Plaid pattern in GenEvoSp. **c** The three-base periodic genomes keep away from random sequences. **d** Simulation of diversification of viruses, bacteria, archaea, and eukaryotes based on TBP Family Bias.

Furthermore, in order to elucidate the mechanism of diversification of life, a high-resolution picture is required, instead of the previous low-resolution global picture based on species with a broad range of origin times. During the COVID-19 epidemic, it is unexpectedly for the first time to obtain a very large number of genomes for a single species, with invaluable sampling time and location information. Thus, a high-resolution picture for the real-time evolution of SARS-CoV-2 can be obtained based on comparing the similar TBP Family Bias among SARS-CoV-2 genomes with different sampling times and locations. The above general theory on diversification of the three domains of life also applies to diversification of SARS-CoV-The successful explanation of contemporary diversification of life in turn corroborates the explanation of the primordial diversification of life. Since the relationship among common features of life mirrors the relationship of the properties of primordial species, investigating the former helps to understand the latter. These common features of life contain not only TBP, the genetic code, the amino acid biosynthetic families, but also the structures of double-stranded DNA and the six reading frames. Their contributions to the early evolution of life will be discussed in detail.

In this paper, diversifications of both the three domains of life (Fig. 1) and SARS-CoV-2 (Fig. 2) have been investigated based on genome features biased between amino acid biosynthetic families, and the diversification of the three domains of life is likewise investigated based on proteome features (Fig. 3). In addition, TBP and its species-specific features have been simulated (Fig. 4).

**Fig. 2.**
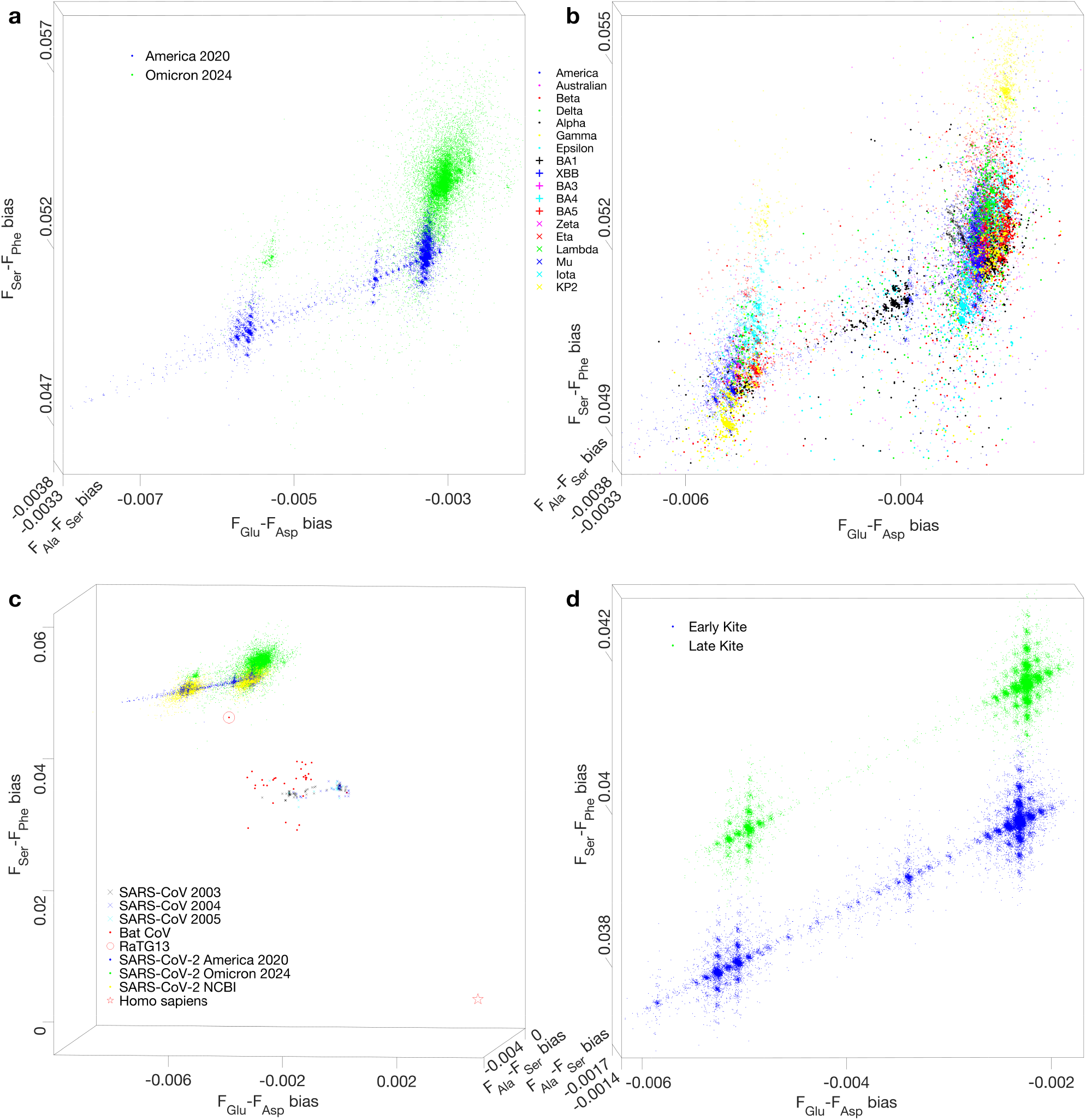
Observations and simulations of the diversification of SARS-CoV-2 in GenEvoSp based on TBP Family Biases in genomes. **a** The kite-like distributions of SARS-CoV-2 in GenEvoSp. **b** America Kite in GenEvoSp, where SARS-CoV-2 variants are marked differently. **c** The evolutionary trend from SARS-CoV to SARS-CoV-2. RaTG13 is the closest bat coronavirus to SARS-CoV-2. **d** Simulation of kite-like distribution of SARS-CoV-2 in GenEvoSp.

**Fig. 3.**
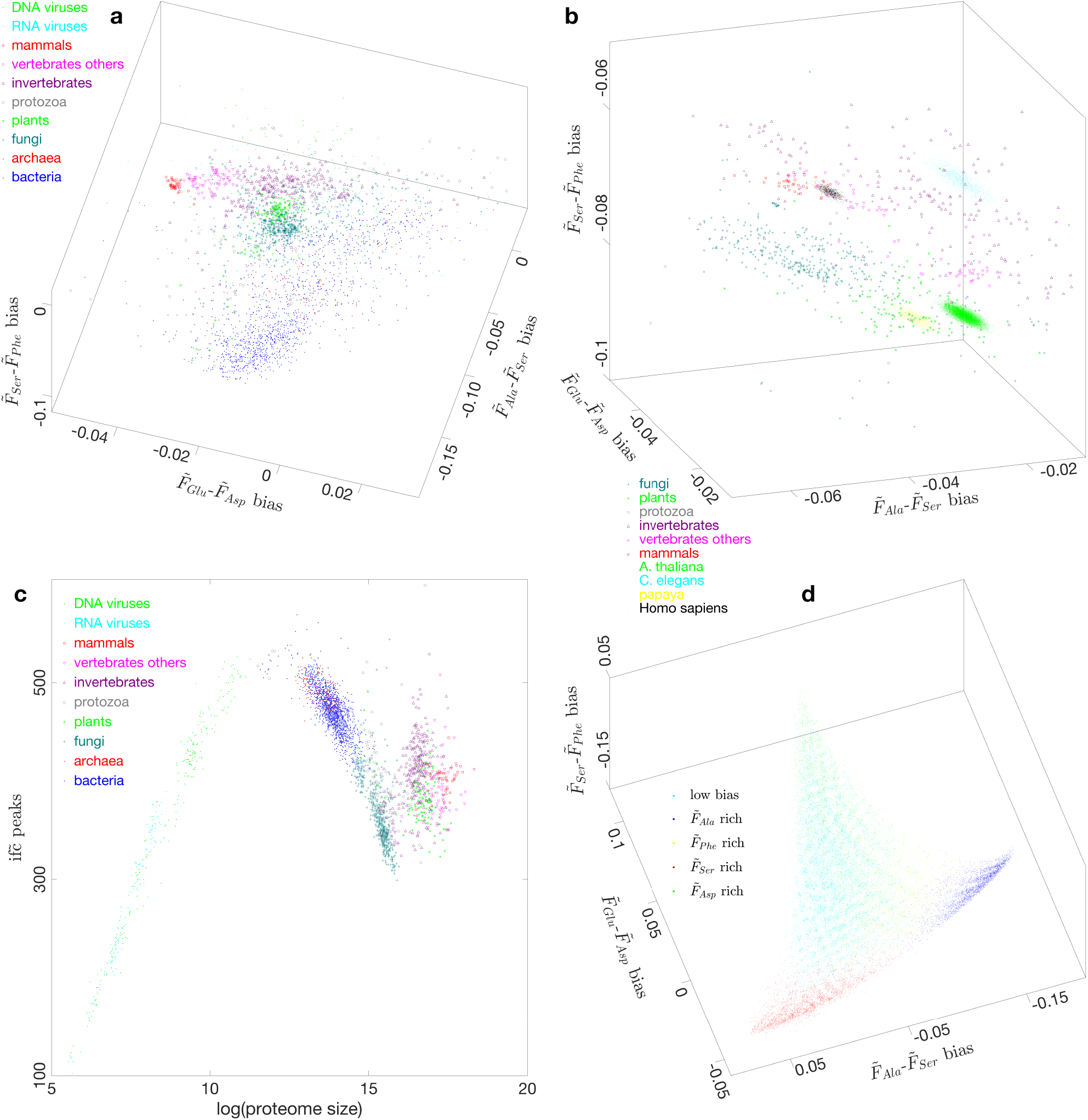
Observations and simulations of the diversification of viruses, bacteria, archaea, and eukaryotes in ProEvoSp based on IFAA Family Biases in proteomes. **a** The three domains of life in ProEvoSp, which relates to GenEvoSp according to their definitions. The sequential order of invertebrates, vertebrates excluding mammals, and mammals in ProEvoSp agrees with their evolutionary trend. **b** Separate clusters of plants and animals in ProEvoSp. The distribution of possible transcriptomes overlap the distribution region of the respective taxon. **c** Cluster analysis of DNA viruses and RNA viruses, bacteria and archaea, fungi and protozoa, plants and animals by comparing their 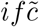. **d** Simulation of diversification of the three domains of life based on IFAA Family Biases. The triangle distribution, in ProEvoSp, of simulated proteomes translated from the 6 reading frames of the above simulated genomes.

**Fig. 4.**
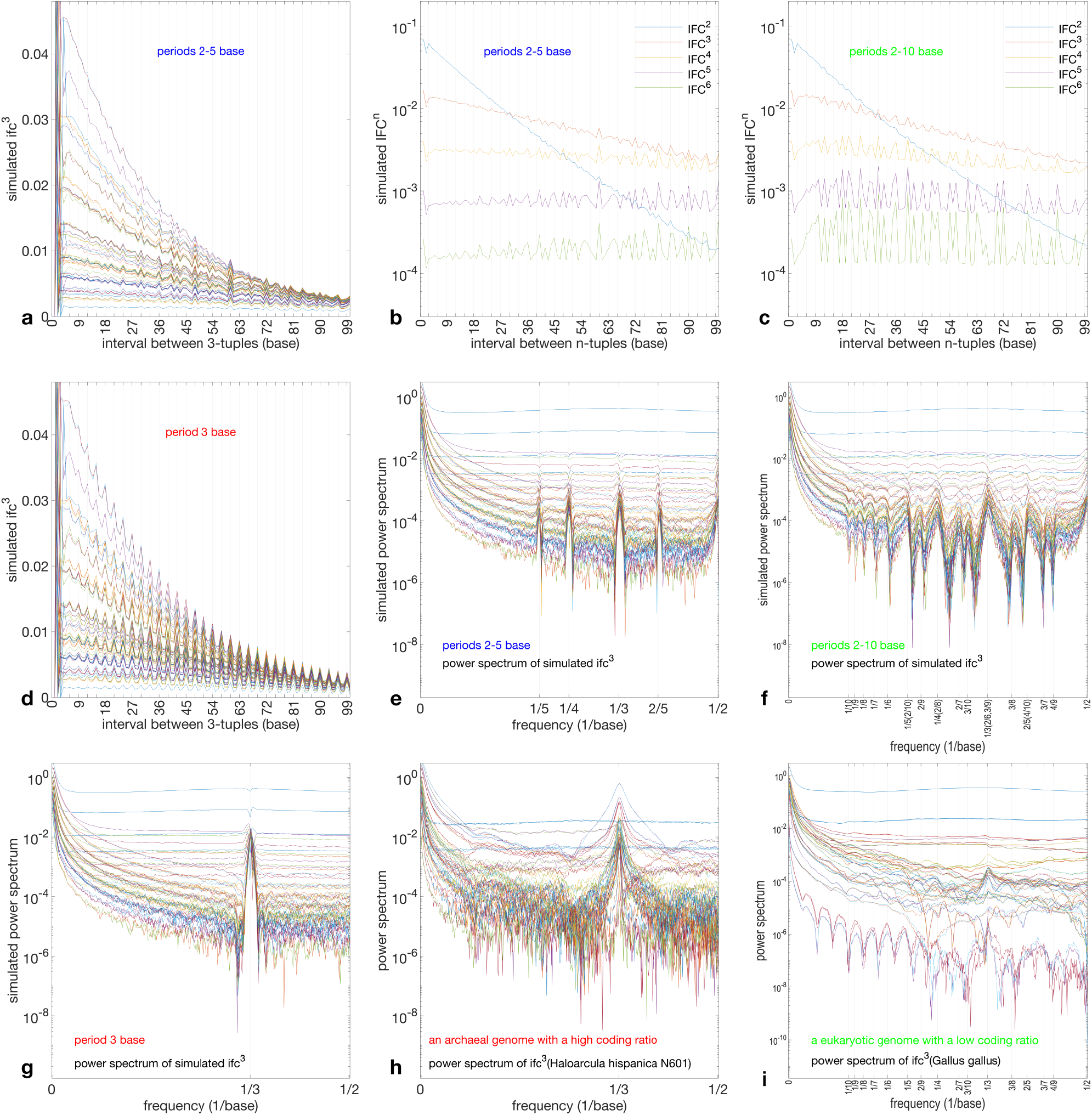
Explanation of the common feature and species-specific features of TBP in genomes. **a** Simulation of TBP in *ifc* by period averaging procedure by mixing 2-, 3-, 4-, 5-base periodic sequences. **b** Calculating *IFC* from *ifc* simulated in (a). **c** Simulation of TBP in *ifc* by period averaging procedure by mixing 2-, 3-, …, 10-base periodic sequences, and calculating IFC from ifc. **d** Simulation of TBP in *ifc* by concatenating randomly selected elements in the proper subsets of the 64 codons. **e** Power spectrum of *ifc* simulated in (a). **f** Power spectrum of *ifc* simulated in (c), which agrees with observation in (i). **g** Power spectrum of *ifc* simulated in (d), which agrees with observation in (h). **h** Power spectrum of *ifc* of Haloarcula hispanica N601. **i** Power spectrum of *ifc* of Gallus gallus.

## Methods

### Data resources

Genomes and proteomes of representative bacteria, archaea, eukaryotes, and viruses were obtained from GenBank on NCBI, and are listed in Supplementary Data 1, 3. SARS-CoV-2 genomes were obtained from COVID-19 Data Portal [23], and are listed in Supplementary Data 2. Simulated results are provided in Supplementary Data 4.

The MATLAB scripts to generate 2- and 3-dimensional Figs. 1 to 4 are in the Supplementary Information. The same set of scripts was employed for the analysis of both biological and simulated genomes/proteomes; these scripts are included in the fifth MATLAB script. I strongly recommend running these scripts to facilitate understanding of the Supplementary methods, which contain dense specialized terminology.

### Definitions of interval frequency of codon (*ifc*), codon weight (*CW*), and genome family bias (*F*-*F* bias)

These three concepts are all calculated based on the genome sequence of a certain species. TBP is quantitatively described by *ifc*, which is the normalised distribution of intervals between the respective 64 trinucleotides in a genome. The trinucleotides correspond one-to-one with the codons, both of which can be grouped in codon families according to their encoded amino acids in certain amino acid biosynthetic families such as *F*_*Ala*_, *F*_*Ser*_, *F*_*Glu*_, *F*_*Asp*_, and *F*_*Phe*_ (Tab. 1). *cw* are the weights of the respective codons in *ifc*, and these weights can also be grouped and averaged so as to obtain *CW* for the respective amino acid biosynthetic families. TBP Family Bias is quantitatively described by *F*_*fam*1_-*F*_*fam*2_ bias, which is defined as the difference of *CW* between *F*_*fam*1_ and *F*_*fam*2_ for a species.

### Definitions of interval frequency of amino acid (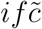**), amino acid weight (**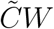**), and proteome family bias (**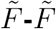 bias)

These three concepts are all calculated based on the protein sequences in the proteome of a certain species. 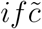 is the normalised distribution of intervals between the respective 20 amino acid residues in a proteome. The 20 amino acids can be grouped according to the amino acid biosynthetic families such as 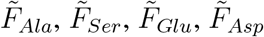, and 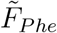 (Tab. 1). 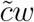 are the weights of the respective amino acids in 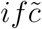, and these weights can also be grouped and averaged so as to obtain 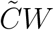 for the respective amino acid biosynthetic families. 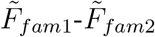 bias is defined as the difference of 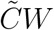 between 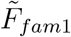 and 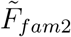 for a species.

### The genome evolution space (GenEvoSp)

GenEvoSp is the three-dimensional space whose coordinates are *F*_*Ala*_-*F*_*Ser*_ bias, *F*_*Glu*_-*F*_*Asp*_ bias, and *F*_*Ser*_-*F*_*Phe*_ bias, respectively. The genome of a species corresponds to a specific coordinate position in GenEvoSp. When studying SARS-CoV-2, a genome of SARS-CoV-2 also corresponds to a specific coordinate position in GenEvoSp. The biases for coordinates here are the best choice for clustering the three domains of life separately [22].

### The proteome evolution space (ProEvoSp)

ProEvoSp is the three-dimensional space whose coordinates are 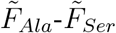 bias, 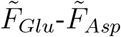 bias, and 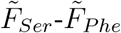 bias, respectively. The proteome of a species corresponds to a specific coordinate position in ProEvoSp. Both *F*_*Ala*_-*F*_*Ser*_ bias and 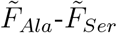 bias are also known as EMP bias, both *F*_*Glu*_-*F*_*Asp*_ bias and 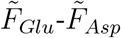 bias as TCA bias, and both *F*_*Ser*_-*F*_*Phe*_ bias and 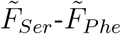 bias as GC% bias.

### The three domains of life and viruses in GenEvoSp

Genomes of bacteria, archaea, eukaryotes, and viruses correspond to sets of specific coordinate positions in GenEvoSp, respectively. The three domains of life, viruses, as well as taxa at different taxonomic levels are distinguished by different markers in GenEvoSp. The global picture of diversification of life is approximately a tetrahedron in GenEvoSp (Fig. 1a).

### The three domains of life and viruses in ProEvoSp

Proteomes of bacteria, archaea, eukaryotes, and viruses correspond to sets of specific coordinate positions in ProEvoSp, respectively. The three domains of life, viruses, as well as taxa at different taxonomic levels are distinguished by different markers in ProEvoSp. The global picture of diversification of life is approximately a triangle in ProEvoSp (Fig. 3a).

### SARS-CoV-2 in GenEvoSp

Genomes of SARS-CoV-2, as well as SARS-CoV, and bat coronaviruses, correspond to sets of specific coordinate positions in GenEvoSp, respectively. The variant strains of SARS-CoV-2 with sampling times and locations are marked differently in GenEvoSp. The distribution shape of SARS-CoV-2 with certain sampling time and location (denoted as Location-time Kite) drifts over time in GenEvoSp (Fig. 2a).

### Simulation of the three domains of life and viruses in GenEvoSp

Numerous simulated three-base periodic genomes, with different probability for selecting codons for the respective amino acid biosynthetic families, are generated, whose distribution shape is a tetrahedron in GenEvoSp. And the four angular regions of the tetrahedron correspond to four amino acid biosynthetic families, respectively. *F*_*Glu*_ is adjacent to the tetrahedron (Supp. Fig. 4dD). Four angular regions around vertices of this tetrahedron, *F*_*Ala*_, *F*_*Phe*_, *F*_*Ser*_, and *F*_*Asp*_, correspond to bacteria with high GC content, bacteria with low GC content, archaea, and eukaryotes, respectively (Fig. 1d).

### Simulation of the three domains of life and viruses in ProEvoSp

The simulated proteomes are obtained by translating the six reading frames of each of the above simulated genomes. The distribution shape of the simulated proteomes is approximately a triangle in ProEvoSp. Two vertices *F*_*Ala*_ and *F*_*Phe*_ of the tetrahedron in GenEvoSp can be combined into one vertex of the triangle in ProEvoSp, due to translating from six rather than one reading frame. Thus, the three angular regions of the triangle in ProEvoSp correspond to the three domains of life, respectively, i.e., *F*_*Ala*_ and *F*_*Phe*_ to Bacteria, *F*_*Ser*_ to Archaea, and *F*_*Asp*_ to Eukarya (Fig. 3d).

### Simulation of SARS-CoV-2 in GenEvoSp

A simulated three-base periodic genome can be generated first with similar GC content of SARS-CoV-2. Taking this genome as the initial SARS-CoV-2 genome, and introducing specific mutations in a small proportion of the initial genome, the Location-Time Kites can be simulated (Fig. 2d).

### Simulation of TBP and its features

The initial TBP can be simulated by averaging different periods in periodic sequences. The genomes with species-specific TBP Family Bias can be simulated based on concatenating codons randomly selected from different subsets of the 64 codons (Fig. 4).

## Results

Based on the SSU rRNA method, all cellular life forms have been divided into Bacteria, Archaea, and Eukarya. Here, based on the TBP Family Bias method, a mechanism of diversification of the three domains of life will be explained from the following five aspects: (1) the sequence features of each of the three domains of life, (2) the application of this general mechanism to the diversification of SARS-CoV-2, (3) the application of this general mechanism to the diversification at different taxonomic levels as well as the speciation, (4) the sequence features of cellular life forms that differ from viral sequences and non-living sequences, and (5) some detailed coincidences between biological observations and simulations.

### Diversification of the three domains of life

Bacteria, archaea, and eukaryotes correspond to the four vertices of the tetrahedron in GenEvoSp respectively (Fig. 1a), as well as to the three vertices of the triangle in ProEvoSp respectively (Fig. 3a). The driving force to separate bacteria and archaea comes from *F*_*Ala*_-*F*_*Ser*_ bias, and the driving force to separate archaea and eukaryotes from *F*_*Glu*_-*F*_*Asp*_ bias. Bacteria correspond to the two vertices *F*_*Ala*_ and *F*_*Phe*_ in GenEvoSp, as well as to the combination of 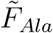 and 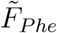 vertex in ProEvoSp (Fig. 1a, 3a). *F*_*Phe*_ with the lowest family GC content (Tab. 1) corresponds to bacteria with low GC content, and *F*_*Ala*_ corresponds to bacteria with high GC content (Fig. 1b). Archaea correspond to the vertex *F*_*Ser*_ in GenEvoSp as well as to the vertex 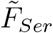 in ProEvoSp (Fig. 1a, 3a). Eukaryotes correspond to the vertex *F*_*Asp*_ in GenEvoSp as well as to the vertex 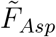 in ProEvoSp (Fig. 1a, 3a). There is a close statistical relationship between *ifc* of genome and 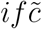 of proteome. Separate clusterings of the three domains of life have been observed in both GenEvoSp and ProEvoSp (Supp. Fig. 4c).

The global pictures of diversification of life in GenEvoSp and ProEvoSp illustrate that the number for highest level classification of life is three. Traditionally, the degeneracy of the genetic code refers to the mapping between the 64 codons and the 20 canonical amino acids as well as termination codons. Considering that amino acids belong to respective biosynthetic families, there is a “codon-to-family generalized degeneracy” as the mapping between the 64 codons and the 5 biosynthetic families as well as others (Tab. 1). The histidine can be temporarily ignored due to its small proportion. All in *F*_*Glu*_ are disorder-driving amino acids [24] [25] [26] [27] [28], such that *F*_*Glu*_-rich genomes tend to encode poorly folded proteins and hence do not represent cellular life forms. And the translation from genome to proteome combines 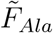 and 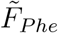 into one group of amino acids (Fig. 3a, 3d). Therefore, 3 groups of amino acids remain (3=5-1-1), corresponding to Bacteria, Archaea, and Eukarya, respectively.

### Diversification of SARS-CoV-2

TBP is also a common feature in all viral genomes. The statistical method based on TBP Family Bias can apply to the diversification of SARS-CoV-2. The information of sampling time and location provides a chance to study the real-time evolution of SARS-CoV-2 in GenEvoSp, according to drifting distribution of SARS-CoV-2 over time. The term Kite denotes the distribution of SARS-CoV-2 with certain sampling time and location in GenEvoSp. Different variant strains are distributed separately in certain parts of a Kite (Fig. 2b). The evolutionary relationships among different Kites can be elucidated based on their drifts in GenEvoSp (Supp. Fig. 2aC, 2dC). There is a unidirectional evolutionary trend from SARS-CoV, to bat coronavirus, to early SARS-CoV-2, to Omicron, and to KP2 in GenEvoSp, which tends to keep away from Homo sapiens (Fig. 2c). RaTG13 is the closest bat coronavirus to SARS-CoV-2 (Fig. 2c), which is a close agreement between the TBP Family Bias method and the results in the literature [29].

A plaid pattern, namely, the three-dimensionally periodic density of SARS-CoV-2 in GenEvoSp, has been observed in a high-resolution Kite based on numerous SARS-CoV-2 genomes. The term subcluster denotes a check in the plaid pattern of a Kite, namely the small region with continuously high density. For example, America Kite (Supp. Fig. 2bC) consists of short-tail array (Fig. 2b), long-tail array (Supp. Fig. 2bB), and middle-tail array (Supp. Fig. 2bA) [30] [31], each of which has the plaid pattern. The formation of America Kite over time has been reproduced based on the information of sampling time for each of the genomes (Supp. Fig. 2cA, 2cB). America Kite began growing from one side of the middle-tail array (Supp. Fig. 2cA, 2cB), and eventually spread throughout its entire region. Notably, the SARS-CoV-2 genome size underwent greater changes during the initial period than in subsequent periods (Supp. Fig. 2cB). The short-tail array and the long-tail array evolved concurrently (Supp. Fig. 2cB). The number of adenines in Poly(A) tail of SARS-CoV-2 increases by 1 per adjacent subcluster along the axis of symmetry of Kite (Supp. Fig. 2bD). There is an obvious interleaved arrangement between the subclusters of America Kite and subclusters of Australia Kite (Supp. Fig. 2aB).

### Diversification at different taxonomic levels

The TBP Family Bias method also applies to different taxonomic levels. The plaid pattern in the high-resolution Kite may enlighten our understanding of the mechanism of speciation. Variant strain generally corresponds to a set of subclusters (Fig. 2b). Elucidating the mechanism underlying the separation of subclusters in a Kite helps reveal the mechanisms of speciation.

The mechanism underlying subcluster separation can be clearly explained via simulation of plaid pattern in a Kite (Fig. 2d). The separation stems from the uneven correspondence between the randomly changed genome sequences and the plaid pattern, or step change in density of species, in GenEvoSp and ProEvoSp, which arises from the codon-to-family generalized degeneracy. Specifically, under random sequence variation and given the similarity of codons belonging to the same biosynthetic family, only a minority of changes of codons can bring about shifts between biosynthetic families. For examples, the changes of codons from CAR to RAC bring about shift from *F*_*Glu*_ to *F*_*Asp*_, while the majority of changes of codons keep them within their original biosynthetic families; the changes of the third base of SUN and GCN in *F*_*Ala*_ keep them within *F*_*Ala*_.

Large-scale plaid patterns have been observed in the distributions of species in GenEvoSp and ProEvoSp, e.g., the stripe structures near vertices (Fig. 1b; Supp. Fig. 1b) and distinct clustered regions in ProEvoSp (Fig. 3a, 3b; Supp. Fig. 3a). Unlike the clear plaid patterns in SARS-CoV-2 Kites, these patterns are less prominent for cellular life forms, due to the wide range of origin times across these diverse species. As the metabolic variations exhibit a stable correlation with the coordinates of GenEvoSp and ProEvoSp, the plaid patterns indicate a step correlation between the metabolism variation and the sequence variations, which accounts for the diversifications at different taxonomic levels.

Plaid patterns have been observed clearly in the simulated tetrahedron in GenEvoSp (Fig. 1d) and the simulated triangle in ProEvoSp (Fig. 3d; Supp. Fig. 3dC, 3dD). Elucidating the origin of large-scale plaid patterns is critical to revealing the mechanisms underlying diversification of life at different taxonomic levels, including phylum, class, etc. The simulation demonstrates a scale-free mechanism of diversification, in either the small-scale plaid patterns in SARS-CoV-2 Kites or the large-scale plaid patterns in the tetrahedron and the triangle for diverse species. Specifically, the long-/medium-/short-term evolution can bring about changes in large/medium/small proportions of genomes, which result in large-/medium-/small-scale plaid patterns for the higher/middle/lower level taxa. Speciation is due to the smallest-scale plaid pattern by the same mechanism.

### Sequence features of cellular life forms

The difference between cellular life forms and viruses can be identified, as the Go proverb goes “Gold corners, silver sides, grass belly”. Namely, cellular life forms are distributed in the four angular regions (gold corners) with high *F*-*F* bias, while viruses in the central region (grass belly) with low *F*-*F* bias, in the tetrahedron in GenEvoSp (Fig. 1a) (see Method). In general, the magnitude of *F*-*F* bias is positively correlated with translation efficiency. High *F*-*F* bias leads to more use of codons for certain subsets of all amino acids, and results in high efficiency for tRNA reuse. Finding suitable tRNAs from fewer types around the ribosome can shorten the time interval for tRNA reuse, and hence improve translation efficiency. The directions extending from the center to the four corners indicate a potential increase in translation efficiency. The viruses in the grass belly cannot achieve autonomous translation possibly due to insufficient efficiency.

TBP is a common genome feature of both cellular life forms and viruses. The distribution of all kinds of real life forms a tetrahedron in GenEvoSp, whose four vertices correspond to the four amino acid biosynthetic families (Supp. Fig. 4dD). The distribution of all random sequences forms a trident star (Fig. 1c). The trident star has intercepted the tetrahedron (Fig. 1c), such that the genomes with TBP in the tetrahedron have deviated from the curved surface of the trident star. Hence, TBP becomes a strategy of life to keep away from randomness. The vertex *F*_*Glu*_ is near to the G-rich vertex of the trident star (Supp. Fig. 1cD); these genomes tend to be random and do not represent cellular life forms.

### Some detailed observations and simulations

The validity of this general mechanism depends on not only the above analysis but also the detailed agreement between simulations and observations. In detailed observation of real life, there is a shift of the bacteria-archaea edge of the tetrahedron to the negative side of the origin of the coordinate *F*_*Glu*_-*F*_*Asp*_ bias in GenEvoSp (Supp. Fig. 1aC). In simulation, there is no such a shift if mistakenly reading only single strand of DNA (Supp. Fig. 1dA), while there does be such a shift if correctly reading both strands of DNA (Supp. Fig. 1dB, 1aD). Such a shift in the simulation stems from the codon-to-family generalized degeneracy; specifically, the reverse complementary codons of *F*_*Ala*_ and *F*_*Ser*_ encode more amino acids in 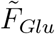.

In another detailed observation of real life, the distribution shape of species is an approximate triangle in ProEvoSp (Fig. 3a). In simulation, the distribution shape of simulated species is still a tetrahedron in ProEvoSp if mistakenly reading only one reading frame (Supp. Fig. 3dA), while the distribution shape of simulated species becomes a triangle in ProEvoSp if correctly reading the six reading frames (Fig. 3dB). Such an agreement also stems from the codon-to-family generalized degeneracy. Specifically, the codons in *F*_*Ala*_ can be roughly represented by NUN, while the codons in *F*_*Phe*_ roughly by UNN. The ambiguity between NUN and UNN can be achieved by reading-frame shift. This accounts for the combination of *F*_*Ala*_ and *F*_*Phe*_ in the tetrahedron in GenEvoSp into the single vertex in the triangle in ProEvoSp.

## Discussion

### Connection and comparison between SSU rRNA method and TBP Family Bias method

Interrelationship among common features of life is a golden key to unlocking the mechanism of diversification of life. Both SSU rRNA method and TBP Family Bias method relate to translation process, and particularly to the origin of the triplet genetic code. TBP and the triplet genetic code are two sides of the same triplet nature of life. The core significance of choice of the triplet nature of life lies in the fact that it sets a standard unit of length for nucleic acid sequences. Such a three-base length unit requires a match among the size of SSU, the size of tRNA anticodon loop, and the size of adjacent codons in mRNA. All these would have dramatically changed if two-or four-base length unit had been chosen. The high efficiency of the translation process is a vital factor for species survival. TBP, which relates to the origin of the genetic code and applies to viruses besides cellular life forms, should be more primordial than SSU rRNA. The variation of TBP Family Bias needs the corresponding variation of SSU rRNA to maintain the efficiency of translation.

Although both TBP Family Bias method and SSU rRNA method can separate the three domains of life as well as taxa at different taxonomic levels, only the former is able to explain the separation between cellular life forms and viruses. The advantage of TBP Family Bias method stems from its direct relationship with the amino acid biosynthetic families, while the SSU rRNA method cannot see through this relationship. The TBP Family Bias method can reveal the relationship between the three domains of life and amino acid biosynthetic families, and the correlation between the equiprobable random variation in sequences and the step variation in metabolism features. Therefore, biases between amino acid biosynthetic families can provide the driving forces for diversification of life at the domain level as well as at the levels lower than domains.

### Why do cellular life forms have three domains? A hypothesis

Evidence shows that a unified mechanism underlies the diversification of life. The advantage of the TBP Family Bias method over the SSU rRNA method depends on its insights into the mechanism of the diversification of life. The concordance between observations and simulations reveals that the number of classification of life at the highest level stems from the number of amino acid biosynthetic families. Among the 5 amino acid biosynthetic families in Tab. 1, *F*_*Glu*_ corresponds to disorder driving amino acids. The simulated *F*_*Glu*_-rich genomes are distributed in the region adjacent to the tetrahedron (Supp. Fig. 4dD), where some viruses can also be found (Fig. 1a). The remaining 5-1 amino acid biosynthetic families correspond to the 4 vertices in the tetrahedron in GenEvoSp, two of which are combined into one vertex in the triangle in ProEvoSp (Fig. 1a, 3a, 1d, 3d). So there are 4-1 vertices in the triangle based on proteome features, which correspond to the 3 domains of cellular life forms (Fig. 3a, 3d). Statistics has shaped the global tetrahedral/triangular distributions of genome/proteome features; diverse genomes/proteomes occupy all regions within the global shapes.

Theories of early evolution are prone to controversy and difficult to subject to direct empirical testing. Thankfully, the TBP Family Bias method, a universally applicable statistical approach, was corroborated indirectly by the contemporary evolutionary event. This general statistical method attributes diversifications at both the domain level and the levels below domains to the uneven impact of sequence variation on metabolism variation, according to large-or small-scale plaid patterns in the global tetrahedron or triangle. Moreover, the TBP Family Bias method applies to viruses, placing their evolutionary status in the remaining central area of the tetrahedron. Owing to the proposed same mechanism of diversification of life for both long- and short-term evolution, the rapid evolutionary event of SARS-CoV-2 provides a chance to corroborate this statistical method.

### Validity of the theory depends on detailed agreements

Life in nature inherently undergoes diversification, and it is not surprising that a certain quantitative method based on sequence features achieves separate clustering. It is necessary to clarify that the mechanism here is the cause rather than the result. Some of the most striking results in this paper emerged from serendipity; details make all the difference. Some critical details are as follows.

The first detail comes from the agreement between the theory and the amino acid recruitment in the prebiotic evolution. The positive directions of the three coordinate axes in GenEvoSp point towards the direction of time, according to the proportions of the early Phase I amino acids over the late Phase II amino acids in pairs of amino acid biosynthetic families in calculating *F*-*F* biases (Tab. 1) [20] [32]. It is therefore reasonable to obtain both the positive direction from bacteria to archaea, and the positive direction from archaea to eukaryotes in GenEvoSp (Fig. 1a).

The second detail comes from the agreement between the theory and the assignment of codons to amino acids, as well as structures of molecules. The shift of the tetrahedron along the negative direction of the *F*_*Glu*_-*F*_*Asp*_ bias axis in GenEvoSp (Supp. Fig. 1aC) has been simulated by reading double-stranded DNA (Supp. Fig. 1dB), due to the fact that there are more codons in *F*_*Glu*_ in the complementary strand with reverse complementary codons of the union of *F*_*Ala*_ and *F*_*Ser*_. The combination of two vertices *F*_*Ala*_ and *F*_*Phe*_ in the tetrahedron in GenEvoSp (Fig. 1a, 1d) into the single vertex in the triangle in ProEvoSp (Fig. 3a, 3d) has been simulated, due to the identity between NUN roughly for *F*_*Ala*_ and UNN roughly for *F*_*Phe*_ after reading-frame shift.

The third detail comes from the agreement between the theory and the known evolutionary trends. The diversification between cellular life forms and viruses has been illustrated by a global picture. The evolutionary trend from invertebrates to mammals points towards the vertex for eukaryotes in ProEvoSp (Fig. 3a), as expected. The evolutionary trend from SARS-CoV to SARS-CoV-2 tends to keep away from Homo sapiens in GenEvoSp (Fig. 2c), and notably, RaTG13 is the closest bat coronavirus to SARS-CoV-2 (Fig. 2c).

The fourth detail comes from the agreement between the theory and the separate clusterings of taxa across all levels. It has been observed in the large-scale separate clusterings between animals and plants (Supp. Fig. 3aB), and between invertebrates and vertebrates (Supp. Fig. 3aA). It has also been observed that the boundary lines between different SARS-CoV-2 variant strains coincide with the gaps between the small-scale subclusters in the plaid patterns (Fig. 2b).

## Supplementary information

- Supplementary methods
- Supplementary figures 1a, 1b, *···*, 4c, 4d
- Supplementary table 1
- MATLAB_script_to_generate_Fig_1.m
- MATLAB_script_to_generate_Fig_2.m
- MATLAB_script_to_generate_Fig_3.m
- MATLAB_script_to_generate_Fig_4.m
- MATLAB_script_to_simulate_genomes_in_Fig_1d_and_proteomes_in_Fig_3d.m
- Supplementary_Data_1_CodonWeight.xlsx
- Supplementary_Data_2_CodonWeight SARS-CoV-2.xlsx
- Supplementary_Data_3_AminoAcidWeight.xlsx
- Supplementary_Data_4_ifaa_ifc_simulation.xlsx

Supplementary Data 1-4 are available in Mendeley Data: Li, DJ (2026) Data: Diversifications of both the three domains of life and SARS-CoV-2 possibly driven by biases between amino acid biosynthetic families, Mendeley Data, V1, doi: 10.17632/x3ghsxs98b.1, https://data.mendeley.com/datasets/x3ghsxs98b/1.

## Acknowledgements

My warm thanks go to Jinyi Li for valuable discussions.

## Use of AI tools declaration

The author declares that he has not used Artificial Intelligence tools in the creation of this article.

## References

[1] Lenormand, T., Roze, D., Rousset, F. (2009) Stochasticity in evolution. Trends in Ecology and Evolution 24(3):157–165.

[2] Som, A. (2014) Causes, consequences and solutions of phylogenetic incongruence. Briefings In Bioinformatics 16(3):536–548.

[3] Koonin, E.V. (2015) The turbulent network dynamics of microbial evolution and the statistical tree of life. J. Mol. Evol. 80:244–250.

[4] Giulio, M.D. (2026) Domains of life identified through genetic codes. BioSystems 266:105848.

[5] Zhao, M., Ding, R., Liu, Y., Ji, Z., Zhao, Y. (2022) Determination of the amino acid recruitment order in early life by genome-wide analysis of amino acid usage bias. Biomolecules 12:171.

[6] Gaydukova, S.A, Moldovan, M.A., Vallesi, A., Heaphy, S.M., Atkins, J.F., Gelfand, M.S., Baranov, P.V., (2023) Nontriplet feature of genetic code in Euplotes ciliates is a result of neutral evolution. Proc. Natl. Acad. Sci. USA 120(22): e2221683120.

[7] Castillo-Villanueva, A., Reyes-Vivas, H., Oria-Hernández, J. (2023) Comparison of cysteine content in whole proteomes across the three domains of life. PLoS ONE 18(11):e0294268.

[8] Woese, C.R., Kandler, O., Wheelis, M.L. (1990) Towards a natural system of organisms: Proposal for the domains Archaea, Bacteria, and Eucarya. Proc. Natl. Acad. Sci. USA 87:4576–4579.

[9] Stackebrandt, E., Goebel, B.M. (1994) Taxonomic note: A place for DNA-DNA reassociation and 16S rRNA sequence analysis in the present species definition in bacteriology. International Journal of Systematic Bacteriology 44(4):846–849.

[10] Bapteste, E., Susko, E., Leigh, J., MacLeod, D., Charlebois, R.L., Doolittle, W.F. (2005) Do orthologous gene phylogenies really support tree-thinking? BMC Evolutionary Biology 5:33

[11] Shepherd, J.C.W. (1981) Periodic correlation in DNA sequences and evidence suggesting their evolutionary origin in a comma-less genetic code. J. Mol. Evol. 17:94–102.

[12] López-Villaseñor, I., José, M.V., Sánchez, J. (2004) Three-base periodicity patterns and self-similarity in whole bacterial chromosomes. Biochem. Biophys. Res. Commun. 325(2):467–478.

[13] Shah, K., Krishnamachari, A. (2012) On the origin of three base periodicity in genomes. BioSystems 107:142–144.

[14] Michel, C.J., Thompson, J.D. (2020) Identification of a circular code periodicity in the bacterial ribosome: origin of codon periodicity in genes? RNA Biology 17(4):571–583.

[15] Eskesen, S.T., Eskesen, F.N., Kinghorn, B., et al. (2004) Periodicity of DNA in exons. BMC Mol. Biol. 5:12.

[16] Howe, E.D., Song, J.S. (2013) Categorical spectral analysis of periodicity in human and viral genomes. Nucl. Acid. Res. 41(3):1395–1405.

[17] Ding, Y., Tang, Y., Kwok, C.K., et al. (2014) In vivo genome-wide profiling of RNA secondary structure reveals novel regulatory features. Nature 505:696–700.

[18] Sánchez, J., López-Villaseñor, I. (2006) A simple model to explain three-base periodicity in coding DNA. FEBS Lett. 580:6413–6422.

[19] Michel, C.J., Sereni, J.-S. (2025) Genome galaxy identified by the circular code theory. Bulletin of Mathematical Biology 87:5.

[20] Wong, J.T. (1975) A coevolution theory of the genetic code. Proc. Natl. Acad. Sci. USA 72:1909–1912.

[21] Li, D.J. (2021) Formation of the codon degeneracy during interdependent development between metabolism and replication. Genes 12(12):2023.

[22] Li, D.J. (2022) Distributional features of triplet codons in genomes underlie the diversification of life. BioSystems 217: 104681.

[23] Harrison, P.W., Lopez, R., Rahman, N., Allen, S.G., Aslam, R., Buso, N., Cummins, C., Fathy, Y., Felix, E., Glont, M., Jayathilaka, S., Kadam, S., Kumar, M., Lauer, K.B., Malhotra, G., Mosaku, A., Edbali, O., Park, Y.M., Parton, A., Pearce, M., Estrada Pena, J.F., Rossetto, J., Russell, C., Selvakumar, S., Sitjà, X.P., Sokolov, A., Thorne, R., Ventouratou, M., Walter, P., Yordanova, G., Zadissa, A., Cochrane, G., Blomberg, N., Apweiler, R. (2021) The COVID-19 Data Portal: accelerating SARS-CoV-2 and COVID-19 research through rapid open access data sharing. Nucleic Acids Res. 49:W619–W623.

[24] Radivojac, P., Iakoucheva, L.M., Oldfield, C.J., Obradovic, Z., Uversky, V.N., Dunker, A.K. (2007) Intrinsic disorder and functional proteomics. Biophysical Journal 92:1439–1456.

[25] Dunker, A.K., Lawson, J.D., Brown, C.J., Williams, R.M., Romero, P., Oh, J.S., Oldfield, C.J., Campen, A.M., Ratliff, C.M., Hipps, K.W., Ausio, J., Nissen, M.S., Reeves, R., Kang, C., Kissinger, C.R., Bailey, R.W., Griswold, M.D., Chiu, W., Garner, E.C., Obradovic, Z. (2001) Intrinsically disordered protein. Journal of Molecular Graphics and Modelling 19:26–59.

[26] Romero, P., Obradovic, Z., Li, X., Garner, E.C., Brown, C.J., Dunker, A.K. (2001) Sequence complexity of disordered protein. Proteins: Structure, Function, and Genetics 42:38–48.

[27] Wootton, J.C., Federhen, S. (1993) Statistics of local complexity in amino acid sequences and sequence databases. Computers Chem. 17(2) 149–163.

[28] Lee, B., Jaberi-Lashkari, N., Calo, E. (2022) A unified view of low complexity regions (LCRs) across species. eLife 11:e77058.

[29] Zhou, P., Yang, X.L., Wang, X.G., Hu, B., Zhang, L., Zhang, W., Si, H., Zhu, Y., Li, B., Huang, C., Chen, H., Chen, J., Luo, Y., Guo, H., Jiang, R., Liu, M., Chen, Y., Shen, X., Wang, X., Zheng, X., Zhao, K., Chen, Q., Deng, F., Liu, L., Yan, B., Zhan, F., Wang, Y., Xiao, G., Shi, Z. (2020) A pneumonia outbreak associated with a new coronavirus of probable bat origin. Nature 579:270–273.

[30] Gao, J., Tang, Y.-D., Hu, W., Zheng, C. (2022) When poly(A) binding proteins meet viral infections, including SARS-CoV-2. J. Virology 97(7):e00136–22.

[31] Maździarz, M.A., Krawczyk, K., Lepiarczyk, E., Paukszto, Ł., Makowczenko, K.G., Moczulska, B., Iwanowicz, P., Kocbach, P., Nosek, K., Sawicki, J., Gromadziński, L., Majewska, M. (2025) Poly(A) tail dynamics, non-adenine incorporation and alternative polyadenylation shape the host transcriptome in COVID-19 pathogenesis. Scientific Reports 15:37986.

[32] Pizzarello, S. (2009) Meteorites and the chemistry that preceded life’s origin. in Wong, J. T., Lazcano, A., Eds., Prebiotic evolution and astrobiology. Landes Bioscience, Austin.

