## Supplementary methods, code, figures for "Diversifications of both the three domains of life and SARS-CoV-2 possibly driven by biases between amino acid biosynthetic families"

Dirson Jian Li

#### Diversification of the three domains of life based on TBP Family Bias in genomes

##### Construction of GenEvoSp based on TBP Family Bias in genomes

**TBP as a common feature and species-specific features** Three-base periodicity (TBP) in genomes is a common feature of bacteria, archaea, eukaryotes, and viruses (Supp. Fig. 4a), which was discovered shortly after the completion of whole-genome sequencing [1]. TBP refers to the preferential spacing of n-tuples, such as trinucleotides, by distances of multiples of three bases [1]. To illustrate, consider the genome sequence $b_1 b_2 \cdots b_k b_{k+1} \cdots b_s$: if the n-tuple $b_{k+1} b_{k+2} \cdots b_{k+n}$ is identical to $b_{m+k+1} b_{m+k+2} \cdots b_{m+k+n}$, the probability that $m$ is a multiple of three tends to be greater than the probability that it is not. A concrete example can be observed in the SARS-CoV-2 genome: if the trinucleotide $b_{k+1} b_{k+2} b_{k+3}$=$b_{m+k+1} b_{m+k+2} b_{m+k+3}$=$CGA$ (or any other codon), the probability that $m = 3j$ tends to be greater than that for $m = 3j \pm 1$, where $j$ is a natural number. This periodicity is significantly more prominent in coding regions than in non-coding regions, and the property has been applied to gene-finding algorithms [2] [3] [4] [5]. In addition to being a common feature of all life forms, TBP also exhibits species-specific features, which means that the preferential spacing of n-tuples, especially trinucleotides, differs across species. For a given species, in general, local TBP features in sufficiently long genomic regions are approximately similar to the global feature in its whole genome (Supp. Fig. 4dC). TBP is species-specific and stores abundant evolutionary information in the complicated fluctuations, which serves as a powerful tool for investigating the diversification of life [6].

**Definitions of $ifc$ based on the genome of a species** Interval frequency of codon ($ifc$) is a quantitative method to describe the species-specific TBP in the genome of a species, which is defined as the frequency of the intervals between the respective 64 trinucleotides (64 codons). Although trinucleotide and codon are two different concepts in the modern sense, their differences became ambiguous during the prebiotic evolution. It is of extreme value to classify trinucleotides by referencing the classification of codons, given that there is a close relationship between the distribution of trinucleotides in a genome and the distribution of codons in coding sequences. In order to emphasize this classification attribute, the term “codon” is often used instead of “trinucleotide” in the following text.

The genome sequence for a species $sp$ is denoted as

$$genome_{sp} = b_1 b_2 \cdots b_j b_{j+1} \cdots b_{gs},$$

where $gs$ is the genome size. If there are many chromosomes for a species, they can be concatenated together. All the positions of respective codons $cod = B_1 B_2 B_3$ in $genome_{sp}$ have been found at $pos_k(cod, sp)$, such that $b_{pos_k} b_{pos_k+1} b_{pos_k+2} = B_1 B_2 B_3$, where $k = 1, 2, \cdots, n_{pos}$. All the intervals between adjacent positions are $int_k = pos_{k+1} - pos_k$, where $k = 1, 2, \cdots, n_{pos} - 1$. The interval frequency is denoted as $ifc_0(int, cod, sp) = n$, which means that there are totally $n$ times for adjacent identical $codon$ with $int$-base spacing, namely

$$\cdots B_1 B_2 B_3 b_{j+4} b_{j+5} \cdots b_{j+int} B_1 B_2 B_3 \cdots ,$$

in $genome_{sp}$, where $int = 1, 2, \cdots, co$. The cutoff $co = 1000$ in the calculation is large enough that any additional errors can be ignored. Normalization is applied to $ifc_0$ so as to rule out the influence of genome size. Hence interval frequency of codon ($ifc$) is defined as follows:

$$ifc(int, cod, sp) = \frac{ifc_0(int, cod, sp)}{\sqrt{\sum_{for\ 64\ codons} \sum_{int=1}^{co} (ifc_0(int, cod, sp) \cdot ifc_0(int, cod, sp))}}.$$

Mean interval frequency of codon, denoted by $IFC$, is defined as the mean of $ifc$:

$$IFC(int, sp) = \frac{1}{64} \sum_{for\ 64\ codons} ifc(int, cod, sp).$$

Logarithm of $ifc$ is defined as $log\_ifc = log(ifc(int, cod, sp))$. Power spectrum of $ifc$, denoted by $ps\_ifc$, is the square of the modulus of its discrete Fourier transformation:

$$ps\_ifc(v, cod, sp) = \mid fft(ifc(int, cod, sp)) \mid^2,$$

where $v = 1,\ 2, \cdots, co$. Fluctuation of $ifc$ is defined as:

$$F\_ifc(sp) = \sum_{for\ 64\ codons} \sum_{int=1}^{co-1} |ifc(int+1, cod, sp) - ifc(int, cod, sp)|.$$

The interval frequency of codon for a taxon is defined as follows

$$\overline{ifc}(int, cod, taxon) = \frac{1}{n_{sp}} \sum_{sp \in taxon} ifc(int, cod, sp),$$

where $n_{sp}$ is the number of species in the taxon.

In addition, the above definitions can be generalized to $n$-order ones by calculating the distributions of the $4^n$ n-tuple $cod^n = B_1 B_2 \cdots B_n$ rather than the $4^3$ $cod$ in $genome_{sp}$. $n$-order $ifc$, denoted by $ifc^n(int, cod^n, sp)$, is the normalized interval frequencies for the respective n-tuples, and its mean is

$$IFC^n(int, sp) = \frac{1}{4^n} \sum_{for\ 4^n\ cod^n s} ifc^n(int, cod^n, sp).$$

**Additivity of $ifc$ and spatiotemporal rescaling** Note that $ifc$ satisfies additivity during sequence duplications. In general, $ifc$ remains unchanged after whole-genome duplication

$$b_1b_2\cdots b_jb_{j+1}\cdots b_{gs}b_1b_2\cdots b_jb_{j+1}\cdots b_{gs},$$

owing to the normalisation in the definition, and approximately unchanged after partial duplication

$$b_1b_2\cdots b_jb_{j+1}\cdots b_{j+k}b_jb_{j+1}\cdots b_{j+k}\cdots b_{gs},$$

owing to the similar interval frequencies between the sufficiently long partial sequence $b_jb_{j+1}\cdots b_{j+k}$ and the whole sequence $b_1b_2\cdots b_jb_{j+1}\cdots b_{j+k}\cdots b_{gs}$ for real species. Considering the approximate equality of $ifc$ between a pair of complementary codons, such as GCA and TGC:

$$ifc(int, B_1B_2B_3, sp) \approx ifc(int, \overline{B}_3\overline{B}_2\overline{B}_1, sp)$$

for real species, $ifc$ remains approximately unchanged after reverse partial duplication

$$b_1b_2\cdots b_jb_{j+1}\cdots b_{j+k}\cdots \bar{b}_{j+k}\cdots \bar{b}_{j+1}\bar{b}_j\cdots b_{gs}.$$

Étienne Geoffroy Saint-Hilaire explicitly stated that major evolutionary transformations relate to the spatiotemporal rescaling of embryogenesis [7] [8] [9] [10]. The diversification of life can be regarded as a spatiotemporal rescaling of the prebiotic evolution, especially the staged evolution of the genetic code. The additivity of $ifc$ in sequence duplications underlies the mechanism of the rescaling at the sequence level.

**Definitions of $CW$, and $F$-$F$ bias based on the genome of a species** Codon weight $CW$ is defined as:

$$CW(fam, sp) = \frac{1}{n_{cod}(fam)} \sum_{cod \in F_{fam}} cw(cod, sp),$$

where

$$cw(cod, sp) = \sum_{int=1}^{co} ifc(int, cod, sp) \cdot ifc(int, cod, sp),$$

considering the normalization of $ifc$: $\sum_{for\ 64\ codons} cw(cod, sp) = 1$. $CW$ for the amino acid biosynthetic families (Tab. 1) are as follows:

$$CW(Ala, sp) = \frac{1}{n_{cod}(Ala)} \sum_{cod \in F_{Ala}} cw(cod, sp),$$

$$CW(Ser, sp) = \frac{1}{n_{cod}(Ser)} \sum_{cod \in F_{Ser}} cw(cod, sp),$$

$$CW(Glu, sp) = \frac{1}{n_{cod}(Glu)} \sum_{cod \in F_{Glu}} cw(cod, sp),$$

$$CW(Asp, sp) = \frac{1}{n_{cod}(Asp)} \sum_{cod \in F_{Asp}} cw(cod, sp),$$

$$CW(Phe, sp) = \frac{1}{n_{cod}(Phe)} \sum_{cod \in F_{Phe}} cw(cod, sp).$$

There is a close relationship between TBP and the triplet genetic code. Since amino acid biosynthetic families played significant roles in the evolution of the genetic code, it is of value to study TBP in terms of amino acid biosynthetic families. As mentioned above, the species-specific TBP can be described quantitatively by $ifc$. Moreover, TBP Family Bias can be described quantitatively by $F$-$F$ bias.

$F$-$F$ bias is a quantitative method to describe TBP Family Bias in the genome of a species, which is defined as the difference between a pair of $CW$ (Tab. 1) as follows:

$$F_{fam_{early}} - F_{fam_{late}}\, bias(sp) = CW(fam_{late}, sp) - CW(fam_{early}, sp).$$

Specifically, three notable $F$-$F$ biases in classification of life are listed below:

$$F_{Ala} - F_{Ser}\, bias(sp) = CW(Ser, sp) - CW(Ala, sp),$$

$$F_{Glu} - F_{Asp}\, bias(sp) = CW(Asp, sp) - CW(Glu, sp),$$

$$F_{Ser} - F_{Phe}\, bias(sp) = CW(Phe, sp) - CW(Ser, sp).$$

**Definitions of GenEvoSp based on genomes** The genome evolution space (GenEvoSp) is spanned by the three $F$-$F$ biases: $F_{Ala} - F_{Ser}$ bias, $F_{Glu} - F_{Asp}$ bias, and $F_{Ser} - F_{Phe}$ bias. Each species corresponds to a fixed point in GenEvoSp:

$$\text{genome point: } (F_{Ala} - F_{Ser}\, bias(sp),\ F_{Glu} - F_{Asp}\, bias(sp),\ F_{Ser} - F_{Phe}\, bias(sp)).$$

Distribution of a taxon in GenEvoSp is composed of all the points of species in this taxon:

$$\text{genome points: } (F_{Ala} - F_{Ser}\, bias(sp),\ F_{Glu} - F_{Asp}\, bias(sp),\ F_{Ser} - F_{Phe}\, bias(sp)),\ sp \in taxon.$$

The data processing methods described above can be applied to both real and simulated genomes.

#### Diversification of the three domains of life in GenEvoSp

The global distribution of all life forms in GenEvoSp approximates a tetrahedron. Its four vertices are occupied by bacteria with high GC content, bacteria with low GC content, archaea, and eukaryotes, whereas viruses are situated inside the tetrahedral volume (Fig. 1a, 1b; Supp. Fig. 1aB, 1bA). Bacteria with high GC content and archaea lie along one tetrahedral edge parallel to the $F_{Ala} - F_{Ser}$ bias axis, with archaea positioned on the positive side of this axis (Supp. Fig. 1aC). Originating from about the midpoint of this edge, eukaryotes extend toward the positive direction of the $F_{Glu} - F_{Asp}$ bias axis, while bacteria with low GC content spread along the positive direction of the $F_{Ser} - F_{Phe}$ bias axis (Supp. Fig. 1aC).

Species distribution density varies substantially throughout the 3-dimensional GenEvoSp, forming the observed plaid distribution (Fig. 1b; Supp. Fig. 1bB, 1bC, 1bD).

The numbers of genomes are listed as follows in studying the distribution of life in GenEvoSp:

- Bacteria: 1040
- Archaea: 147
- Eukaryotes: 94
- Viruses: 10225

#### Simulation of diversification of the three domains of life in GenEvoSp

TBP can be generated by concatenating the triplet codons selected from proper subsets of all codons with different weights. The 64 codons are divided into six groups: $F_{Ala}$, $F_{Ser}$, $F_{Glu}$, $F_{Asp}$, $F_{Phe}$, $F_{others}$. $F_{others}$ consists of the codons encoding histidine and the three termination codons. Denote the six groups by $F_{fam_i}$, and denote the weight for $F_{fam_i}$ as $w_i$, $i = 1, 2, \cdots, 6$. The number of codons in $F_{fam_i}$ is $n_{cod_i}$, and $n_{cod_i}$=14, 12, 16, 12, 5, 5. (Tab. 1). Normal selection refers to selecting codons from $F_{fam_i}$ by probability $\frac{n_{cod_i}}{64}$. Biased selection refers to selecting codons from $F_{fam_i}$ by probability $\frac{n_{cod_i} \cdot w_i}{\sum_{i=1}^{6}(n_{cod_i} \cdot w_i)}$. In the simulation, the weights for Virus, Bacteria (high GC%), Bacteria (low GC%), Archaea, Eukarya are $w_i(dom)$, where $dom = 1, 2, 3, 4, 5, 6$, respectively (Supp. Tab. 1). Biased selection is such that codons are selected from $F_{fam_i}$ for different domains with different probabilities. For example, more codons are selected from $F_{Ala}$ for simulated bacteria with high GC content; more from $F_{Ser}$ for simulated archaea; more from $F_{Asp}$ for simulated eukaryotes; and more from $F_{Phe}$ for simulated bacteria with low GC content (Supp. Tab. 1) (Fig. 1d).

Supp. Tab. 1: Weights for amino acid biosynthetic families in simulation of diversification of life

| **domain** | $F_{Ala}$ | $F_{Ser}$ | $F_{Glu}$ | $F_{Asp}$ | $F_{Phe}$ | $F_{others}$ |
|---|---|---|---|---|---|---|
| viruses ($F_{Glu}$-rich, disorder driving aa) | 1 | 1 | 4 | 1 | 1 | 0.2 |
| viruses (low bias) | 1 | 2 | 0.2 | 3 | 6 | 0.2 |
| bacteria (high GC%) | 4 | 1 | 0.1 | 1 | 1 | 0.2 |
| bacteria (low GC%) | 1 | 1 | 0.1 | 1 | 6 | 0.2 |
| archaea | 1 | 4 | 0.1 | 1 | 1 | 0.2 |
| eukaryotes | 1 | 1 | 0.1 | 4 | 1 | 0.2 |

The distribution pattern of certain taxa can be observed and simulated in GenEvoSp. A variety of simulated genomes have been generated by evenly changing parameters to explain the separation of the three domains of life and viruses in GenEvoSp. The simulated global distribution area of life is approximately a tetrahedron, whose four vertices correspond to the four amino acid biosynthetic families: $F_{Ala}$, $F_{Phe}$, $F_{Ser}$ and $F_{Asp}$, respectively (Fig. 1d; Supp. Fig. 4dD). Simulated $F_{Ala}$ rich genomes are distributed outside the triangular surface $F_{Ala}F_{Phe}F_{Ser}$ (Supp. Fig. 4dD), where some ssDNA viruses can be found (Fig. 1d). $F_{Ala}$ rich genomes encode additional disorder driving amino acids, which are not conducive to protein folding.

Cellular life forms are distributed around the four vertices of the tetrahedron, while viruses are distributed inside the tetrahedron. There is greater TBP Family Bias at the corners, resulting in greater translation efficiency. This can be illustrated by the principle “Gold corners, silver sides, grass Belly”. In the game of Go, Go stones are easier to survive at the four gold corners than in the grass belly on the Go board.

Both observation and simulation of the tetrahedron in GenEvoSp indicate that the driving force to separate archaea and bacteria came from increasing weight of $F_{Ser}$ and decreasing weight of $F_{Ala}$, while the driving force to separate eukaryotes and archaea comes from increasing weight of $F_{Asp}$ and decreasing weight of $F_{Glu}$ (Fig. 1a; Supp. Fig. 1aC).

There is a negative shift of the tetrahedron for real life, which means that the edge $F_{Ala}F_{Ser}$ of the tetrahedron moves along the negative direction (Supp. Fig. 1aC). If only one strand of DNA is calculated in the simulation, there is no shift of the tetrahedron, i.e., zero $F_{Glu}$-$F_{Asp}$ bias for the edge $F_{Ala}F_{Ser}$ of the tetrahedron (Supp. Fig. 1dA, 1aC). If both strands of DNA are calculated in the simulation, the $F_{Glu}$-$F_{Asp}$ bias of the edge $F_{Ala}F_{Ser}$ becomes minus (Supp. Fig. 1dB), which agrees with the observation of real life (Fig. 1a, 1d; Supp. Fig. 1dD).

A clear plaid pattern is visible on the tetrahedron obtained in the simulation. The distribution density of simulated genomes exhibits distinct high-low fluctuations, which arise from the degeneracy of the genetic code grouped by amino acid biosynthetic families (Fig. 1d; Supp. Fig. 1dC, 1dD). Random and even sequence variations lead to the step changes in GenEvoSp and ProEvoSp, owing to the grouping of $ifc(int, cod, sp)$ by amino acid biosynthetic families in terms of codons.

Generally, only the minority of changes of codons are able to bring about shifts between biosynthetic families, while the remaining majority of changes keep the codons within their original biosynthetic families. For example, changes such as UAC to UCA, UUC to UCU and UGG to GGU can bring about shift from $F_{Phe}$ to $F_{Ser}$. For another example, changes such as CAG to GAC, CAA to AAC and CCA to ACC can bring about shift from $F_{Glu}$ to $F_{Asp}$.

The three-base periodicity is crucial in the simulation. If we abandon such a common feature of life (Supp. Fig. 1cA), the global distribution of genomes generated randomly by evenly varying all possible parameters forms a trident star (Supp. Fig. 1cC). The trident star corresponding to randomly simulated genome sequences differs substantially and cuts across the tetrahedron corresponding to three-base periodic genomes (Fig. 1c; Supp. Fig. 1cD). Especially the four vertices of the tetrahedron lie far from the curved surface of the trident star (Fig. 1c; Supp. Fig. 1cD). TBP in genomes ensures that real genomes are kept far away from randomness.

### Diversification of SARS-CoV-2 based on TBP Family Bias in genomes

#### Diversification of SARS-CoV-2 in GenEvoSp

The diversification and evolutionary trend of SARS-CoV-2 can be studied by the distributions of SARS-CoV-2 genomes with different collection regions, collection times, and types, in GenEvoSp.

In this study, the term “Kite” is used to represent the kite-like distribution of SARS-CoV-2 genomes in GenEvoSp for a certain collection region, certain collection time, or certain type (Fig. 2; Supp. Fig. 2a). For example, America Kite represents the distribution of SARS-CoV-2 genomes from America collected in 2020 (Supp. Fig. 2aA). Each Kite exhibits a single axis of symmetry, and the symmetry axes of all identified Kites are mutually parallel (Fig. 2a, 2c; Supp. Fig. 2aC, 2aD). Ordered subclusters align along this symmetry axis, with the poly(A) tail length (the number of consecutive adenines) of SARS-CoV-2 genomes differing by precisely one nucleotide between adjacent subclusters (Supplementary Fig. 2aA, 2bD). A complete Kite (Supp. Fig. 2bC) may consist of three structural components: long-tail array (Supp. Fig. 2bB), middle-tail array (Supp. Fig. 2bA), and short-tail array (Fig. 2b). Within each array, more subclusters extend outward in both flanking directions. The periodic density of SARS-CoV-2 genome distribution within a Kite exhibits a plaid pattern at the local scale and a kite-shaped distribution profile at the global scale (Supp. Fig. 2bC). The distribution density of subclusters is continuous in the ventral-dorsal direction. Generally, the short-tail array is bigger than long-tail array (Supp. Fig. 2bC). The middle-tail array is smallest in America Kite and often absent in other Kites (Supp. Fig. 2aC). The long-tail array may vanish in certain Kite sometimes (Supp. Fig. 2aC).

Many different Kites have been obtained. The early China Kite is situated on one flank of America Kite (Supp. Fig. 2cC), whereas Omicron Kite and farther KP2 Kite on the other flank (Supp. Fig. 2cD).

SARS-CoV-2 evolves synchronously in both short-tail array and long-tail array (Supp. Fig. 2cA, 2cB). Strikingly, the boundary between different SARS-CoV-2 variants aligns precisely with the boundary between different subclusters in GenEvoSp.

When marking collection time by day in the America Kite, SARS-CoV-2 evolution starts from one wing of the middle array, then expands to the entire Kite, including the long-tail array and short-tail array, as well as the other wing of the middle-tail array (Supp. Fig. 2cA). When additionally marking genome sizes, an oscillation in genome size has been observed. The genome size initially decreases, then increases rapidly, and ultimately stabilizes at a nearly constant size (Supp. Fig. 2cB).

The numbers of genomes are listed as follows in studying the distribution of SARS-CoV-2 in GenEvoSp:

- America (America 2020): 17706, Australian: 9894, China: 2785, Iceland: 21268, Italy: 1422, New Zealand: 16636, Spain: 1295;
- Beta: 1173, Delta: 1094, Alpha: 979, Gamma: 926, Epsilon: 1134, BA1: 1379, XBB: 985, BA3: 435, BA4: 1664, BA5: 1774, Zeta: 1550, Eta: 705, Lambda: 719, Mu: 2105, Iota: 1777, KP2: 1326;

- Omicron randomly selected from all: 20000, Omicron 2024: 21710, SARS-CoV-2 NCBI: 12449;
- China 2019: 19, China Jan 2020: 105, China Feb 2020: 327, China Mar 2020: 116, China Apr-Dec: 104;
- SARS 2003: 69, SARS 2004: 44, SARS 2005: 23, Bat CoV: 35.

##### Evolutionary trend of SARS-CoV-2 in GenEvoSp

The evolutionary trend can be studied by observing SARS-CoV Kite, bat coronavirus Kite, and early China Kite, America Kite, Omicron Kite in a sequential order in GenEvoSp (Fig. 2c). The trend roughly points towards the positive direction of $F_{Ser} - F_{Phe}$ bias. Among all bat coronaviruses analyzed, RaTG13 is the closest to the early SARS-CoV-2 Kites in GenEvoSp (Fig. 2c), consistent with prior reports [11]. According to the drift of SARS-CoV Kites from 2003 to 2005, the evolution direction of SARS-CoV-2 is the same as the evolution direction of SARS-CoV, both of which keep away from the position of human beings in GenEvoSp (Fig. 2c). Within the inner region of the tetrahedron in GenEvoSp, both SARS-CoV and SARS-CoV-2 Kites are near to the vertex for eukaryotes (Supp. Fig. 1aA).

##### Simulations of diversification and evolutionary trend of SARS-CoV-2 in GenEvoSp

The Kite-shaped distribution patterns of SARS-CoV-2 have been simulated (Fig. 2d; Supp. Fig. 2d). Simulations began with the generation of random genomes with GC content of about 40%. Subclusters along the symmetry axis in a Kite were simulated by introducing poly(A) tracts with varying lengths of 0, 1, 2, $\cdots$, 40 adenines. The flank direction distributions of short-tail array, middle-tail array, and long-tail array in a Kite were then simulated by varying AT content in genomes with poly(A) lengths about 0 to 2 adenines, 13 adenines, 31 and 33 adenines, respectively. The plaid Kite pattern was reproduced via substitution of only a small fraction of bases. The shape and orientation of the simulated Kite approximately agree with the observation based on SARS-CoV-2 genomes (Supp. Fig. 2dA, Fig. 2a).

Simulations by changing base compositions result in Kite drifts over sufficiently long time (Supp. Fig. 2dB). The overall evolutionary trend of SARS-CoV-2 is roughly along the positive direction of $F_{Ser} - F_{Phe}$ bias axis (Supp. Fig. 2dC, 2dD).

#### Diversification of the three domains of life based on IFAA Family Bias in proteomes

##### Construction of ProEvoSp based on IFAA Family Bias in proteomes

There is a direct relationship between the interval between codons in a genome and the interval between the corresponding amino acids in the translated proteome. Hence, ProEvoSp can be defined similarly as the above genome evolution space.

**Definitions of $if\tilde{c}$ based on the proteome of a species** Interval frequency of amino acids ($if\tilde{c}$) is a quantitative method to describe the species-specific proteome features, which is defined as the frequency of the intervals between the respective 20 amino acids. The following interval frequencies will be calculated for each protein in the proteome of a certain species and then summed up for each of the 20 amino acids, respectively.

The proteome sequence for a species $sp$ is denoted as

$$proteome_{sp} = \{protein_1, protein_2, \cdots, protein_i, protein_{i+1} \cdots, protein_{n_p}\},$$

where $n_p$ is the number of proteins in this proteome. A protein sequence is

$$protein_i = a_1 a_2 \cdots a_j a_{j+1} \cdots a_l,$$

where $l$ is the number of amino acid residues in this protein. All the positions of respective amino acid residues $aa = A$ in $protein_i$ have been found at $pos_k(i, aa, sp)$, such that $a_{pos_k} = A$, where $k = 1, 2, \cdots, n_{pos}$. All the intervals between adjacent positions are $int_k = pos_{k+1} - pos_k$, where $k = 1, 2, \cdots, n_{pos} - 1$. The interval frequency is denoted as $ifaa_0(i, int, aa, sp) = n$, which means that there are totally $n$ times for adjacent identical $aa$ with $int$-amino acid residue spacing, namely

$$\cdots A a_{j+2} a_{j+3} \cdots a_{j+int} A \cdots,$$

in $protein_i$, where $int = 1, 2, \cdots, co$. The cutoff $co = 1000$ in the calculation is large enough that any additional errors can be ignored. Sum up all protein results $ifaa_0$ to obtain the proteome results $ifaa$:

$$ifaa(int, aa, sp) = \sum_{i=1}^{n_p} ifaa_0(i, int, aa, sp).$$

Normalization is applied to $ifaa$ so as to rule out the influence of protome size. Hence interval frequency of amino acid (IFAA, or $if\tilde{c}$) is defined as follows:

$$if\tilde{c}(int, aa, sp) = \frac{ifaa(int, aa, sp)}{\sqrt{\sum_{aa \in 20 \ amino \ acids} \sum_{int=1}^{co} (ifaa(int, aa, sp) \cdot ifaa(int, aa, sp))}}.$$

Mean interval frequency of amino acid, denoted by $IF\tilde{C}$, is defined as the mean of $if\tilde{c}$:

$$IF\tilde{C}(int, sp) = \frac{1}{20} \sum_{aa \in 20 \ amino \ acids} if\tilde{c}(int, aa, sp).$$

Logarithm of $if\tilde{c}$ is defined as $log\_if\tilde{c} = log(if\tilde{c}(int, aa, sp))$. Power spectrum of $if\tilde{c}$, denoted by $ps\_if\tilde{c}$, is the square of the modulus of its discrete Fourier transformation:

$$ps\_if\tilde{c}(e, aa, sp) = |\ fft(if\tilde{c}(int, aa, sp))\ |^2,$$

where $e = 1,\ 2, \cdots, co$. The interval frequency of amino acid for a taxon is defined as follows

$$\overline{if\tilde{c}}(int, aa, taxon) = \frac{1}{n_{sp}} \sum_{sp \in taxon} if\tilde{c}(int, aa, sp),$$

where $n_{sp}$ is the number of species in the taxon, and its logarithm is $log(\overline{if\tilde{c}}(int, aa, taxon))$ (Supp. Fig. 3c). $if\tilde{c}$ peak is defined as:

$$if\tilde{c}\, peak(sp) = \sum_{aa \in 20\ amino\ acids} \sum_{int=2}^{co-1} Pk(if\tilde{c}(int, aa, sp)),$$

where $Pk(if\tilde{c}(int, aa, sp)) = 1$ if $if\tilde{c}(int, aa, sp)$ is greater than both $if\tilde{c}(int+1, aa, sp)$ and $if\tilde{c}(int-1, aa, sp)$, and $Pk(if\tilde{c}(int, aa, sp)) = 0$ otherwise. $if\tilde{c}\, peak(sp)$ describes the magnitude of $if\tilde{c}$ fluctuations for a species. The correlation coefficient matrix between pairs of species $sp_i$ and $sp_j$ in a group of $n_{sp}$ species is: $cc(i, j) = corrcoef(if\tilde{c}(int, aa, sp_i), if\tilde{c}(int, aa, sp_j))$, and the mean correlation coefficient of $if\tilde{c}$ for species $sp_i$ is defined as $\overline{cc}(sp_i) = \frac{1}{n_{sp}} \sum_{j=1}^{n_{sp}} cc(i, j)$, and logarithm of mean correlation is $log(\overline{cc}(sp_i))$. $\overline{cc}$ describes the mean correlation coefficient between a species and others. The diversification of life can be studied in the plane spanned by $log(\overline{cc})$ and $if\tilde{c}\, peak$ (Supp. Fig. 3aD).

The positions of synonymous codons for a certain amino acid can be denoted in the genome, then we calculate the intervals between these positions and obtain the distribution of intervals between synonymous codons in the genome, which bridges the genomic $ifc$ and the proteomic $if\tilde{c}$. Clustering of closely related species in GenEvoSp or ProEvoSp result from their clustering in the high-dimensional space based on $ifc$ or $if\tilde{c}$ before dimension reduction.

**Definitions of $\tilde{C}W$, and $\tilde{F}$-$\tilde{F}$ bias based on the proteome of a species** Amino acid weight $\tilde{C}W$ is defined as:

$$\tilde{C}W(fam, sp) = \frac{1}{n_{aa}(fam)} \sum_{aa \in \tilde{F}_{fam}} \tilde{c}w(aa, sp),$$

where

$$\tilde{c}w(aa, sp) = \sum_{int=1}^{co} if\tilde{c}(int, aa, sp) \cdot if\tilde{c}(int, aa, sp),$$

considering the normalization of $if\tilde{c}$: $\sum_{aa \in 20\ amino\ acids} \tilde{c}w(aa, sp) = 1$. $\tilde{C}W$ for the amino acid biosynthetic families (Tab. 1) are as follows:

$$\tilde{C}W(Ala, sp) = \frac{1}{n_{aa}(Ala)} \sum_{aa \in \tilde{F}_{Ala}} \tilde{c}w(aa, sp),$$

$$\tilde{C}W(Ser, sp) = \frac{1}{n_{aa}(Ser)} \sum_{aa \in \tilde{F}_{Ser}} \tilde{c}w(aa, sp),$$

$$\tilde{C}W(Glu, sp) = \frac{1}{n_{aa}(Glu)} \sum_{aa \in \tilde{F}_{Glu}} \tilde{c}w(aa, sp),$$

$$\tilde{CW}(Asp, sp) = \frac{1}{n_{aa}(Asp)} \sum_{aa \in \tilde{F}_{Asp}} \tilde{cw}(aa, sp),$$

$$\tilde{CW}(Phe, sp) = \frac{1}{n_{aa}(Phe)} \sum_{aa \in \tilde{F}_{Phe}} \tilde{cw}(aa, sp).$$

IFAA Family Bias can be described quantitatively by $\tilde{F}$-$\tilde{F}$ bias.

$\tilde{F}$-$\tilde{F}$ bias is a quantitative method to describe IFAA Family Bias in the proteome of a species, which is defined as the difference between a pair of $\tilde{CW}$ (Tab. 1) as follows:

$$\tilde{F}_{fam_{early}} - \tilde{F}_{fam_{late}}\, bias(sp) = \tilde{CW}(fam_{late}, sp) - \tilde{CW}(fam_{early}, sp).$$

Specifically, three notable $\tilde{F}$-$\tilde{F}$ biases in classification of life are listed below:

$$\tilde{F}_{Ala} - \tilde{F}_{Ser}\, bias(sp) = \tilde{CW}(Ser, sp) - \tilde{CW}(Ala, sp),$$

$$\tilde{F}_{Glu} - \tilde{F}_{Asp}\, bias(sp) = \tilde{CW}(Asp, sp) - \tilde{CW}(Glu, sp),$$

$$\tilde{F}_{Ser} - \tilde{F}_{Phe}\, bias(sp) = \tilde{CW}(Phe, sp) - \tilde{CW}(Ser, sp).$$

**Definitions of ProEvoSp based on proteome** The proteome evolution space (ProEvoSp) is spanned by the three $\tilde{F}$-$\tilde{F}$ biases: $\tilde{F}_{Ala} - \tilde{F}_{Ser}$ bias, $\tilde{F}_{Glu} - \tilde{F}_{Asp}$ bias, and $\tilde{F}_{Ser} - \tilde{F}_{Phe}$ bias. Each species corresponds to a fixed point in ProEvoSp:

$$\text{proteome point: } (\tilde{F}_{Ala} - \tilde{F}_{Ser}\, bias(sp),\ \tilde{F}_{Glu} - \tilde{F}_{Asp}\, bias(sp),\ \tilde{F}_{Ser} - \tilde{F}_{Phe}\, bias(sp)).$$

Distribution of a taxon in ProEvoSp is composed of all the points of species in this taxon:

$$\text{proteome points: } (\tilde{F}_{Ala} - \tilde{F}_{Ser}\, bias(sp),\ \tilde{F}_{Glu} - \tilde{F}_{Asp}\, bias(sp),\ \tilde{F}_{Ser} - \tilde{F}_{Phe}\, bias(sp)),\ sp \in taxon.$$

The data processing methods here can also be applied to both real and simulated proteomes.

#### Diversification of the three domains of life in ProEvoSp

Random sampling of species from all life forms recovers a global triangular distribution of life in the ProEvoSp space. Bacteria, archaea, and eukaryotes are distributed in the three angular regions of the triangle, respectively (Fig. 3a), while viruses are roughly inside the triangle (Fig. 3a). Specifically, clearly separated clusters are distinguishable between animals and plants (Supp. Fig. 3aB), as well as between invertebrates and vertebrates (Supp. Fig. 3aA). The evolutionary trend from invertebrates to vertebrates, and further specifically to mammals follows a sequential path pointing toward the eukaryote vertex (Supp. Fig. 3aC). Notably, both driving forces in GenEvoSp and ProEvoSp are due to the biases between amino acid biosynthetic families.

A two-dimensional distribution of life can be constructed using proteome size and number of peaks in $if\tilde{c}$ as the two coordinate axes (Fig. 3c). Along a continuous curve in this space, four successive paired taxonomic groups are arranged in sequence: DNA viruses and RNA viruses, bacteria and archaea, fungi and protozoa, plants and animals (Fig. 3c; Supp. Fig. 3c).

A proper subset of a full proteome corresponds to a candidate collection of translated proteins, or a candidate transcriptome. Denote $R_{pp}$ as the ratio of number of proteins in the proper subset and number of proteins in the proteome of a species. For A. thaliana and papaya in Supp. Fig. 3bA, $R_{pp}$ is set to $1\%$, $10\%$, and $83.3\%$, and for the species in Fig. 3b and Supp. Fig. 3bB, $R_{pp}$ is set to $10\%$. In ProEvoSp, the spatial distribution of a large number of randomly sampled proteome proper subsets is aligned with the distribution of the taxonomic group to which the source species belongs in direction (Fig. 3b; Supp. Fig. 3bA, 3bB).

Given a proteome of a certain species, the density of proteins distributed in ProEvoSp is uneven. For example, high density in discrete small subregions of the ProEvoSp space has been observed for the large number of proteins in the proteome of Bufo bufo (Supp. Fig. 3bC) or in the proteome of Homo sapiens (Supp. Fig. 3bD).

The numbers of proteomes are listed as follows in studying the distribution of life in ProEvoSp:

- Bacteria: 1477
- Archaea: 160
- Eukaryotes: 1144 (fungi: 545, plants: 134, protozoa: 94, invertebrates: 257; vertebrates excluding mammals (vertebrate other): 75, mammals: 39)
- Viruses: 329 (RNA viruses: 115, DNA viruses: 214)

#### Simulation of diversification of the three domains of life in ProEvoSp

Simulations help understand the compression from the tetrahedron in GenEvoSp (Fig. 1d) to the triangle in ProEvoSp (Fig. 3d). The simulated proteomes are obtained by translating the simulated genomes in section 1.3 according to the canonical genetic code. The distribution of the simulated proteomes in ProEvoSp is a regular tetrahedron (Supp. Fig. 3dA). The parameters are chosen randomly and evenly among all possible parameters. The plaid pattern in this tetrahedron is due to the degeneracy of the genetic code grouped by amino acid biosynthetic families (Supp. Fig. 3dA). The mechanism that compresses the tetrahedron into a triangle arises from the six reading frames. The genome is only taken as 1 reading frame in the previous translation. If considering the three (Supp. Fig. 3dB) or the six (Fig. 3d) reading frames of the genome in the translation process, namely

1 reading frame: $b_1b_2b_3\cdots b_{3l-2}b_{3l-1}b_{3l}$,

3 reading frames: $b_1b_2b_3\cdots b_{3l-2}b_{3l-1}b_{3l}b_2b_3b_4\cdots b_{3l-4}b_{3l-3}b_{3l-2}b_3b_4b_5\cdots b_{3l-3}b_{3l-2}b_{3l-1}$,

6 reading frames: $b_1b_2b_3\cdots b_{3l-2}b_{3l-1}b_{3l}b_2b_3\cdots b_{3l-2}b_3\cdots b_{3l-2}b_{3l-1}\bar{b}_{3l-1}\bar{b}_{3l-2}\cdots\bar{b}_3\bar{b}_{3l-2}\cdots\bar{b}_3\bar{b}_2\bar{b}_{3l}\bar{b}_{3l-1}\bar{b}_{3l-2}\cdots\bar{b}_3\bar{b}_2\bar{b}_1$,

simulated proteomes with 3 or 6 times the original length are obtained for each genome. Based on these reading-frame mixing simulated proteomes, the two vertices of bacteria with high and low GC content in the tetrahedron are compressed into one vertex in the triangle (Fig. 3d; Supp. Fig. 3d), and the global distribution of life becomes a triangle in ProEvoSp, which agrees with the observation for real life.

According to the above process from the tetrahedron in GenEvoSp to the triangle in ProEvoSp, the three domains of life correspond to the three vertices of the triangle, which provides an intuitive explanation for the number of domains of cellular life forms.

### Possible origin of the triplet nature of life and its role in translation efficiency

#### Simulation of initial TBP by the period averaging procedure

Evidence shows that periodicity appeared early in the prebiotic sequence evolution [12] [13] [14]. Frenkel et al. proposed that most of the genomic sequences evolved through everlasting acts of triplet expansions with subsequent accumulation of changes [15] [16] [17]. Ohno et al. suggested that the first set of base sequences translated at the beginning of life were oligonucleotide repeats beneficial for forming protein secondary structures [18] [19]. Shiba et al. found that periodicity played a pivotal role in the origin of many genes [20]. It is reasonable to suggest that the three-base periodicity formed earlier than the triplet nature of the genetic code.

A period averaging procedure is proposed here to provide a possible explanation for the initial emergence of TBP in nucleic acid sequences. This approach does not presuppose the initial specificity of the three-base unit. It is assumed that diverse periodic patterns may have existed in primordial sequences before the formation of both TBP and the triplet genetic code. The key question is whether the combined effects arising from the concatenation of various periodic sequences lead to the selective stabilization of TBP. It is shown that the combinatorial mixing of different periodic sequences tends to generate TBP as a statistically predominant outcome.

The $p$-base periodicity can be obtained by mixing $p$-tuples ($p = 2, 3, 4, 5, \cdots$), respectively. For a certain period $p$, there are $N_C = 4^p$ p-tuples in the set of $p$-base repeatable permutations:

$$C^p = \{b_j^1 b_j^2 \cdots b_j^p | b_j^i \in B\},\ j = 1, 2, \cdots, N_C,$$

where $B$ is the set of four nucleotide bases. The set $P^p$ denotes any proper subset of $C^p$, namely $P^p \subset C^p$, whose $N_P$ elements are selected randomly from the elements of $C^p$. Let $R_{Proper} = N_P/N_C$ and take it as a moderate value; excessive or insufficient values of $R_{Proper}$ can lead to the vanishing of $p$-base periodicity. The simulated $p$-base short periodic sequences are

$$s^p = b_1^1 b_1^2 \cdots b_1^p b_2^1 b_2^2 \cdots b_2^p \cdots\cdots b_k^1 b_k^2 \cdots b_k^p \cdots\cdots b_{[L/p]}^1 b_{[L/p]}^2 \cdots b_{[L/p]}^p,$$

where $[\cdot]$ is the rounding symbol so as to obtain the approximately same sequence length $L$ for different short periodic sequences $s^p$. For example, simulated 2-, 3-, 4-, and 5-base short periodic sequences, in which some $p$-tuples are not used, are as follows, respectively:

$$s^2 = b_1^1 b_1^2 b_2^1 b_2^2 \cdots\cdots b_k^1 b_k^2 \cdots b_{[L/2]}^1 b_{[L/2]}^2, \text{ where } b_k^1 b_k^2 \in P^2,$$
$$s^3 = b_1^1 b_1^2 b_1^3 b_2^1 b_2^2 b_2^3 \cdots\cdots b_k^1 b_k^2 b_k^3 \cdots\cdots b_{[L/3]}^1 b_{[L/3]}^2 b_{[L/3]}^3, \text{ where } b_k^1 b_k^2 b_k^3 \in P^3,$$
$$s^4 = b_1^1 b_1^2 b_1^3 b_1^4 b_2^1 b_2^2 b_2^3 b_2^4 \cdots\cdots b_k^1 b_k^2 b_k^3 b_k^4 \cdots\cdots b_{[L/4]}^1 b_{[L/4]}^2 b_{[L/4]}^3 b_{[L/4]}^4, \text{ where } b_k^1 b_k^2 b_k^3 b_k^4 \in P^4,$$
$$s^5 = b_1^1 b_1^2 \cdots b_1^5 b_2^1 b_2^2 \cdots b_2^5 \cdots\cdots b_k^1 b_k^2 \cdots b_k^5 \cdots\cdots b_{[L/5]}^1 b_{[L/5]}^2 \cdots b_{[L/5]}^5, \text{ where } b_k^1 b_k^2 \cdots b_k^5 \in P^5.$$

Let $P_j^p$ be a distinct randomly selected proper subsets: $P_j^p \subset C^p$. Short periodic sequences generated from $P_j^p$ are as follows:

$$s_j^p = b_1^1 b_1^2 \cdots b_1^p b_2^1 b_2^2 \cdots b_2^p \cdots\cdots b_k^1 b_k^2 \cdots b_k^p \cdots\cdots b_{[L/p]}^1 b_{[L/p]}^2 \cdots b_{[L/p]}^p, \text{ where } b_k^1 b_k^2 \cdots b_k^p \in P_j^p,$$

where $j = 1, 2, \cdots, H$, and $H$ is the same so as to obtain the approximately same sequence length $HL$ for different periods. The simulated $p$-base long periodic sequences are as follows:

$$S^p = s_1^p s_2^p \cdots s_H^p.$$

For example, simulated 2-, 3-, 4-, and 5-base long periodic sequences, in which all $p$-tuples are possibly used, are as follows:

$$S^2 = s_1^2 s_2^2 \cdots s_H^2,$$
$$S^3 = s_1^3 s_2^3 \cdots s_H^3,$$
$$S^4 = s_1^4 s_2^4 \cdots s_H^4,$$
$$S^5 = s_1^5 s_2^5 \cdots s_H^5.$$

The simulated genome sequence is

$$gm_K = S^2 S^3 \cdots S^p \cdots S^K,$$

For example:

$$gm_5 = S^2 S^3 S^4 S^5,$$
$$gm_6 = S^2 S^3 S^4 S^5 S^6,$$
$$gm_7 = S^2 S^3 S^4 S^5 S^6 S^7,$$
$$gm_8 = S^2 S^3 S^4 S^5 S^6 S^7 S^8,$$
$$gm_9 = S^2 S^3 S^4 S^5 S^6 S^7 S^8 S^9,$$
$$gm_{10} = S^2 S^3 S^4 S^5 S^6 S^7 S^8 S^9 S^{10},$$
$$\cdots\cdots.$$

For example, in case of $K = 5$, the genome sequence is (Fig. 4a, 4b, 4e):

$$gm_5 = s_1^2 s_2^2 \cdots s_H^2 s_1^3 s_2^3 \cdots s_H^3 s_1^4 s_2^4 \cdots s_H^4 s_1^5 s_2^5 \cdots s_H^5;$$

for another example, in case of $K = 10$, the genome sequence is (Fig. 4c, 4f):

$$gm_{10} = s_1^2 \cdots s_H^2 s_1^3 \cdots s_H^3 s_1^4 \cdots s_H^4 s_1^5 \cdots s_H^5 s_1^6 \cdots s_H^6 s_1^7 \cdots s_H^7 s_1^8 \cdots s_H^8 s_1^9 \cdots s_H^9 s_1^{10} \cdots s_H^{10}.$$

Such a simulated $if\tilde{c}$ is similar to the $if\tilde{c}$ of Gallus gallus (Fig. 4i). The order of $s_i^p$ segments within $gm$ is permutable. In the simulation, the genome sequence can be generated as:

$$gm = S^{p_1} S^{p_2} \cdots S^{p_i} \cdots ,$$

where $p_i$ are randomly selected from natural numbers.

TBP has been simulated by the period mixing procedure. The 3-base periodicity roughly becomes the average period in $gm_K$ and $gm$ by randomly concatenating these 2-base, 3-base, ... , 10-base, etc. periodic sequences. Considering the same length $HL$ of the periodic sequences across different periods, fewer $p$-tuples for larger periods lead to their weaker contributions to the average period. Hence the weights for large periods become negligible. Simulations show that the number 3 is the overall period obtained by mixing sequences with different periods.

Diversification of primordial genomes prior to the establishment of the genetic code can be simulated by base substitutions between amino acid biosynthetic families after generating numerous genomes with random parameters (Supp. Fig. 4bC; Tab. 1), and the diversification of primordial proteomes can be obtained by translating these diverse genomes (Supp. Fig. 4bD). For instance, a primordial genome can be generated by the period mixing procedure (Supp. Fig. 4bA), and its corresponding proteome can be obtained by translating this genome (Supp. Fig. 4bB). Diverse primordial genomes that occupy GenEvoSp positions corresponding to bacteria with high GC content, bacteria with low GC content, archaea, and eukaryotes, can be obtained, respectively, by randomly introducing certain base substitutions, namely, no specific substitutions for viruses, UCA-to-UAC, UCU-to-UUC and AU-to-GC substitutions for bacteria with high GC content, substitutions to increase AT content for bacteria with low GC content, UCA-to-UAC and UCU-to-UUC substitutions for archaea, and UAC-to-UCA, UUC-to-UCU, UC-to-CU substitutions for eukaryotes.

#### Simulation of TBP features based on the staged evolution of the genetic code

Besides the period averaging procedure, TBP can also be generated via the triplet genetic code. During the staged evolution of the triplet genetic code, TBP can be generated by concatenating the triplet codons from proper subsets of all the 64 codons (Fig. 4d, 4g). Such a simulated $if\tilde{c}$ is similar to $if\tilde{c}$ of bacteria and archaea, e.g., Haloarcula hispanica N601 (Fig. 4h). Denote all the 64 codons by $C_{codon}^3$, and let $P^3$ be a proper subset with $N_P$ elements of $C_{codon}^3$: $P^3 \subset C_{codon}^3$. Short three-base periodic sequences generated from $P_j^p$ are as follows:

$$s_j^3 = b_1^1 b_1^2 b_1^3 b_2^1 b_2^2 b_2^3 \cdots\cdots b_k^1 b_k^2 b_k^3 \cdots\cdots b_l^1 b_l^2 b_l^3, \text{ where } b_k^1 b_k^2 b_k^3 \in P_j^3,$$

where $j = 1, 2, \cdots , H$. Thus, the three-base periodic genome sequences can be obtained by concatenating triplet codons as follows:

$$gm_c = s_1^3 s_2^3 \cdots s_H^3.$$

Let $R_{Proper} = N_P/64$. $R_{Proper}$ should be moderate. The more the number of elements in the proper subsets, the less the amplitude of the fluctuations in the three-base periodic $ifc$ (Fig. 4d; Supp. Fig. 1cB). The $ifc$ amplitude of viruses is generally greater than that of bacteria or archaea. The amplitude of eukaryotes is generally less than that of bacteria or archaea (Supp. Fig. 4a). If there are many certain repeated sequences in a eukaryotic genome, there are many sharp peaks in $ifc$ that generally are not three-base periodic (Supp. Fig. 4aG, 4aH).

Evidence shows that 10 of the 20 amino acids belong to phase I, which existed before the emergence of life, and the other 10 amino acids belong to phase II (Tab. 1). The temporal order of the five biosynthetic families, $F_{Ala}$, $F_{Ser}$, $F_{Glu}$, $F_{Asp}$, and $F_{Phe}$ can be roughly inferred by the ratios of number of phase I amino acids to number of phase II amino acids, 3:0, 2:1, 3:2, 2:3, and 0:3, respectively (Tab. 1). All positive directions of the three coordinates in GenEvoSp, $F_{Glu} - F_{Asp}$ bias, $F_{Ala} - F_{Ser}$ bias, and $F_{Ser} - F_{Phe}$ bias, represent the time direction from early to late. The direction from bacteria to archaea in GenEvoSp is towards the positive direction of $F_{Ala} - F_{Ser}$ bias axis (Supp. Fig. 2aC), and the direction from archaea to eukaryotes in GenEvoSp is also towards the positive direction of $F_{Glu} - F_{Asp}$ bias axis (Supp. Fig. 2aC). Family GC content is defined as the average GC content of the codons in certain amino acid biosynthetic family (Tab. 1). The least family GC content is 0.267 for $F_{Phe}$ (Tab. 1). The positive direction of $F_{Ser} - F_{Phe}$ bias axis generally corresponds to the increase of GC content.

The simulated genomes can be divided by the dominant ratio of amino acids in one of the five biosynthetic families of amino acids. The elements of the proper subset $P_j^p$ can be selected with different weights from the five subsets of $C_{codon}^3$: $F_{Ala}$, $F_{Phe}$, $F_{Ser}$, $F_{Asp}$, and $F_{Glu}$. Five types of simulated genomes correspond to the above five subsets, respectively. The former four of them are distributed around the four vertices of the tetrahedron in the evolution space, as the global distribution shape of simulated genomes (Fig. 1d), while the last $F_{Glu}$ is just outside the tetrahedron (Supp. Fig. 4dD). Dominant biosynthetic families benefit the efficiency to find the corresponding tRNAs around ribosome in the translation process, such that the cellular life forms are distributed around the vertices of the tetrahedron. The inner region of the tetrahedron is occupied by the simulated viral genomes without obviously dominant biosynthetic families, which is not conducive to translation efficiency.

The period averaging procedure may account for the initial formation of TBP. Once the initial sequence was three-base periodic and hence kept away from randomness, the choice of the triplet codons is associated with the high efficiency of finding tRNA in the translation process (Supp. Fig. 4dC). Reciprocally, TBP can be obtained again in that uneven usage of triplet codons led to TBP in concatenation of codons from a proper subset of the 64 codons. During the staged evolution of the genetic code, the recruited codons constitute a proper subset of all codons. TBP benefits the efficiency of translation. There is a positive feedback process that applies to any k-base nature of life. The choice of "three" initiated the long standing triplet nature of life.

##### Role of the triplet nature of life in translation efficiency

Each dominant biosynthetic family corresponds to a distinct genome type and metabolic type. Sequence evolution is not independent of metabolic evolution. There is a balance among the interval frequency of codons in the genome, interval frequency of the corresponding amino acids in the proteome, and concentrations of tRNAs around the ribosome (Supp. Fig. 4dC). Mismatch among them results in low translation efficiency.

Based on a simple genome/proteome data processing method, a wealth of evolutionary phenomena have been observed in this study. Several aspects of the results are particularly noteworthy. First, the presence of TBP in nucleic acid sequences provides the fundamental basis for translation efficiency. Substantial TBP Family bias can benefit translation efficiency and hence separate cellular life forms and viruses. Second, protein foldability constrains genome features of cellular life forms around the four vertices of the tetrahedron. Protein foldability played a crucial role in the separation between cellular life forms and viruses. Given the existence of a forbidden zone for life in GenEvoSp, both TBP in nucleotide acids and protein foldability jointly shape the species-specific features of TBP Family bias.

#### Data processing methods

##### MATLAB script to generate Fig. 1

```
% Supplementary program to Plotting Fig. 1
% MATLAB_R2024a

clear; clc;

% 1:Bacteria; 2:Archaea; 3:Eukaryotes; 4:Viruses
sheetname={'bacteria';'archaea';'eukaryotes';'viruses'};
Nsheet=size(sheetname,1);
SuppDataCW{1}=readmatrix('Supplementary_Data_1_CodonWeight.xlsx','sheet',sheetname{1},'Range','F2');
SuppDataCW{2}=readmatrix('Supplementary_Data_1_CodonWeight.xlsx','sheet',sheetname{2},'Range','B2');
SuppDataCW{3}=readmatrix('Supplementary_Data_1_CodonWeight.xlsx','sheet',sheetname{3},'Range','B2');
SuppDataCW{4}=readmatrix('Supplementary_Data_1_CodonWeight.xlsx','sheet',sheetname{4},'Range','C2');
for i=1:Nsheet
   N_Taxa(i)=size(SuppDataCW{i},1);
end

for i=1:Nsheet
   M_Taxa(i)=0;
   for j=1:i
      M_Taxa(i)=M_Taxa(i)+N_Taxa(j);
   end
end

SuppDataCW_all=[];
for i=1:Nsheet
   SuppDataCW_all=[SuppDataCW_all;SuppDataCW{i}];
end

M0_Taxa=[0,M_Taxa];
for i=1:Nsheet
   NUM_TAXA{i}=[M0_Taxa(i)+1:M0_Taxa(i+1)];
end
```

```
% Calculate F-F bias from codon weights

cwdist=SuppDataCW_all';
N_species=size(cwdist,2);
% The numbers from 1 to 64 for the 64 codons, respectively: AAA, AAC, AAG, AAT, ACA, ACC, ACG, ACT, AGA,
    ↪  AGC, AGG, AGT, ATA, ATC, ATG, ATT, CAA, CAC, CAG, CAT, CCA, CCC, CCG, CCT, CGA, CGC, CGG, CGT,
    ↪ CTA, CTC, CTG, CTT, GAA, GAC, GAG, GAT, GCA, GCC, GCG, GCT, GGA, GGC, GGG, GGT, GTA, GTC, GTG,
    ↪ GTT, TAA, TAC, TAG, TAT, TCA, TCC, TCG, TCT, TGA, TGC, TGG, TGT, TTA, TTC, TTG TTT
% The biosynthetic familes in textbooks
Family_Ala_num_codon=[30 32 31 29 63 61 45 46 47 48 37 38 39 40]; %14
Family_Ser_num_codon=[53 54 55 56 10 12 58 60 41 42 43 44]; %12
Family_Glu_num_codon=[22 11 24 35 9 33 23 21 26 28 27 19 25 17 1 3]; %16
Family_Asp_num_codon=[34 2 36 4 7 5 15 13 6 14 8 16]; %12
Family_Phe_num_codon=[62 64 50 52 59]; %5
for spec=1:N_species
   cw0=0;
   for cod=Family_Ala_num_codon
      cw0=cw0+cwdist(cod,spec);
   end
   CW_Ala(spec)=cw0/14;

   cw0=0;
   for cod=Family_Ser_num_codon
      cw0=cw0+cwdist(cod,spec);
   end
   CW_Ser(spec)=cw0/12;

   cw0=0;
   for cod=Family_Glu_num_codon
      cw0=cw0+cwdist(cod,spec);
   end
   CW_Glu(spec)=cw0/16;

   cw0=0;
   for cod=Family_Asp_num_codon
      cw0=cw0+cwdist(cod,spec);
   end
   CW_Asp(spec)=cw0/12;

   cw0=0;
   for cod=Family_Phe_num_codon
      cw0=cw0+cwdist(cod,spec);
   end
   CW_Phe(spec)=cw0/5;
end
bias_Ala_Ser=CW_Ser-CW_Ala;
bias_Glu_Asp=CW_Asp-CW_Glu;
bias_Ser_Phe=CW_Phe-CW_Ser;
axis_x=bias_Glu_Asp;
axis_y=bias_Ala_Ser;
axis_z=bias_Ser_Phe;

disp('Plotting Fig 1')

figure(1)
WIDTH=18;
LENGTH=18;
axis([0,WIDTH,0,LENGTH])
set (gcf,'Units','centimeters','Position',[0,0,WIDTH,LENGTH], 'color','w')
axis off
tlsubplot=tiledlayout(2,2);
tlsubplot.TileSpacing='Compact';
tlsubplot.Padding='Compact';

disp('Plotting Fig 1a')

% Parameters
ParaMarkerSize_B=2;
ParaMarkerSize_A=2;
ParaMarkerSize_E=6;
ParaMarkerSize_V=1;
PARA_FontSize=7;
PARA_labelSize=7;
```

```
nexttile

hold on
plot3(axis_x(NUM_TAXA{4}),axis_y(NUM_TAXA{4}),axis_z(NUM_TAXA{4}),'c.','MarkerSize',ParaMarkerSize_V)
plot3(axis_x(NUM_TAXA{1}),axis_y(NUM_TAXA{1}),axis_z(NUM_TAXA{1}),'b.','MarkerSize',ParaMarkerSize_B)
plot3(axis_x(NUM_TAXA{2}),axis_y(NUM_TAXA{2}),axis_z(NUM_TAXA{2}),'r.','MarkerSize',ParaMarkerSize_A)
plot3(axis_x(NUM_TAXA{3}),axis_y(NUM_TAXA{3}),axis_z(NUM_TAXA{3}),'g.','MarkerSize',ParaMarkerSize_E)

legend('viruses','bacteria','archaea','eukaryotes','box','off','FontSize',6,'Position'
    ↪ ,[0.2,0.58,0.05,0.0666])
Rotation_x=-52;
Rotation_y=20;
Rotation_z=90;
alx=xlabel('F_{Glu}-F_{Asp} bias','FontSize',PARA_FontSize,'Rotation',Rotation_x,'Position'
    ↪ ,[0,-0.023,-0.11]);
aly=ylabel('F_{Ala}-F_{Ser} bias','FontSize',PARA_FontSize,'Rotation',Rotation_y,'Position'
    ↪ ,[0.027,0.0003,-0.1066]);
alz=zlabel('F_{Ser}-F_{Phe} bias','FontSize',PARA_FontSize,'Rotation',Rotation_z,'Position'
    ↪ ,[-0.0406,-0.0235,0]);
xtickangle(Rotation_x)
ytickangle(Rotation_y)
ztickangle(Rotation_z)
view(60,45)
axis([-0.039,0.028,-0.021,0.029,-0.08,0.08])
set(gca,'Linewidth',0.1,'GridAlpha',0.1,'Gridcolor',0.3*[1 1 1],'GridLineStyle','-')
set(gca,'XTick',[-0.02:0.02:0.02])
set(gca,'XTickLabel',{' -0.02 ',' 0 ',' 0.02 '},'FontSize',PARA_labelSize)
set(gca,'YTick',[-0.02:0.01:0.02])
set(gca,'YTickLabel',{' -0.02 ',' -0.01 ',' 0 ',' 0.01 ',' 0.02 '},'FontSize',PARA_labelSize)
set(gca,'ZTick',[-0.06:0.06:0.06])
set(gca,'ZTickLabel',{'-0.06','0','0.06'},'FontSize',PARA_labelSize)
box on
hold off
text_fs=10;
title('a                             ','FontSize',text_fs,'FontWeight','bold','HorizontalAlignment','
    ↪ right','VerticalAlignment','top')

disp('Plotting Fig 1b')

virusgenomesize10150=readmatrix('Supplementary_Data_1_CodonWeight.xlsx','sheet','viruses','Range','B2:
    ↪ B10151');
N10150=size(virusgenomesize10150,1);
NUM_TAXA_V1=[];
NUM_TAXA_V2=[];
for i=1:N10150
   if virusgenomesize10150(i)<23000
      NUM_TAXA_V1=[NUM_TAXA_V1,i];
   else
      NUM_TAXA_V2=[NUM_TAXA_V2,i];
   end
end
NUM_TAXA_V={M_Taxa(3)+NUM_TAXA_V1;M_Taxa(3)+NUM_TAXA_V2};

BaseComposition_ACGT=readmatrix('Supplementary_Data_1_CodonWeight.xlsx','sheet','bacteria','Range','B2:
    ↪ E1041');
GCcontent_bacteria=(BaseComposition_ACGT(:,2)+BaseComposition_ACGT(:,3))./sum(BaseComposition_ACGT,2);
N1040=size(GCcontent_bacteria,1);
NUM_TAXA_B1=[];
NUM_TAXA_B2=[];
for i=1:N1040
   if GCcontent_bacteria(i)>0.5
      NUM_TAXA_B1=[NUM_TAXA_B1,i];
   else
      NUM_TAXA_B2=[NUM_TAXA_B2,i];
   end
end
NUM_TAXA_B={NUM_TAXA_B1;NUM_TAXA_B2};

nexttile

hold on
```

```
plot3(axis_x(NUM_TAXA_V{1}),axis_y(NUM_TAXA_V{1}),axis_z(NUM_TAXA_V{1}),'c.','MarkerSize',
    ↪ ParaMarkerSize_V)
plot3(axis_x(NUM_TAXA_V{2}),axis_y(NUM_TAXA_V{2}),axis_z(NUM_TAXA_V{2}),'m.','MarkerSize',
    ↪ ParaMarkerSize_V)
plot3(axis_x(NUM_TAXA_B{1}),axis_y(NUM_TAXA_B{1}),axis_z(NUM_TAXA_B{1}),'b.','MarkerSize',
    ↪ ParaMarkerSize_B)
plot3(axis_x(NUM_TAXA_B{2}),axis_y(NUM_TAXA_B{2}),axis_z(NUM_TAXA_B{2}),'y.','MarkerSize',
    ↪ ParaMarkerSize_B)
plot3(axis_x(NUM_TAXA{2}),axis_y(NUM_TAXA{2}),axis_z(NUM_TAXA{2}),'r.','MarkerSize',ParaMarkerSize_A)
plot3(axis_x(NUM_TAXA{3}),axis_y(NUM_TAXA{3}),axis_z(NUM_TAXA{3}),'g.','MarkerSize',ParaMarkerSize_E)

legend('viruses (<23kbp)','viruses (>23kbp)','bacteria (GC%>0.5)','bacteria (GC%<0.5)','archaea','
    ↪ eukaryotes','box','off','FontSize',6,'Position',[0.67,0.578,0.05,0.1])
Rotation_x=-65;
Rotation_y=7;
Rotation_z=90;
alx=xlabel('F_{Glu}-F_{Asp} bias','FontSize',PARA_FontSize,'Rotation',Rotation_x,'Position'
    ↪ ,[-0.018,-0.021,-0.1159]);
aly=ylabel('F_{Ala}-F_{Ser} bias','FontSize',PARA_FontSize,'Rotation',Rotation_y,'Position'
    ↪ ,[0.027,0,-0.0991]);
alz=zlabel('F_{Ser}-F_{Phe} bias','FontSize',PARA_FontSize,'Rotation',Rotation_z,'Position'
    ↪ ,[-0.0399,-0.0235,0]);
xtickangle(Rotation_x)
ytickangle(Rotation_y)
ztickangle(Rotation_z)
view(75,30)
axis([-0.039,0.028,-0.021,0.029,-0.08,0.08])
set(gca,'Linewidth',0.1,'GridAlpha',0.1,'Gridcolor',0.3*[1 1 1],'GridLineStyle','-')
set(gca,'XTick',[-0.02:0.02:0.02])
set(gca,'XTickLabel',{' -0.02 ',' 0 ',' 0.02 '},'FontSize',PARA_labelSize)
set(gca,'YTick',[-0.02:0.01:0.02])
set(gca,'YTickLabel',{' -0.02 ',' -0.01 ',' 0 ',' 0.01 ',' 0.02 '},'FontSize',PARA_labelSize)
set(gca,'ZTick',[-0.06:0.06:0.06])
set(gca,'ZTickLabel',{'-0.06','0','0.06'},'FontSize',PARA_labelSize)
box on
hold off
text_fs=10;
title('b                                ','FontSize',text_fs,'FontWeight','bold','HorizontalAlignment','
    ↪ right','VerticalAlignment','top')

disp('Plotting Fig 1c')

% Parameters
ParaMarkerSize_B=2;
ParaMarkerSize_A=2;
ParaMarkerSize_E=6;
ParaMarkerSize_V=0.5;
ParaMarkerSize_simulation=3;
PARA_FontSize=7;
PARA_labelSize=7;

nexttile

hold on
plot3(axis_x(NUM_TAXA{4}),axis_y(NUM_TAXA{4}),axis_z(NUM_TAXA{4}),'c.','MarkerSize',ParaMarkerSize_V)
plot3(axis_x(NUM_TAXA{1}),axis_y(NUM_TAXA{1}),axis_z(NUM_TAXA{1}),'b.','MarkerSize',ParaMarkerSize_B)
plot3(axis_x(NUM_TAXA{2}),axis_y(NUM_TAXA{2}),axis_z(NUM_TAXA{2}),'r.','MarkerSize',ParaMarkerSize_A)
plot3(axis_x(NUM_TAXA{3}),axis_y(NUM_TAXA{3}),axis_z(NUM_TAXA{3}),'g.','MarkerSize',ParaMarkerSize_E)
bias_A_rich=readmatrix('Supplementary_Data_4_ifaa_ifc_simulation.xlsx','sheet','simulation Fig 1c','
    ↪ Range','A2:C3001');
bias_C_rich=readmatrix('Supplementary_Data_4_ifaa_ifc_simulation.xlsx','sheet','simulation Fig 1c','
    ↪ Range','D2:F3001');
bias_G_rich=readmatrix('Supplementary_Data_4_ifaa_ifc_simulation.xlsx','sheet','simulation Fig 1c','
    ↪ Range','G2:I3001');
bias_T_rich=readmatrix('Supplementary_Data_4_ifaa_ifc_simulation.xlsx','sheet','simulation Fig 1c','
    ↪ Range','J2:L3001');
bias_random=readmatrix('Supplementary_Data_4_ifaa_ifc_simulation.xlsx','sheet','simulation Fig 1c','
    ↪ Range','M2:O12001');
bias_xyz=bias_A_rich;
plot3(bias_xyz(:,1),bias_xyz(:,2),bias_xyz(:,3),'k.','MarkerSize',1,'LineWidth',0.1);
bias_xyz=bias_C_rich;
plot3(bias_xyz(:,1),bias_xyz(:,2),bias_xyz(:,3),'k.','MarkerSize',1,'LineWidth',0.1);
bias_xyz=bias_G_rich;
```

```
plot3(bias_xyz(:,1),bias_xyz(:,2),bias_xyz(:,3),'k.','MarkerSize',1,'LineWidth',0.1);
bias_xyz=bias_T_rich;
plot3(bias_xyz(:,1),bias_xyz(:,2),bias_xyz(:,3),'k.','MarkerSize',1,'LineWidth',0.1);
bias_xyz=bias_random;
plot3(bias_xyz(:,1),bias_xyz(:,2),bias_xyz(:,3),'k.','MarkerSize',1,'LineWidth',0.1);


legend('viruses','bacteria','archaea','eukaryotes','random genomes','box','off','FontSize',6,'Position'
    ↪ ,[0.13,0.35,0.05,0.09])
Rotation_x=65;
Rotation_y=-8;
Rotation_z=90;
alx=xlabel('F_{Glu}-F_{Asp} bias','FontSize',PARA_FontSize,'Rotation',Rotation_x,'Position'
    ↪ ,[0.022,-0.061,-0.1]);
aly=ylabel('F_{Ala}-F_{Ser} bias','FontSize',PARA_FontSize,'Rotation',Rotation_y,'Position'
    ↪ ,[0.064,0.01,-0.1]);
alz=zlabel('F_{Ser}-F_{Phe} bias','FontSize',PARA_FontSize,'Rotation',Rotation_z,'Position'
    ↪ ,[0.081,0.085,0.05]);
xtickangle(Rotation_x)
ytickangle(Rotation_y)
ztickangle(Rotation_z)
view(-102.8,-33.5)
axis([-0.063,0.0541,-0.05,0.085,-0.1,0.2])
set(gca,'Linewidth',0.1,'GridAlpha',0.1,'Gridcolor',0.3*[1 1 1],'GridLineStyle','-')
set(gca,'XTick',[-0.06:0.03:0.03])
set(gca,'XTickLabel',{' ',' -0.03 ',' 0 ',' 0.03 '},'FontSize',PARA_labelSize)
set(gca,'YTick',[-0.05:0.05:0.05])
set(gca,'YTickLabel',{' -0.05 ',' 0 ',' 0.05 '},'FontSize',PARA_labelSize)
set(gca,'ZTick',[-0.1:0.1:0.2])
set(gca,'ZTickLabel',{' -0.1 ',' 0 ',' 0.1 ',' 0.2 '},'FontSize',PARA_labelSize)
box on
hold off
text_fs=10;
title('c                              ','FontSize',text_fs,'FontWeight','bold','HorizontalAlignment','
    ↪ right','VerticalAlignment','top')

disp('Plotting Fig 1d')

clear;

PARA_FontSize=7;
PARA_labelSize=7;

nexttile

hold on
plot3(0,0,1,'c.','MarkerSize',3)
plot3(0,0,1,'b.','MarkerSize',3)
plot3(0,0,1,'y.','MarkerSize',3)
plot3(0,0,1,'r.','MarkerSize',3)
plot3(0,0,1,'g.','MarkerSize',3)
bias_V=readmatrix('Supplementary_Data_4_ifaa_ifc_simulation.xlsx','sheet','simulation Fig 1d','Range','
    ↪ A2:C12001');
bias_Bh=readmatrix('Supplementary_Data_4_ifaa_ifc_simulation.xlsx','sheet','simulation Fig 1d','Range','
    ↪ D2:F3001');
bias_Bl=readmatrix('Supplementary_Data_4_ifaa_ifc_simulation.xlsx','sheet','simulation Fig 1d','Range','
    ↪ G2:I3001');
bias_A=readmatrix('Supplementary_Data_4_ifaa_ifc_simulation.xlsx','sheet','simulation Fig 1d','Range','
    ↪ J2:L3001');
bias_E=readmatrix('Supplementary_Data_4_ifaa_ifc_simulation.xlsx','sheet','simulation Fig 1d','Range','
    ↪ M2:O3001');
bias_xyz=bias_V;
plot3(bias_xyz(:,1),bias_xyz(:,2),bias_xyz(:,3),'c.','MarkerSize',1);
bias_xyz=bias_Bh;
plot3(bias_xyz(:,1),bias_xyz(:,2),bias_xyz(:,3),'b.','MarkerSize',1);
bias_xyz=bias_Bl;
plot3(bias_xyz(:,1),bias_xyz(:,2),bias_xyz(:,3),'y.','MarkerSize',1);
bias_xyz=bias_A;
plot3(bias_xyz(:,1),bias_xyz(:,2),bias_xyz(:,3),'r.','MarkerSize',1);
bias_xyz=bias_E;
plot3(bias_xyz(:,1),bias_xyz(:,2),bias_xyz(:,3),'g.','MarkerSize',1);
```

```
legend('low bias','F_{Ala} rich','F_{Phe} rich','F_{Ser} rich','F_{Asp} rich','box','off','FontSize',6,'
    ↪ Position',[0.66,0.06,0.05,0.095])
Rotation_x=-49;
Rotation_y=20;
Rotation_z=90;
alx=xlabel('F_{Glu}-F_{Asp} bias','FontSize',PARA_FontSize,'Rotation',Rotation_x,'Position'
    ↪ ,[0.015,-0.012,-0.081]);
aly=ylabel('F_{Ala}-F_{Ser} bias','FontSize',PARA_FontSize,'Rotation',Rotation_y,'Position'
    ↪ ,[0.038,0.005,-0.0866]);
alz=zlabel('F_{Ser}-F_{Phe} bias','FontSize',PARA_FontSize,'Rotation',Rotation_z,'Position'
    ↪ ,[-0.0406,-0.0135,0]);
xtickangle(Rotation_x)
ytickangle(Rotation_y)
ztickangle(Rotation_z)
view(60,45)
axis([-0.039,0.041,-0.011,0.022,-0.061,0.08])
set(gca,'Linewidth',0.1,'GridAlpha',0.1,'Gridcolor',0.3*[1 1 1],'GridLineStyle','-')
set(gca,'XTick',[-0.025:0.025:0.025])
set(gca,'XTickLabel',{' -0.025 ',' 0 ','0.025'},'FontSize',PARA_labelSize)
set(gca,'YTick',[0:0.015:0.015])
set(gca,'YTickLabel',{' 0 ',' 0.015 '},'FontSize',PARA_labelSize)
set(gca,'ZTick',[-0.06:0.06:0.06])
set(gca,'ZTickLabel',{'-0.06','0','0.06'},'FontSize',PARA_labelSize)
box on
hold off
text_fs=10;
title('d                              ','FontSize',text_fs,'FontWeight','bold','HorizontalAlignment','
    ↪ right','VerticalAlignment','top')

disp('Saving figures')
n='1';
saveas(gcf,['Fig_',n,'.fig']);
exportgraphics(gcf,['Fig_',n,'.pdf'],'Resolution',1500)
exportgraphics(gcf,['Fig_',n,'.tif'],'Resolution',1500)
exportgraphics(gcf,['Fig_',n,'.png'],'Resolution',1500)
exportgraphics(gcf,['Fig_',n,'_r1500.eps'],'Resolution',1500)

disp('Finished')
```

### MATLAB script to generate Fig. 2

```
% Supplementary program to Plotting Fig. 2
% MATLAB_R2024a

clear; clc;

% 1:America; 2:Australian; 3:China; 4:Iceland; 5:Italy; 6:New Zealand; 7:Spain;
% 8:Beta; 9:Delta; 10:Alpha; 11:Gamma; 12:Epsilon; 13:BA1; 14:XBB; 15:BA3; 16:BA4; 17:BA5; 18:Zeta; 19:
    ↪ Eta; 20:Lambda; 21:Mu; 22:Iota; 23:KP2; 24:Omicron random; 25:Omicron 2024; 26:SARS-CoV-2 NCBI;
% 27:China 2019; 28:China Jan 2020; 29:China Feb 2020; 30:China Mar 2020; 31:China Apr-Dec 2020;
% 32:SARS 2003; 33:SARS 2004; 34:SARS 2005; 35:Bat Cov'
sheetname={'America';'Australian';'China';'Iceland';'Italy';'New Zealand';'Spain';'Beta';'Delta';'Alpha'
    ↪ ;'Gamma';'Epsilon';'BA1';'XBB';'BA3';'BA4';'BA5';'Zeta';'Eta';'Lambda';'Mu';'Iota';'KP2';'Omicron
    ↪  random';'Omicron 2024';'SARS-CoV-2 NCBI';'China 2019';'China Jan 2020';'China Feb 2020';'China
    ↪ Mar 2020';'China Apr-Dec 2020';'SARS 2003';'SARS 2004';'SARS 2005';'Bat Cov'};
Nsheet=size(sheetname,1);

disp('Reading data from Supplementary_Data_2_CodonWeight_SARS-CoV-2.xlsx')
for i=1
   sheetname0=sheetname{i};
   SuppDataCW{i}=readmatrix('Supplementary_Data_2_CodonWeight_SARS-CoV-2.xlsx','sheet',sheetname0,'Range
       ↪ ','E2');
   n_of_the_total_35=i
end
for i=2:Nsheet
   sheetname0=sheetname{i};
```

```
    SuppDataCW{i}=readmatrix('Supplementary_Data_2_CodonWeight_SARS-CoV-2.xlsx','sheet',sheetname0,'Range
        ↪ ','B2');
    n_of_the_total_35=i
end
save('Supplementary_Data_2_CodonWeight_SARSCoV2.mat','SuppDataCW')
% load Supplementary_Data_2_CodonWeight_SARSCoV2.mat

% 36:Eukaryotes
SuppDataCW{36}=readmatrix('Supplementary_Data_1_CodonWeight.xlsx','sheet','eukaryotes','Range','B2')
Nsheet=Nsheet+1;

for i=1:Nsheet
    N_Taxa(i)=size(SuppDataCW{i},1);
end
for i=1:Nsheet
    M_Taxa(i)=0;
    for j=1:i
        M_Taxa(i)=M_Taxa(i)+N_Taxa(j);
    end
end
SuppDataCW_all=[];
for i=1:Nsheet
    SuppDataCW_all=[SuppDataCW_all;SuppDataCW{i}];
end
M0_Taxa=[0,M_Taxa];
for i=1:Nsheet
    NUM_TAXA{i}=[M0_Taxa(i)+1:M0_Taxa(i+1)];
end

% Calculate F-F bias from codon weights

cwdist=SuppDataCW_all';
N_species=size(cwdist,2);

% The numbers from 1 to 64 for the 64 codons, respectively: AAA, AAC, AAG, AAT, ACA, ACC, ACG, ACT, AGA,
     ↪  AGC, AGG, AGT, ATA, ATC, ATG, ATT, CAA, CAC, CAG, CAT, CCA, CCC, CCG, CCT, CGA, CGC, CGG, CGT,
     ↪ CTA, CTC, CTG, CTT, GAA, GAC, GAG, GAT, GCA, GCC, GCG, GCT, GGA, GGC, GGG, GGT, GTA, GTC, GTG,
     ↪ GTT, TAA, TAC, TAG, TAT, TCA, TCC, TCG, TCT, TGA, TGC, TGG, TGT, TTA, TTC, TTG TTT
% The biosynthetic familes in textbooks
Family_Ala_num_codon=[30 32 31 29 63 61 45 46 47 48 37 38 39 40]; %14
Family_Ser_num_codon=[53 54 55 56 10 12 58 60 41 42 43 44]; %12
Family_Glu_num_codon=[22 11 24 35 9 33 23 21 26 28 27 19 25 17 1 3]; %16
Family_Asp_num_codon=[34 2 36 4 7 5 15 13 6 14 8 16]; %12
Family_Phe_num_codon=[62 64 50 52 59]; %5
for spec=1:N_species
    cw0=0;
    for cod=Family_Ala_num_codon
        cw0=cw0+cwdist(cod,spec);
    end
    CW_Ala(spec)=cw0/14;

    cw0=0;
    for cod=Family_Ser_num_codon
        cw0=cw0+cwdist(cod,spec);
    end
    CW_Ser(spec)=cw0/12;

    cw0=0;
    for cod=Family_Glu_num_codon
        cw0=cw0+cwdist(cod,spec);
    end
    CW_Glu(spec)=cw0/16;

    cw0=0;
    for cod=Family_Asp_num_codon
        cw0=cw0+cwdist(cod,spec);
    end
    CW_Asp(spec)=cw0/12;

    cw0=0;
    for cod=Family_Phe_num_codon
        cw0=cw0+cwdist(cod,spec);
    end
```

```
    CW_Phe(spec)=cw0/5;
end
bias_Ala_Ser=CW_Ser-CW_Ala;
bias_Glu_Asp=CW_Asp-CW_Glu;
bias_Ser_Phe=CW_Phe-CW_Ser;
axis_x=bias_Glu_Asp;
axis_y=bias_Ala_Ser;
axis_z=bias_Ser_Phe;

disp('Plotting Fig 2')

figure(2)
WIDTH=18;
LENGTH=18;
axis([0,WIDTH,0,LENGTH])
set (gcf,'Units','centimeters','Position',[0,0,WIDTH,LENGTH], 'color','w')
axis off
tlsubplot=tiledlayout(2,2);
tlsubplot.TileSpacing='Compact';
tlsubplot.Padding='Compact';

disp('Plotting Fig 2a')

nexttile

hold on
MarkerAndColorLegend=['b.';'g.'];
for i=1:2
    plot3(0,0,1,MarkerAndColorLegend(i,:),'MarkerSize',3)
end
plot3(axis_x(NUM_TAXA{1}),axis_y(NUM_TAXA{1}),axis_z(NUM_TAXA{1}),'b.','MarkerSize',0.5)
plot3(axis_x(NUM_TAXA{25}),axis_y(NUM_TAXA{25}),axis_z(NUM_TAXA{25}),'g.','MarkerSize',0.5)
legend('America 2020','Omicron 2024','box','off','FontSize',6,'Position',[0.12,0.888,0.05,0.04])

PARA_FontSize=7;
PARA_labelSize=7;
view(1.1652,-1.7251)
axis([-0.008,-0.002,-0.0038,-0.0033,0.046,0.059])
set(gca,'Linewidth',0.1,'GridAlpha',0.1,'Gridcolor',0.3*[1 1 1],'GridLineStyle','-')
set(gca,'XTick',[-0.007:0.002:-0.003])
set(gca,'XTickLabel',{' -0.007 ',' -0.005 ',' -0.003 '},'FontSize',PARA_labelSize)
set(gca,'YTick',[-0.0038:0.0005:-0.0033])
set(gca,'YTickLabel',{'-0.0038 ','-0.0033 '},'FontSize',PARA_labelSize)
set(gca,'ZTick',[0.047:0.005:0.057])
set(gca,'ZTickLabel',{'0.047 ','0.052 ','0.057 '},'FontSize',PARA_labelSize)
Rotation_x=0;
Rotation_y=45;
Rotation_z=90;
alx=xlabel('F_{Glu}-F_{Asp} bias','FontSize',PARA_FontSize,'Rotation',Rotation_x,'Position'
    ↪ ,[-0.005,-0.00395,0.04439],'HorizontalAlignment','center');
aly=ylabel('F_{Ala}-F_{Ser} bias','FontSize',PARA_FontSize,'Rotation',Rotation_y,'Position'
    ↪ ,[-0.008,-0.00355,0.04481],'HorizontalAlignment','center');
alz=zlabel('F_{Ser}-F_{Phe} bias','FontSize',PARA_FontSize,'Rotation',Rotation_z,'Position'
    ↪ ,[-0.00839,-0.0038,0.052],'HorizontalAlignment','center');
xtickangle(Rotation_x)
ytickangle(Rotation_y)
ztickangle(Rotation_z)
box on
hold off
text_fs=10;
title('a                              ','FontSize',text_fs,'FontWeight','bold','HorizontalAlignment','
    ↪ right','VerticalAlignment','top')

disp('Plotting Fig 2b')

% Parameters
ParaMarkerSize_V=0.5;
ParaMarkerSize_V1=0.5;
ParaMarkerSize_V2=2;
ParaLineWidth_1=0.1;
ParaLineWidth_2=0.1;
PARA_FontSize=7;
PARA_labelSize=7;
```

```
nexttile

hold on
MarkerAndColorLegend=['b.';'m.';'r.';'g.';'k.';'y.';'c.';'k+';'b+';'m+';'c+';'r+';'mx';'rx';'gx';'bx';'
    ↪ cx';'yx'];
for i=1:18
   plot3(0,0,1,MarkerAndColorLegend(i,:),'MarkerSize',3)
end
plot3(axis_x(NUM_TAXA{1}),axis_y(NUM_TAXA{1}),axis_z(NUM_TAXA{1}),MarkerAndColorLegend(1,:),'MarkerSize'
    ↪ ,ParaMarkerSize_V)
plot3(axis_x(NUM_TAXA{2}),axis_y(NUM_TAXA{2}),axis_z(NUM_TAXA{2}),MarkerAndColorLegend(2,:),'MarkerSize'
    ↪ ,ParaMarkerSize_V)
plot3(axis_x(NUM_TAXA{8}),axis_y(NUM_TAXA{8}),axis_z(NUM_TAXA{8}),MarkerAndColorLegend(3,:),'MarkerSize'
    ↪ ,ParaMarkerSize_V2,'LineWidth',ParaLineWidth_2)
plot3(axis_x(NUM_TAXA{9}),axis_y(NUM_TAXA{9}),axis_z(NUM_TAXA{9}),MarkerAndColorLegend(4,:),'MarkerSize'
    ↪ ,ParaMarkerSize_V2,'LineWidth',ParaLineWidth_2)
plot3(axis_x(NUM_TAXA{10}),axis_y(NUM_TAXA{10}),axis_z(NUM_TAXA{10}),MarkerAndColorLegend(5,:),'
    ↪ MarkerSize',ParaMarkerSize_V2,'LineWidth',ParaLineWidth_1)
plot3(axis_x(NUM_TAXA{11}),axis_y(NUM_TAXA{11}),axis_z(NUM_TAXA{11}),MarkerAndColorLegend(6,:),'
    ↪ MarkerSize',ParaMarkerSize_V2,'LineWidth',ParaLineWidth_2)
plot3(axis_x(NUM_TAXA{12}),axis_y(NUM_TAXA{12}),axis_z(NUM_TAXA{12}),MarkerAndColorLegend(7,:),'
    ↪ MarkerSize',ParaMarkerSize_V2,'LineWidth',ParaLineWidth_2)
plot3(axis_x(NUM_TAXA{13}),axis_y(NUM_TAXA{13}),axis_z(NUM_TAXA{13}),MarkerAndColorLegend(8,:),'
    ↪ MarkerSize',ParaMarkerSize_V1,'LineWidth',ParaLineWidth_1)
plot3(axis_x(NUM_TAXA{14}),axis_y(NUM_TAXA{14}),axis_z(NUM_TAXA{14}),MarkerAndColorLegend(9,:),'
    ↪ MarkerSize',ParaMarkerSize_V1,'LineWidth',ParaLineWidth_1)
plot3(axis_x(NUM_TAXA{15}),axis_y(NUM_TAXA{15}),axis_z(NUM_TAXA{15}),MarkerAndColorLegend(10,:),'
    ↪ MarkerSize',ParaMarkerSize_V1,'LineWidth',ParaLineWidth_1)
plot3(axis_x(NUM_TAXA{16}),axis_y(NUM_TAXA{16}),axis_z(NUM_TAXA{16}),MarkerAndColorLegend(11,:),'
    ↪ MarkerSize',ParaMarkerSize_V1,'LineWidth',ParaLineWidth_1)
plot3(axis_x(NUM_TAXA{17}),axis_y(NUM_TAXA{17}),axis_z(NUM_TAXA{17}),MarkerAndColorLegend(12,:),'
    ↪ MarkerSize',ParaMarkerSize_V1,'LineWidth',ParaLineWidth_1)
plot3(axis_x(NUM_TAXA{18}),axis_y(NUM_TAXA{18}),axis_z(NUM_TAXA{18}),MarkerAndColorLegend(13,:),'
    ↪ MarkerSize',ParaMarkerSize_V1,'LineWidth',ParaLineWidth_2)
plot3(axis_x(NUM_TAXA{19}),axis_y(NUM_TAXA{19}),axis_z(NUM_TAXA{19}),MarkerAndColorLegend(14,:),'
    ↪ MarkerSize',ParaMarkerSize_V1,'LineWidth',ParaLineWidth_2)
plot3(axis_x(NUM_TAXA{20}),axis_y(NUM_TAXA{20}),axis_z(NUM_TAXA{20}),MarkerAndColorLegend(15,:),'
    ↪ MarkerSize',ParaMarkerSize_V1,'LineWidth',ParaLineWidth_2)
plot3(axis_x(NUM_TAXA{21}),axis_y(NUM_TAXA{21}),axis_z(NUM_TAXA{21}),MarkerAndColorLegend(16,:),'
    ↪ MarkerSize',ParaMarkerSize_V1,'LineWidth',ParaLineWidth_2)
plot3(axis_x(NUM_TAXA{22}),axis_y(NUM_TAXA{22}),axis_z(NUM_TAXA{22}),MarkerAndColorLegend(17,:),'
    ↪ MarkerSize',ParaMarkerSize_V1,'LineWidth',ParaLineWidth_2)
plot3(axis_x(NUM_TAXA{23}),axis_y(NUM_TAXA{23}),axis_z(NUM_TAXA{23}),MarkerAndColorLegend(18,:),'
    ↪ MarkerSize',ParaMarkerSize_V1,'LineWidth',ParaLineWidth_1)

legend('America','Australian','Beta','Delta','Alpha','Gamma','Epsilon','BA1','XBB','BA3','BA4','BA5','
    ↪ Zeta','Eta','Lambda','Mu','Iota','KP2','box','off','FontSize',5,'Position',[0.465,0.71,0.05,0.2])
view(1.1652,-1.7251)
axis([-0.0066,-0.0023,-0.0038,-0.0033,0.0486,0.056])
set(gca,'Linewidth',0.1,'GridAlpha',0.1,'Gridcolor',0.3*[1 1 1],'GridLineStyle','-')
set(gca,'XTick',[-0.006:0.002:-0.004])
set(gca,'XTickLabel',{' -0.006 ',' -0.004 '},'FontSize',PARA_labelSize)
set(gca,'YTick',[-0.0038:0.0005:-0.0033])
set(gca,'YTickLabel',{'-0.0038 ','-0.0033 '},'FontSize',PARA_labelSize)
set(gca,'ZTick',[0.049:0.003:0.055])
set(gca,'ZTickLabel',{'0.049 ','0.052 ','0.055 '},'FontSize',PARA_labelSize)
Rotation_x=0;
Rotation_y=46;
Rotation_z=90;
alx=xlabel('F_{Glu}-F_{Asp} bias','FontSize',PARA_FontSize,'Rotation',Rotation_x,'Position'
    ↪ ,[-0.004,-0.00395,0.04765],'HorizontalAlignment','center');
aly=ylabel('F_{Ala}-F_{Ser} bias','FontSize',PARA_FontSize,'Rotation',Rotation_y,'Position'
    ↪ ,[-0.00705,-0.0038,0.0485],'HorizontalAlignment','center');
alz=zlabel('F_{Ser}-F_{Phe} bias','FontSize',PARA_FontSize,'Rotation',Rotation_z,'Position'
    ↪ ,[-0.0068,-0.0038,0.05],'HorizontalAlignment','center');
xtickangle(Rotation_x)
ytickangle(Rotation_y)
ztickangle(Rotation_z)
box on
hold off
text_fs=10;
```

```
title('b                              ','FontSize',text_fs,'FontWeight','bold','HorizontalAlignment','right','VerticalAlignment','top')

disp('Plotting Fig 2c')

nexttile

hold on
MarkerAndColorLegend=['kx';'bx';'cx';'r.';'ro';'b.';'g.';'y.';'rp'];
for i=1:9
   plot3(0,0,1,MarkerAndColorLegend(i,:),'MarkerSize',3,'LineWidth',0.1)
end
plot3(axis_x(NUM_TAXA{1}),axis_y(NUM_TAXA{1}),axis_z(NUM_TAXA{1}),'b.','MarkerSize',0.5)
plot3(axis_x(NUM_TAXA{25}),axis_y(NUM_TAXA{25}),axis_z(NUM_TAXA{25}),'g.','MarkerSize',0.5)
plot3(axis_x(NUM_TAXA{26}),axis_y(NUM_TAXA{26}),axis_z(NUM_TAXA{26}),'y.','MarkerSize',0.5)
plot3(axis_x(NUM_TAXA{32}),axis_y(NUM_TAXA{32}),axis_z(NUM_TAXA{32}),'kx','MarkerSize',1,'LineWidth',0.1)
plot3(axis_x(NUM_TAXA{33}),axis_y(NUM_TAXA{33}),axis_z(NUM_TAXA{33}),'bx','MarkerSize',1,'LineWidth',0.1)
plot3(axis_x(NUM_TAXA{34}),axis_y(NUM_TAXA{34}),axis_z(NUM_TAXA{34}),'cx','MarkerSize',1,'LineWidth',0.1)
plot3(axis_x(NUM_TAXA{35}),axis_y(NUM_TAXA{35}),axis_z(NUM_TAXA{35}),'r.','MarkerSize',2)
N_RaTG13=17;
plot3(axis_x(NUM_TAXA{35}(N_RaTG13)),axis_y(NUM_TAXA{35}(N_RaTG13)),axis_z(NUM_TAXA{35}(N_RaTG13)),'ro','MarkerSize',5,'LineWidth',0.1)
N_Homo_sapiens=1;
plot3(axis_x(NUM_TAXA{36}(N_Homo_sapiens)),axis_y(NUM_TAXA{36}(N_Homo_sapiens)),axis_z(NUM_TAXA{36}(N_Homo_sapiens)),'rp','MarkerSize',5,'LineWidth',0.1)

legend('SARS-CoV 2003','SARS-CoV 2004','SARS-CoV 2005','Bat CoV','RaTG13','SARS-CoV-2 America 2020','SARS-CoV-2 Omicron 2024','SARS-CoV-2 NCBI','Homo sapiens','box','off','FontSize',6,'Position',[0.105,0.1,0.05,0.11])
PARA_FontSize=7;
PARA_labelSize=7;
view(5.65,3.85)
axis([-0.01,0.005,-0.0041,0,-0.005,0.062])
set(gca,'Linewidth',0.1,'GridAlpha',0.1,'Gridcolor',0.3*[1 1 1],'GridLineStyle','-')
set(gca,'XTick',[-0.006:0.004:0.002])
set(gca,'XTickLabel',{' -0.006 ',' -0.002 ',' 0.002 '},'FontSize',PARA_labelSize)
set(gca,'YTick',[-0.004:0.004:0])
set(gca,'YTickLabel',{'-0.004 ',' 0 '},'FontSize',PARA_labelSize)
set(gca,'ZTick',[0:0.02:0.06])
set(gca,'ZTickLabel',{' 0 ',' 0.02 ',' 0.04 ',' 0.06 '},'FontSize',PARA_labelSize)
Rotation_x=0;
Rotation_y=36;
Rotation_z=90;
alx=xlabel('F_{Glu}-F_{Asp} bias','FontSize',PARA_FontSize,'Rotation',Rotation_x,'Position',[-0.0025,-0.0041,-0.0103],'HorizontalAlignment','center');
aly=ylabel('F_{Ala}-F_{Ser} bias ','FontSize',PARA_FontSize,'Rotation',Rotation_y,'Position',[0.006,-0.002,-0.005],'HorizontalAlignment','center');
alz=zlabel('F_{Ser}-F_{Phe} bias','FontSize',PARA_FontSize,'Rotation',Rotation_z,'Position',[-0.0108,-0.0041,0.03],'HorizontalAlignment','center');
xtickangle(Rotation_x)
ytickangle(Rotation_y)
ztickangle(Rotation_z)
box on
hold off
text_fs=10;
title('c                              ','FontSize',text_fs,'FontWeight','bold','HorizontalAlignment','right','VerticalAlignment','top')

disp('Plotting Fig 2d')

clear;
nexttile

hold on
plot3(0,0,1,'b.','MarkerSize',3)
plot3(0,0,1,'g.','MarkerSize',3)
bias_earlyKite=readmatrix('Supplementary_Data_4_ifaa_ifc_simulation.xlsx','sheet','simulation Fig 2d','Range','A2:C30001');
bias_lateKite=readmatrix('Supplementary_Data_4_ifaa_ifc_simulation.xlsx','sheet','simulation Fig 2d','Range','D2:F30001');
```

```
bias_xyz=bias_earlyKite;
plot3(bias_xyz(:,1),bias_xyz(:,2),bias_xyz(:,3),'b.','MarkerSize',0.5);
bias_xyz=bias_lateKite;
plot3(bias_xyz(:,1),bias_xyz(:,2),bias_xyz(:,3),'g.','MarkerSize',0.5);

legend('Early Kite','Late Kite','box','off','FontSize',6,'Position',[0.58,0.4,0.05,0.04])
PARA_FontSize=7;
PARA_labelSize=7;
view(1.1652,-1.7251)
axis([-0.0062,-0.001686,-0.0017,-0.0014,0.037,0.043])
set(gca,'Linewidth',0.1,'GridAlpha',0.1,'Gridcolor',0.3*[1 1 1],'GridLineStyle','-')
set(gca,'XTick',[-0.006:0.002:-0.002])
set(gca,'XTickLabel',{' -0.006 ',' -0.004 ',' -0.002 '},'FontSize',PARA_labelSize)
set(gca,'YTick',[-0.0017:0.0003:-0.0014])
set(gca,'YTickLabel',{'-0.0017 ','-0.0014 '},'FontSize',PARA_labelSize)
set(gca,'ZTick',[0.038:0.002:0.042])
set(gca,'ZTickLabel',{'0.038 ','0.04 ','0.042 '},'FontSize',PARA_labelSize)
Rotation_x=0;
Rotation_y=36;
Rotation_z=90;
alx=xlabel('F_{Glu}-F_{Asp} bias','FontSize',PARA_FontSize,'Rotation',Rotation_x,'Position'
    ↪ ,[-0.004,-0.0014,0.03655],'HorizontalAlignment','center');
aly=ylabel('F_{Ala}-F_{Ser} bias','FontSize',PARA_FontSize,'Rotation',Rotation_y,'Position'
    ↪ ,[-0.00669,-0.0017,0.0369],'HorizontalAlignment','center');
alz=zlabel('F_{Ser}-F_{Phe} bias','FontSize',PARA_FontSize,'Rotation',Rotation_z,'Position'
    ↪ ,[-0.00635,-0.0017,0.04],'HorizontalAlignment','center');
xtickangle(Rotation_x)
ytickangle(Rotation_y)
ztickangle(Rotation_z)
box on
hold off
text_fs=10;
title('d                              ','FontSize',text_fs,'FontWeight','bold','HorizontalAlignment','
    ↪ right','VerticalAlignment','top')

disp('Saving figures')
n='2';
saveas(gcf,['Fig_',n,'.fig']);
exportgraphics(gcf,['Fig_',n,'.pdf'],'Resolution',1500)
exportgraphics(gcf,['Fig_',n,'.tif'],'Resolution',1500)
exportgraphics(gcf,['Fig_',n,'.png'],'Resolution',1500)
exportgraphics(gcf,['Fig__',n,'.eps'],'Resolution',1500)
exportgraphics(gcf,['Fig_',n,'_r3000.pdf'],'Resolution',3000)
exportgraphics(gcf,['Fig_',n,'_r3000.tif'],'Resolution',3000)
exportgraphics(gcf,['Fig_',n,'_r3000.png'],'Resolution',3000)
exportgraphics(gcf,['Fig__',n,'_r3000.eps'],'Resolution',3000)

disp('Finished')
```

### MATLAB script to generate Fig. 3

```
% Supplementary program to Plotting Fig. 3
% MATLAB_R2024a

clear; clc;

% 1:bacteria; 2:archaea; 3:fungi; 4:plants; 5:protozoa; 6:invertebrates; 7:vertebrates excluding mammals
    ↪  (vertebrates other); 8:mammals; 9:RNA viruses; 10:DNA viruses
taxa={'bacteria';'archaea';'fungi';'plants';'protozoa';'invertebrates';'vertebrates excluding mammals';'
    ↪ mammals';'RNA viruses';'DNA viruses'};
Num_Taxa_Data=size(taxa,1);

for i=1:Num_Taxa_Data
   taxa0=taxa{i};
   SuppDataAAW{i}=readmatrix('Supplementary_Data_3_AminoAcidWeight.xlsx','sheet',taxa0,'Range','B2');
end
for i=1:Num_Taxa_Data
```

```
   N_Taxa(i)=size(SuppDataAAW{i},1);
end
for i=1:Num_Taxa_Data
   M_Taxa(i)=0;
   for j=1:i
      M_Taxa(i)=M_Taxa(i)+N_Taxa(j);
   end
end
SuppDataAAW_all_new=[];
for i=1:Num_Taxa_Data
   SuppDataAAW_all_new=[SuppDataAAW_all_new;SuppDataAAW{i}];
end
M0_Taxa=[0,M_Taxa];
for i=1:Num_Taxa_Data
   NUM_TAXA{i}=[M0_Taxa(i)+1:M0_Taxa(i+1)];
end

% Calculate tildeF-tildeF bias from amino acid weights

aawdist=SuppDataAAW_all_new';
N_species=size(aawdist,2);

% The numbers from 1 to 20 for the 20 amino acids, respectively: A, C, D, E, F, G, H, I, K, L, M, N, P,
    ↪ Q, R, S, T, V, W, Y,
% namely, Ala, Cys, Asp, Glu, Phe, Gly, His, Ile, Lys, Leu, Met, Asn, Pro, Gln, Arg, Ser, Thr, Val, Trp,
    ↪ Tyr
% The biosynthetic familes in textbooks
Family_Ala_num_aminoacid=[18 1 10]; %3
Family_Ser_num_aminoacid=[16 6 2]; %3
Family_Glu_num_aminoacid=[4 13 15 14 9]; %5
Family_Asp_num_aminoacid=[3 17 8 11 12]; %5
Family_Phe_num_aminoacid=[5 19 20]; %3
Family_His_num_aminoacid=[7]; %1
for spec=1:N_species
   aaw0=0;
   for cod=Family_Ala_num_aminoacid
      aaw0=aaw0+aawdist(cod,spec);
   end
   AAW_Ala(spec)=aaw0/3;

   aaw0=0;
   for cod=Family_Ser_num_aminoacid
      aaw0=aaw0+aawdist(cod,spec);
   end
   AAW_Ser(spec)=aaw0/3;

   aaw0=0;
   for cod=Family_Glu_num_aminoacid
      aaw0=aaw0+aawdist(cod,spec);
   end
   AAW_Glu(spec)=aaw0/5;

   aaw0=0;
   for cod=Family_Asp_num_aminoacid
      aaw0=aaw0+aawdist(cod,spec);
   end
   AAW_Asp(spec)=aaw0/5;

   aaw0=0;
   for cod=Family_Phe_num_aminoacid
      aaw0=aaw0+aawdist(cod,spec);
   end
   AAW_Phe(spec)=aaw0/3;
end
bias_Ala_Ser=AAW_Ser-AAW_Ala;
bias_Glu_Asp=AAW_Asp-AAW_Glu;
bias_Ser_Phe=AAW_Phe-AAW_Ser;
axis_x=bias_Glu_Asp;
axis_y=bias_Ala_Ser;
axis_z=bias_Ser_Phe;

disp('Plotting Fig 3')
```

```
figure(3)
WIDTH=18;
LENGTH=18;
axis([0,WIDTH,0,LENGTH])
set (gcf,'Units','centimeters','Position',[0,0,WIDTH,LENGTH], 'color','w')
axis off
tlsubplot=tiledlayout(2,2);
tlsubplot.TileSpacing='Compact';
tlsubplot.Padding='Compact';

disp('Plotting Fig 3a')

% parameters
Para_MarkerSize=1;
Para_LineWidth=0.1;
Para_legendFontSize=6;
Para_FontSize=6;
Para_LabelSize=7;

nexttile

hold on
% 1:bacteria; 2:archaea; 3:fungi; 4:plants; 5:protozoa; 6:invertebrates; 7:vertebrates excluding mammals
    ↪  (vertebrates other); 8:mammals; 9:DNA viruses; 10:RNA virus
marker_taxa='..+*o^vs..';
color_taxa=[0 0 1
   1 0 0
   0 0.5 0.5
   0 1 0
   0.5 0.5 0.5
   0.5 0 0.5
   1 0 1
   1 0 0
   0 1 1
   0 1 0];
for i_taxon=Num_Taxa_Data:-1:1
   plot3(axis_x(NUM_TAXA{i_taxon}),axis_y(NUM_TAXA{i_taxon}),axis_z(NUM_TAXA{i_taxon}),marker_taxa(
       ↪ i_taxon),'color',color_taxa(i_taxon,:),'MarkerSize',Para_MarkerSize,'LineWidth',Para_LineWidth
       ↪ );
end

taxa_legend={'bacteria';'archaea';'fungi';'plants';'protozoa';'invertebrates';'vertebrates others';'
    ↪ mammals';'RNA viruses';'DNA viruses'};
Para_legend_x=-0.393/5+0.0008;
Para_legend_y=-0.28/3-0.001;
Para_legend_z=-0.195/3+0.062/3;
Para_legend_step=0.062/3;
for i_taxon=1:Num_Taxa_Data
   text(Para_legend_x,Para_legend_y,Para_legend_z+i_taxon*Para_legend_step,taxa_legend(i_taxon),'color',
       ↪ color_taxa(i_taxon,:),'FontSize',Para_legendFontSize);
end
Rotation_x=-13;
Rotation_y=73;
Rotation_z=90;
alx=xlabel('$\tilde{F}_{Glu}$-$\tilde{F}_{Asp}$ bias','Interpreter','Latex','FontSize',Para_LabelSize,'
    ↪ Rotation',Rotation_x,'Position',[-0.05,-0.64,-0.38]./[5 3 3]);
aly=ylabel('$\tilde{F}_{Ala}$-$\tilde{F}_{Ser}$ bias','Interpreter','Latex','FontSize',Para_LabelSize,'
    ↪ Rotation',Rotation_y,'Position',[0.205,-0.22,-0.4]./[5 3 3]);
alz=zlabel('$\tilde{F}_{Ser}$-$\tilde{F}_{Phe}$ bias','Interpreter','Latex','FontSize',Para_LabelSize,'
    ↪ Rotation',Rotation_z,'Position',[-0.33,-0.51,-0.3]./[5 3 3]);
xtickangle(Rotation_x)
ytickangle(Rotation_y)
ztickangle(Rotation_z)
axis([[-0.28,0.17]/5,[-0.595,0.1,-0.35,0.05]/3])
set(gca,'Linewidth',0.1,'GridAlpha',0.1,'Gridcolor',0.3*[1 1 1],'GridLineStyle','-')
set(gca,'XTick',[-0.2:0.1:0.1]/5)
set(gca,'XTickLabel',{' -0.04 ',' -0.02 ',' 0 ',' 0.02 '},'FontSize',Para_LabelSize)
set(gca,'YTick',[-0.45:0.15:0.1]/3)
set(gca,'YTickLabel',{' -0.15 ',' -0.10 ',' -0.05 ',' 0 '},'FontSize',Para_LabelSize)
set(gca,'ZTick',[-0.3:0.3:0]/3)
set(gca,'ZTickLabel',{' -0.1 ',' 0 '},'FontSize',Para_LabelSize)
set(gca,'Linewidth',0.1)
view(16.5,68)
```

```
legend(' ',' ',' ',' ',' ',' ',' ',' ',' ','box','off','Position',[0.02,0.8,0.01,0.169])
text(-0.3,-0.76,-0.35,'a','FontSize',9,'FontWeight','bold')
box on
hold off
text_fs=10;
title('a                    ','FontSize',text_fs,'FontWeight','bold','HorizontalAlignment','right','
    ↪ VerticalAlignment','top')

disp('Plotting Fig 3b')

nexttile

hold on
for i_taxon=3:Num_Taxa_Data-2
   plot3(axis_x(NUM_TAXA{i_taxon}),axis_y(NUM_TAXA{i_taxon}),axis_z(NUM_TAXA{i_taxon}),marker_taxa(
       ↪ i_taxon),'color',color_taxa(i_taxon,:),'MarkerSize',Para_MarkerSize,'LineWidth',Para_LineWidth
       ↪ );
end
taxa_legend_add={'bacteria'
   'archaea'
   'fungi'
   'plants'
   'protozoa'
   'invertebrates'
   'vertebrates others'
   'mammals'
   'A. thaliana'
   'C. elegans'
   'papaya'
   'Homo sapiens'};
color_add=[0 1 0
   0 1 1
   1 1 0
   0 0 0];
color_taxa_add=[color_taxa([1:8],:);color_add];
for i_taxon=9:12
   plot3(1,1,1,'.','color',color_taxa_add(i_taxon,:),'MarkerSize',0.2,'LineWidth',Para_LineWidth);
end
Para_legend_x=-0.05;
Para_legend_y=-0.065;
Para_legend_z=-0.1095;
Para_legend_step=-0.00205;
for i_taxon=3:12
   text(Para_legend_x,Para_legend_y,Para_legend_z+i_taxon*Para_legend_step,taxa_legend_add(i_taxon),'
       ↪ color',color_taxa_add(i_taxon,:),'FontSize',Para_legendFontSize);
end
bias_Athaliana=readmatrix('Supplementary_Data_4_ifaa_ifc_simulation.xlsx','sheet','Fig 3b','Range','A2:
    ↪ C10001');
bias_Celegans=readmatrix('Supplementary_Data_4_ifaa_ifc_simulation.xlsx','sheet','Fig 3b','Range','D2:
    ↪ F1001');
bias_Homosapiens=readmatrix('Supplementary_Data_4_ifaa_ifc_simulation.xlsx','sheet','Fig 3b','Range','G2
    ↪ :I1001');
bias_papaya=readmatrix('Supplementary_Data_4_ifaa_ifc_simulation.xlsx','sheet','Fig 3b','Range','J2:
    ↪ L1001');
bias_xyz=bias_Athaliana;
plot3(bias_xyz(:,1),bias_xyz(:,2),bias_xyz(:,3),'g.','MarkerSize',0.2);
bias_xyz=bias_Celegans;
plot3(bias_xyz(:,1),bias_xyz(:,2),bias_xyz(:,3),'c.','MarkerSize',0.2);
bias_xyz=bias_Homosapiens;
plot3(bias_xyz(:,1),bias_xyz(:,2),bias_xyz(:,3),'k.','MarkerSize',0.2);
bias_xyz=bias_papaya;
plot3(bias_xyz(:,1),bias_xyz(:,2),bias_xyz(:,3),'y.','MarkerSize',0.2);

legend(' ',' ',' ',' ',' ',' ',' ',' ',' ','box','off','FontSize',6,'Position'
    ↪ ,[0.533,0.435,0.06,0.125])
Rotation_x=-50;
Rotation_y=6;
Rotation_z=90;
alx=xlabel('$\tilde{F}_{Glu}$-$\tilde{F}_{Asp}$ bias','Interpreter','Latex','FontSize',Para_LabelSize,'
    ↪ Rotation',Rotation_x,'Position',[-0.041,-0.075,-0.1]);
aly=ylabel('$\tilde{F}_{Ala}$-$\tilde{F}_{Ser}$ bias','Interpreter','Latex','FontSize',Para_LabelSize,'
    ↪ Rotation',Rotation_y,'Position',[-0.005,-0.04,-0.1]);
```

```
alz=zlabel('$\tilde{F}_{Ser}$-$\tilde{F}_{Phe}$ bias','Interpreter','Latex','FontSize',Para_LabelSize,'Rotation',Rotation_z,'Position',[-0.05,-0.074,-0.07]);
xtickangle(Rotation_x)
ytickangle(Rotation_y)
ztickangle(Rotation_z)
axis([-0.05,-0.01,-0.07,-0.01,-0.1,-0.05])
set(gca,'Linewidth',0.1,'GridAlpha',0.1,'Gridcolor',0.3*[1 1 1],'GridLineStyle','-')
set(gca,'XTick',[-0.04:0.02:0])
set(gca,'XTickLabel',{' -0.04 ',' -0.02 ',' 0 '},'FontSize',Para_LabelSize)
set(gca,'YTick',[-0.06:0.02:0])
set(gca,'YTickLabel',{' -0.06 ',' -0.04 ',' -0.02 ',' 0 '},'FontSize',Para_LabelSize)
set(gca,'ZTick',[-0.1:0.02:-0.06])
set(gca,'ZTickLabel',{' -0.1 ',' -0.08 ',' -0.06 '},'FontSize',Para_LabelSize)
view(70.9,20.1)
box on
hold off
text_fs=10;
title('b                              ','FontSize',text_fs,'FontWeight','bold','HorizontalAlignment','right','VerticalAlignment','top')

disp('Plotting Fig 3c')

ifaa_table=readmatrix('Supplementary_Data_4_ifaa_ifc_simulation.xlsx','sheet','ifaa in proteome','Range','C2:BXZ3111');
temp_dist_1=ifaa_table';
temp_dist_2=reshape(temp_dist_1,100,20,3110);
ifaa=permute(temp_dist_2,[3,2,1]);
N_species=size(ifaa,1);
AAIFC_peak=zeros(1,N_species);
cutoff=100;
for i=1:N_species
   for j=1:20
      for k=2:cutoff-1
         if ifaa(i,j,k)>ifaa(i,j,k-1) && ifaa(i,j,k)>ifaa(i,j,k+1)
            AAIFC_peak(i)=AAIFC_peak(i)+1;
         end
      end
   end
end
proteome_size=zeros(1,N_species);
for i=1:N_species
   dist_species=reshape(ifaa(i,:,:),20,100);
   proteome_size(i)=sum(sum(dist_species,1),2);
end
log_proteome_size=log(proteome_size);

nexttile

hold on
Para_legend_x=5.4;
Para_legend_text_x=5.7;
Para_legend_y=380;
Para_legend_step=20;
for i_taxon=1:Num_Taxa_Data
   plot(Para_legend_x,Para_legend_y+Para_legend_step*i_taxon,marker_taxa(i_taxon),'color',color_taxa(i_taxon,:),'MarkerSize',Para_MarkerSize,'LineWidth',Para_LineWidth);
   text(Para_legend_text_x,Para_legend_y+Para_legend_step*i_taxon+1,taxa_legend(i_taxon),'color',color_taxa(i_taxon,:),'FontSize',Para_legendFontSize);
end
for i_taxon=1:Num_Taxa_Data
   plot(log_proteome_size(NUM_TAXA{i_taxon}),AAIFC_peak(NUM_TAXA{i_taxon}),marker_taxa(i_taxon),'color',color_taxa(i_taxon,:),'MarkerSize',Para_MarkerSize,'LineWidth',Para_LineWidth);
end

xlabel('log(proteome size)');
ylabel('if$\tilde{\mbox{c}}$ peaks','Interpreter','latex');
set(gca,'Linewidth',0.1)
axis([5,20,100,605])
set(gca,'XTick',[5:5:20])
set(gca,'XTickLabel',{'5','10','15','20'},'FontSize',Para_LabelSize)
set(gca,'YTick',[100:200:500])
set(gca,'YTickLabel',{'100','300','500'},'FontSize',Para_LabelSize)
Rotation_x=0;
```

```
Rotation_y=90;
xtickangle(Rotation_x)
ytickangle(Rotation_y)
set(gca,'Linewidth',0.1,'GridAlpha',0.1,'Gridcolor',0.3*[1 1 1],'GridLineStyle','-')
box on
hold off
text_fs=10;
title('c                              ','FontSize',text_fs,'FontWeight','bold','HorizontalAlignment','right','VerticalAlignment','top')

disp('Plotting Fig 3d')

nexttile

hold on
plot3(0,0,1,'c.','MarkerSize',3)
plot3(0,0,1,'b.','MarkerSize',3)
plot3(0,0,1,'y.','MarkerSize',3)
plot3(0,0,1,'r.','MarkerSize',3)
plot3(0,0,1,'g.','MarkerSize',3)
bias_V=readmatrix('Supplementary_Data_4_ifaa_ifc_simulation.xlsx','sheet','simulation Fig 3d','Range','A2:C12001');
bias_Bh=readmatrix('Supplementary_Data_4_ifaa_ifc_simulation.xlsx','sheet','simulation Fig 3d','Range','D2:F3001');
bias_Bl=readmatrix('Supplementary_Data_4_ifaa_ifc_simulation.xlsx','sheet','simulation Fig 3d','Range','G2:I3001');
bias_A=readmatrix('Supplementary_Data_4_ifaa_ifc_simulation.xlsx','sheet','simulation Fig 3d','Range','J2:L3001');
bias_E=readmatrix('Supplementary_Data_4_ifaa_ifc_simulation.xlsx','sheet','simulation Fig 3d','Range','M2:O3001');
bias_xyz=bias_V;
plot3(bias_xyz(:,1),bias_xyz(:,2),bias_xyz(:,3),'c.','MarkerSize',0.2);
bias_xyz=bias_Bh;
plot3(bias_xyz(:,1),bias_xyz(:,2),bias_xyz(:,3),'b.','MarkerSize',0.2);
bias_xyz=bias_Bl;
plot3(bias_xyz(:,1),bias_xyz(:,2),bias_xyz(:,3),'y.','MarkerSize',0.2);
bias_xyz=bias_A;
plot3(bias_xyz(:,1),bias_xyz(:,2),bias_xyz(:,3),'r.','MarkerSize',0.2);
bias_xyz=bias_E;
plot3(bias_xyz(:,1),bias_xyz(:,2),bias_xyz(:,3),'g.','MarkerSize',0.2);

legend('low bias','$\tilde{F}_{Ala}$ rich','$\tilde{F}_{Phe}$ rich','$\tilde{F}_{Ser}$ rich','$\tilde{F}_{Asp}$ rich','Interpreter','Latex','box','off','FontSize',6,'Position',[0.58,0.19,0.05,0.11])
view(-108.97,69.85)
Rotation_x=-70;
Rotation_y=15;
Rotation_z=94;
alx=xlabel('$\tilde{F}_{Glu}$-$\tilde{F}_{Asp}$ bias','Interpreter','Latex','FontSize',Para_LabelSize,'Rotation',Rotation_x,'Position',[0.03,0.13,-0.15]);
aly=ylabel('$\tilde{F}_{Ala}$-$\tilde{F}_{Ser}$ bias','Interpreter','Latex','FontSize',Para_LabelSize,'Rotation',Rotation_y,'Position',[-0.075,-0.05,-0.15]);
alz=zlabel('$\tilde{F}_{Ser}$-$\tilde{F}_{Phe}$ bias','Interpreter','Latex','FontSize',Para_LabelSize,'Rotation',Rotation_z,'Position',[0.12,0.12,-0.05]);
xtickangle(Rotation_x)
ytickangle(Rotation_y)
ztickangle(Rotation_z)
axis([-0.06,0.12,-0.2,0.1,-0.15,0.05])
set(gca,'Linewidth',0.1,'GridAlpha',0.1,'Gridcolor',0.3*[1 1 1],'GridLineStyle','-')
set(gca,'XTick',[-0.05:0.05:0.1])
set(gca,'XTickLabel',{'-0.05',' 0 ',' 0.05 ',' 0.1 '},'FontSize',Para_LabelSize)
set(gca,'YTick',[-0.15:0.1:0.05])
set(gca,'YTickLabel',{'-0.15','-0.05','0.05'},'FontSize',Para_LabelSize)
set(gca,'ZTick',[-0.15:0.2:0.05])
set(gca,'ZTickLabel',{' -0.15 ','0.05 '},'FontSize',Para_LabelSize)
box on
hold off
text_fs=10;
title('d  ','FontSize',text_fs,'FontWeight','bold','HorizontalAlignment','right','VerticalAlignment','top')

disp('Saving figures')
n='3';
saveas(gcf,['Fig_',n,'.fig']);
```

```
exportgraphics(gcf,['Fig_',n,'.pdf'],'Resolution',1500)
exportgraphics(gcf,['Fig_',n,'.tif'],'Resolution',1500)
exportgraphics(gcf,['Fig_',n,'.png'],'Resolution',1500)
exportgraphics(gcf,['Fig_',n,'_r1500.eps'],'Resolution',1500)

disp('Finished')
```

### MATLAB script to generate Fig. 4

```
% Supplementary program to Plotting Fig. 4
% MATLAB_R2024a

clear;clc;

disp('Plotting Fig 4')

figure(4)
WIDTH=18;
LENGTH=18;
hold on
axis([0,WIDTH,0,LENGTH])
set (gcf,'Units','centimeters','Position',[0,0,WIDTH,LENGTH], 'color','w')
axis off
hold off
tlsubplot=tiledlayout(3,3);
tlsubplot.TileSpacing='Compact';
tlsubplot.Padding='Compact';

disp('Plotting Fig 4a')

clear;
nexttile

fs=6; fontsizefortext=5;
PARA_x=50;
PARA_y=0.0405;
simulatedifc_4a=readmatrix('Supplementary_Data_4_ifaa_ifc_simulation.xlsx','sheet','simulation Fig 4','
    ↪ Range','A1:ALL64');
dist0=simulatedifc_4a;
dist1=reshape(dist0,[1,size(dist0,1)*size(dist0,2)]);
total=sqrt(dist1*dist1');
dist2=dist0./total;dist=dist2(:,[2:end]);
distaverage_Ath(7,:)=mean(dist(:,[1:249]),1);
hold on
plot(dist','LineWidth',0.1)
axis([0,100,0,0.048])
text(PARA_x,PARA_y,'periods 2-5 base','HorizontalAlignment','center','FontSize',fontsizefortext,'color',
    ↪ 'b')
set(gca,'XTick',0:3:100)
set(gca,'XTickLabel',{'0',' ',' ','9',' ',' ','18',' ',' ','27',' ',' ','36',' ',' ','45',' ',' ','54','
    ↪ ',' ','63',' ',' ','72',' ',' ','81',' ',' ','90',' ',' ','99'},'FontSize',fs)
set(gca,'YTick',[0:0.01:0.04])
set(gca,'YTickLabel',{'0','0.01','0.02','0.03','0.04'},'FontSize',fs)
xlabel('interval between 3-tuples (base)','FontSize',fs)
ylabel('simulated ifc^3','FontSize',fs)
set(gca,'Xgrid','on')
set(gca,'Linewidth',0.1,'GridAlpha',0.1,'Gridcolor',0.3*[1 1 1],'GridLineStyle','-')
box on
hold off
text_fs=9;
text_x=-10*0.81;
text_y=0;
text(text_x,text_y,'a','FontSize',text_fs,'FontWeight','bold','HorizontalAlignment','center')

disp('Plotting Fig 4b')

clear;
```

```
nexttile

simulatedifc_4b=readmatrix('Supplementary_Data_4_ifaa_ifc_simulation.xlsx','sheet','simulation Fig 4','
    ↪ Range','A65:IO69');
fs=6; fontsizefortext=5;
hold on
plot(simulatedifc_4b','LineWidth',0.1)
axis([0,100,3e-5,0.2])
text(36,5e-2,'periods 2-5 base','HorizontalAlignment','center','FontSize',fontsizefortext,'color','b')
set(gca,'XTick',0:3:100,'FontSize',fs)
set(gca,'XTickLabel',{'0',' ',' ','9',' ',' ','18',' ',' ','27',' ',' ','36',' ',' ','45',' ',' ','54','
    ↪  ',' ','63',' ',' ','72',' ',' ','81',' ',' ','90',' ',' ','99'},'FontSize',fs)
xlabel('interval between n-tuples (base)','FontSize',fs)
ylabel('simulated IFC^n','FontSize',fs,'FontSize',fs)
set(gca,'yscale','log')
legend('','','','','','location','Northeast')
legend('IFC^2','IFC^3','IFC^4','IFC^5','IFC^6','location','NorthEast','box','off','FontSize',5,'Position
    ↪ ',[0.555,0.88,0.07,0.07])
set(gca,'Xgrid','on')
box on
set(gca,'Linewidth',0.1,'GridAlpha',0.1,'Gridcolor',0.3*[1 1 1],'GridLineStyle','-')
hold off
text_fs=9;
text_x=-10*0.81;
text_y=3e-5;
text(text_x,text_y,'b','FontSize',text_fs,'FontWeight','bold','HorizontalAlignment','center')

disp('Plotting Fig 4c')

clear;
nexttile

simulatedifc_4c=readmatrix('Supplementary_Data_4_ifaa_ifc_simulation.xlsx','sheet','simulation Fig 4','
    ↪ Range','A70:IO74');
fs=6; fontsizefortext=5;
PARA_y=3e-4;
hold on
plot(simulatedifc_4c','LineWidth',0.1)
axis([0,100,3e-5,0.2])
text(36,5e-2,'periods 2-10 base','HorizontalAlignment','center','FontSize',fontsizefortext,'color','g')
set(gca,'XTick',0:3:100,'FontSize',fs)
set(gca,'XTickLabel',{'0',' ',' ','9',' ',' ','18',' ',' ','27',' ',' ','36',' ',' ','45',' ',' ','54','
    ↪  ',' ','63',' ',' ','72',' ',' ','81',' ',' ','90',' ',' ','99'},'FontSize',fs)
xlabel('interval between n-tuples (base)','FontSize',fs)
ylabel('simulated IFC^n','FontSize',fs,'FontSize',fs)
set(gca,'yscale','log')
legend('','','','','','location','Northeast')
legend('IFC^2','IFC^3','IFC^4','IFC^5','IFC^6','location','NorthEast','box','off','FontSize',5,'Position
    ↪ ',[0.87,0.88,0.07,0.07])
set(gca,'Xgrid','on')
box on
set(gca,'Linewidth',0.1,'GridAlpha',0.1,'Gridcolor',0.3*[1 1 1],'GridLineStyle','-')
hold off
text_fs=9;
text_x=-10*0.81;
text_y=3e-5;
text(text_x,text_y,'c','FontSize',text_fs,'FontWeight','bold','HorizontalAlignment','center')

disp('Plotting Fig 4d')

clear;
nexttile

simulatedifc_4d=readmatrix('Supplementary_Data_4_ifaa_ifc_simulation.xlsx','sheet','simulation Fig 4','
    ↪ Range','A75:ALL138');
fs=6; fontsizefortext=5;
PARA_x=50;
PARA_y=0.0405;
dist0=simulatedifc_4d;
dist1=reshape(dist0,[1,size(dist0,1)*size(dist0,2)]);
total=sqrt(dist1*dist1');
dist2=dist0./total;dist=dist2(:,[2:end]);
distaverage_Ath(7,:)=mean(dist(:,[1:249]),1);
```

```
hold on
plot(dist','LineWidth',0.1)
axis([0,100,0,0.048])
text(PARA_x,PARA_y,'period 3 base','HorizontalAlignment','center','FontSize',fontsizefortext,'color','r'
    ↪ )
set(gca,'XTick',0:3:100)
set(gca,'XTickLabel',{'0',' ',' ','9',' ',' ','18',' ',' ','27',' ',' ','36',' ',' ','45',' ',' ','54','
    ↪  ',' ','63',' ',' ','72',' ',' ','81',' ',' ','90',' ',' ','99'},'FontSize',fs)
set(gca,'YTick',[0:0.01:0.04])
set(gca,'YTickLabel',{'0','0.01','0.02','0.03','0.04'},'FontSize',fs)
xlabel('interval between 3-tuples (base)','FontSize',fs)
ylabel('simulated ifc^3','FontSize',fs)
set(gca,'Xgrid','on')
set(gca,'Linewidth',0.1,'GridAlpha',0.1,'Gridcolor',0.3*[1 1 1],'GridLineStyle','-')
box on
hold off
text_fs=9;
text_x=-10*0.81;
text_y=0;
text(text_x,text_y,'d','FontSize',text_fs,'FontWeight','bold','HorizontalAlignment','center')

disp('Plotting Fig 4e')

clear;
nexttile

simulatedifc_4e=readmatrix('Supplementary_Data_4_ifaa_ifc_simulation.xlsx','sheet','simulation Fig 4','
    ↪ Range','A139:ALL202');
dist0=simulatedifc_4e;
dist0_reshape=reshape(dist0,[1 64000]);
ifc=dist0/sqrt(dist0_reshape*dist0_reshape');
fft_ifc=fft(ifc')';
PS_ifc=fft_ifc.*conj(fft_ifc);
fs=6; fontsizefortext=5;
hold on
plot(PS_ifc','LineWidth',0.1)
axis([0,501,2*10^-10,3])
text(38,1*10^-8,'periods 2-5 base','HorizontalAlignment','left','FontSize',fontsizefortext,'color','b')
text(38,3*10^-9,'power spectrum of simulated ifc^3','HorizontalAlignment','left','FontSize',
    ↪ fontsizefortext)
set(gca,'yscale','log')
set(gca,'XTick',[0,1/5,1/4,1/3,2/5,1/2]*1000)
set(gca,'XTickLabel',{'0','1/5','1/4','1/3','2/5','1/2'},'FontSize',fs)
set(gca,'YTick',[10^-8,10^-6,10^-4,10^-2,1])
set(gca,'YTickLabel',{'10^{-8}','10^{-6}','10^{-4}','10^{-2}','10^0'},'FontSize',fs)
xlabel('frequency (1/base)','FontSize',fs)
ylabel('simulated power spectrum','FontSize',fs)
box on
set(gca,'Xgrid','on')
set(gca,'Linewidth',0.1,'GridAlpha',0.1,'Gridcolor',0.0*[1 1 1],'GridLineStyle','-')
hold off
text_fs=9;
text_x=-50*0.81;
text_y=2*10^-10;
text(text_x,text_y,'e','FontSize',text_fs,'FontWeight','bold','HorizontalAlignment','center')

disp('Plotting Fig 4f')

clear;
nexttile

simulatedifc_4f=readmatrix('Supplementary_Data_4_ifaa_ifc_simulation.xlsx','sheet','simulation Fig 4','
    ↪ Range','A203:ALL266');
dist0=simulatedifc_4f;
dist0_reshape=reshape(dist0,[1 64000]);
ifc=dist0/sqrt(dist0_reshape*dist0_reshape');
fft_ifc=fft(ifc')';
PS_ifc=fft_ifc.*conj(fft_ifc);
fs=6; fontsizefortext=5;
hold on
plot(PS_ifc','LineWidth',0.1)
axis([0,501,2*10^-10,3])
text(38,1*10^-8,'periods 2-10 base','HorizontalAlignment','left','FontSize',fontsizefortext,'color','g')
```

```
text(38,3*10^-9,'power spectrum of simulated ifc^3','HorizontalAlignment','left','FontSize',
    ↪ fontsizefortext)
set(gca,'yscale','log')
set(gca,'XTick',[0,1/10,1/9,1/8,1/7,1/6,1/5,2/9,1/4,2/7,3/10,1/3,3/8,2/5,3/7,4/9,1/2]*1000)
set(gca,'XTickLabel',{'0',' 1/10','1/9','1/8','1/7','1/6','1/5(2/10)','2/9','1/4(2/8)','2/7','3/10','
    ↪ 1/3(2/6,3/9)','3/8','2/5(4/10)','3/7','4/9','1/2'},'FontName','Arial Narrow','FontSize',fs-2)
xtickangle(90)
xlabel('frequency (1/base)','FontSize',fs)
ylabel('simulated power spectrum','FontSize',fs)
box on
set(gca,'Xgrid','on')
set(gca,'Linewidth',0.1,'GridAlpha',0.1,'Gridcolor',0.0*[1 1 1],'GridLineStyle','-')
hold off
text_fs=9;
text_x=-50*0.81;
text_y=2*10^-10;
text(text_x,text_y,'f','FontSize',text_fs,'FontWeight','bold','HorizontalAlignment','center')

disp('Plotting Fig 4g')

clear;
nexttile

simulatedifc_4g=readmatrix('Supplementary_Data_4_ifaa_ifc_simulation.xlsx','sheet','simulation Fig 4','
    ↪ Range','A267:ALL330');
dist0=simulatedifc_4g;
dist0_reshape=reshape(dist0,[1 64000]);
ifc=dist0/sqrt(dist0_reshape*dist0_reshape');
fft_ifc=fft(ifc')';
PS_ifc=fft_ifc.*conj(fft_ifc);
fs=6; fontsizefortext=5;
hold on
plot(PS_ifc','LineWidth',0.1)
axis([0,501,8*10^-11,3])
text(38,1.6*10^-9,'period 3 base','HorizontalAlignment','left','FontSize',fontsizefortext,'color','r')
text(38,3*10^-10,'power spectrum of simulated ifc^3','HorizontalAlignment','left','FontSize',
    ↪ fontsizefortext)
set(gca,'yscale','log')
set(gca,'XTick',[0,1/3,1/2]*1000)
set(gca,'XTickLabel',{'0','1/3','1/2'},'FontSize',fs)
set(gca,'YTick',[10^-8,10^-6,10^-4,10^-2,1])
set(gca,'YTickLabel',{'10^{-8}','10^{-6}','10^{-4}','10^{-2}','10^0'},'FontSize',fs)
xlabel('frequency (1/base)','FontSize',fs)
ylabel('simulated power spectrum','FontSize',fs)
box on
set(gca,'Xgrid','on')
set(gca,'Linewidth',0.1,'GridAlpha',0.1,'Gridcolor',0.0*[1 1 1],'GridLineStyle','-')
hold off
text_fs=9;
text_x=-50*0.81;
text_y=2*10^-10;
text(text_x,text_y,'g','FontSize',text_fs,'FontWeight','bold','HorizontalAlignment','center')

disp('Plotting Fig 4h')

clear;
nexttile

simulatedifc_4h=readmatrix('Supplementary_Data_4_ifaa_ifc_simulation.xlsx','sheet','ifc(Haloarcula
    ↪ hispanica N601)','Range','A1:ALL64');
dist0=simulatedifc_4h;
dist0_reshape=reshape(dist0,[1 64000]);
ifc=dist0/sqrt(dist0_reshape*dist0_reshape');
fft_ifc=fft(ifc')';
PS_ifc=fft_ifc.*conj(fft_ifc);
fs=6; fontsizefortext=5;
hold on
plot(PS_ifc','LineWidth',0.1)
axis([0,501,8*10^-11,3])
text(15,1.6*10^-9,'an archaeal genome with a high coding ratio','HorizontalAlignment','left','FontSize',
    ↪ fontsizefortext,'color','r')
text(15,3*10^-10,'power spectrum of ifc^3(Haloarcula hispanica N601)','HorizontalAlignment','left','
    ↪ FontSize',fontsizefortext)
```

```
set(gca,'yscale','log')
set(gca,'XTick',[0,1/3,1/2]*1000)
set(gca,'XTickLabel',{'0','1/3','1/2'},'FontSize',fs)
set(gca,'YTick',[10^-8,10^-6,10^-4,10^-2,1])
set(gca,'YTickLabel',{'10^{-8}','10^{-6}','10^{-4}','10^{-2}','10^0'},'FontSize',fs)
xlabel('frequency (1/base)','FontSize',fs)
ylabel('power spectrum','FontSize',fs)
box on
set(gca,'Xgrid','on')
set(gca,'Linewidth',0.1,'GridAlpha',0.1,'Gridcolor',0.0*[1 1 1],'GridLineStyle','-')
hold off
text_fs=9;
text_x=-50*0.81;
text_y=2*10^-10;
text(text_x,text_y,'h','FontSize',text_fs,'FontWeight','bold','HorizontalAlignment','center')

disp('Plotting Fig 4i')

clear;
nexttile

simulatedifc_4i=readmatrix('Supplementary_Data_4_ifaa_ifc_simulation.xlsx','sheet','ifc(Gallus gallus)',
    ↪ 'Range','A1:ALL64');
fs=6; fontsizefortext=5;
ifc0=simulatedifc_4i;
ifc_reshape=reshape(ifc0,[1 64000]);
ifc=ifc0/sqrt(ifc_reshape*ifc_reshape');
fft_ifc=fft(ifc')';
PS_ifc=fft_ifc.*conj(fft_ifc);
hold on
plot(PS_ifc','LineWidth',0.1)
axis([0,501,9^-12,3])
text(38,1*10^-10,'a eukaryotic genome with a low coding ratio','HorizontalAlignment','left','FontSize',
    ↪ fontsizefortext,'color','g')
text(38,2*10^-11,'power spectrum of ifc^3(Gallus gallus)','HorizontalAlignment','left','FontSize',
    ↪ fontsizefortext)
set(gca,'yscale','log')
set(gca,'XTick',[0,1/10,1/9,1/8,1/7,1/6,1/5,2/9,1/4,2/7,3/10,1/3,3/8,2/5,3/7,4/9,1/2]*1000)
set(gca,'XTickLabel',{'0','1/10','1/9','1/8','1/7','1/6','1/5','2/9','1/4','2/7','3/10','1/3','3/8','2/5
    ↪ ','3/7','4/9','1/2'},'FontName','Arial Narrow','FontSize',fs-2)
xtickangle(90)
xlabel('frequency (1/base)','FontSize',fs)
ylabel('power spectrum','FontSize',fs)
box on
set(gca,'Xgrid','on')
set(gca,'Linewidth',0.1,'GridAlpha',0.1,'Gridcolor',0.0*[1 1 1],'GridLineStyle','-')
hold off
text_fs=9;
text_x=-50*0.81;
text_y=9^-12;
text(text_x,text_y,'i','FontSize',text_fs,'FontWeight','bold','HorizontalAlignment','center')

disp('Saving figures')
saveas(gcf,'Fig_4.fig');
exportgraphics(gcf,'Fig_4.pdf','Resolution',1500)
exportgraphics(gcf,'Fig_4.tif','Resolution',1500)
exportgraphics(gcf,'Fig_4.eps','Resolution',1500)
exportgraphics(gcf,'Fig_4.png','Resolution',1500)

disp('Finished')
```

### MATLAB script to simulate genomes in Fig. 1d and proteomes in Fig. 3d

```
% Supplementary program to simulate genomes in Fig. 1d and proteomes in Fig. 3d
% MATLAB_R2024a

clear; clc;
```

```
PlotFigures = 1;  % 2 strands, 6 reading frames, Fig. 1d and Fig. 3d;
% PlotFigures = 2; % 1 strand, 1 reading frame, Supp. Fig. 1dA and Supp. Fig. 3dA;
% PlotFigures = 3; % 2 strands, 2 reading frames, Supp. Fig. 1dB;
% PlotFigures = 4; % 1 strand, 3 reading frames, Supp. Fig. 3dB;

% KEY IDEA: specific amino acid biosynthetic families with a dominant probability of occurrence; F_Ala,
    ↪ F_Ser, F_Glu, F_Asp, F_Phe, others
ParameterSuppTab1_VgVlBhBlAE=[1 1 4 1 1 0.2 % viruses(F_Glu-rich) F_Glu rich
1 2 0.2 3 6 0.2                    % viruses(low bias): no family rich
4 1 0.1 1 1 0.2                    % bacteria(high GC): F_Ala rich
1 1 0.1 1 6 0.2                    % bacteria(low GC): F_Phe rich
1 4 0.1 1 1 0.2                    % archaea: F_Ser rich
1 1 0.1 4 1 0.2];                  % eukaryotes: F_Asp rich

codon=[];
for i=1:4
   for j=1:4
      for k=1:4
         codon=[codon;[i,j,k]]; % the 64 codons; '1', '2', '3', '4' stand for 'A', 'C', 'G', 'U'/'T',
             ↪ respectively
      end
   end
end

complementarystrand=[4 3 2 1]; % A to T, G to C, C to G, T to A

%=========================================================
% The amino acid biosynthetic familes in textbooks
codons_Ala=[30 32 31 29 63 61 45 46 47 48 37 38 39 40]; %14
codons_Ser=[53 54 55 56 10 12 58 60 41 42 43 44]; %12
codons_Glu=[22 11 24 35 9 33 23 21 26 28 27 19 25 17 1 3]; %16
codons_Asp=[34 2 36 4 7 5 15 13 6 14 8 16]; %12
codons_Phe=[62 64 50 52 59]; %5
codons_others=[18 20 49 51 57]; %5
%=========================================================
codons_AAFamily={codons_Ala;codons_Ser;codons_Glu;codons_Asp;codons_Phe;codons_others};
codons_fam0=[14 12 16 12 5 5];

TheGeneticCode='KNKNTTTTRSRSIIMIQHQHPPPPRRRRLLLLEDEDAAAAGGGGVVVVZYZYSSSSZCWCLFLF'; % Z: stop codons

NumGenPro0=1000; % input this number
NumGenPro=3*NumGenPro0;
SimulatedGenomeNumber_VgVlBhBlAE=[NumGenPro 4*NumGenPro NumGenPro NumGenPro NumGenPro NumGenPro]; %
    ↪ numbers of simulated genomes/proteomes

cutoff=1000;
choice_num=19;
n1_c=500;

CW_taxon_all=[];
position_ProEvoSp_all=[];
ifaa_normal_all=[];

for dom_VgVlBhBlAE_6InTotal=1:6 % viruses(F_Glu-rich), viruses(low bias), bacteria(high GC), bacteria(
    ↪ low GC), archaea, eukaryotes

   dom_VgVlBhBlAE_6InTotal=dom_VgVlBhBlAE_6InTotal % Display Progress

   N_SimulatedGenPro=SimulatedGenomeNumber_VgVlBhBlAE(dom_VgVlBhBlAE_6InTotal); % number of simulated
       ↪ genomes
   upweight=ParameterSuppTab1_VgVlBhBlAE(dom_VgVlBhBlAE_6InTotal,:); % Key parameter

   codons_fam=codons_fam0.*upweight; % Upweight specific amino acid biosynthetic family for current loop
   codons_fam_probility=codons_fam/sum(codons_fam);
   codons_fam_probility_step=zeros(1,7);
   codons_fam_probility_step(1)=0;
   for i=1:6
      codons_fam_probility_step(i+1)=codons_fam_probility_step(i)+codons_fam_probility(i);
   end

   CW_taxon=zeros(64,1);
   position_ProEvoSp_taxon=zeros(3,1);
```

```
distribution_20AA_normal=zeros(20,cutoff);
ifaa_normal=zeros(N_SimulatedGenPro,20,cutoff);

for N_GenomeProteome=1:N_SimulatedGenPro % number of simulated genomes

   codon_upweight=[];
   for i=1:choice_num
      fam_choice=1;
      rand_fam=rand(1);
      for j=1:6
         if rand_fam>codons_fam_probility_step(j) && rand_fam<codons_fam_probility_step(j+1)
            fam_choice=j;
         end
      end
      codon_upweight=[codon_upweight,codons_AAFamily{fam_choice}];
   end

   % ----------------------------------------------
   % Part I (Fig. 1d): Generating simulated genomes
   % ----------------------------------------------

   size_codon_upweight=size(codon_upweight,2);
   n2_c=ceil(size_codon_upweight/4);
   selectedcodons0=zeros(n1_c,n2_c);
   for i=1:n1_c
      selectedcodons0(i,:)=randperm(size_codon_upweight,n2_c); % selecting codons from proper subsets
          ↪  of the 64 codons to achieve three-base periodicity
   end
   n1n2=size(selectedcodons0,1)*size(selectedcodons0,2);
   selectedcodons0_t=selectedcodons0';
   selectedcodons=reshape(selectedcodons0_t,[1 n1n2]);

   g0=zeros(n1n2,3);
   for i=1:n1n2
      g0(i,:)=codon(codon_upweight(selectedcodons(i)),:);
   end
   g1=g0';
   genome0=reshape(g1,[1 3*n1n2]); % simulated genome

   switch PlotFigures
      case 1 % 2 strands, 6 reading frames, Fig. 1d and Fig. 3d
         genome_reverse=genome0(end:-1:1);
         genome_antisense=complementarystrand(genome_reverse); % antisense strand
         genome=[genome0,genome_antisense]; % double strands
      case 2 % 1 strand, 1 reading frame, Supp. Fig. 1dA and Supp. Fig. 3dA
         genome=genome0; % single strand
      case 3 % 2 strands, 2 reading frames, Supp. Fig. 1dB;
         genome_reverse=genome0(end:-1:1);
         genome_antisense=complementarystrand(genome_reverse); % antisense strand
         genome=[genome0,genome_antisense]; % double strands
      case 4 % 1 strand, 3 reading frames, Supp. Fig. 3dB;
         genome=genome0; % single strand
   end

   % Calculating ifc for the simulated genome

   h=genome; % calculating ifc for a simulated genome
   h_part_size=floor(size(h,2)/5);
   ifc=[];
   for N_codon=1:64
      interval0=[];
      for i_part=1:4
         interval0_part=[];
         for i=(i_part-1)*h_part_size+1:i_part*h_part_size
            if h(i)==codon(N_codon,1) && h(i+1)==codon(N_codon,2) && h(i+2)==codon(N_codon,3)
               interval0_part=[interval0_part i];
            end
         end
         interval0=[interval0 interval0_part];
      end
      for i=4*h_part_size+1:(size(h,2)-3)
         if h(i)==codon(N_codon,1) && h(i+1)==codon(N_codon,2) && h(i+2)==codon(N_codon,3)
            interval0=[interval0 i];
```

```
        end
    end
    interval=interval0(2:end)-interval0(1:end-1);

    distribution=zeros(1,cutoff);
    size_interval=size(interval,2);
    for i=1:size_interval
        interval_i=int32(interval(i));
        if interval_i≥1 && interval_i≤cutoff
            distribution(interval_i)=distribution(interval_i)+1;
        end
    end
    ifc=[ifc; distribution];
end

distdist=ifc.*ifc;
sumdistdist=sum(distdist,2);
sumsumdistdist=sum(sumdistdist,1);
CW=sumdistdist/sumsumdistdist; % codon weight for the simulated genome
CW_taxon(:,N_GenomeProteome)=CW; % codon weights for the N_GenomeProteome genomes

% -----------------------------------------------------------------------------
% Part II (Fig. 3d): translation from simulated genomes to simulated proteomes
% -----------------------------------------------------------------------------

switch PlotFigures
    case 1 % 2 strands, 6 reading frames, Fig. 1d and Fig. 3d
        genome00=[genome,genome(2:end-2),genome(3:end-1)]; % 3 reading frames, N_GenomeProteome is selected as a multiple of three
        genome=genome00;
    case 2 % 1 strand, 1 reading frame, Supp. Fig. 1dA and Supp. Fig. 3dA
        genome=genome;
    case 3 % 2 strands, 2 reading frames, Supp. Fig. 1dB;
        genome=genome;
    case 4 % 1 strand, 3 reading frames, Supp. Fig. 3dB;
        genome00=[genome,genome(2:end-2),genome(3:end-1)]; % 3 reading frames, N_GenomeProteome is selected as a multiple of three
        genome=genome00;
end

genomesize=size(genome,2);
proteomesize=floor(genomesize/3);
proteome=blanks(proteomesize);
for i=1:proteomesize
    codon_num=16*(genome(3*i-2)-1)+4*(genome(3*i-1)-1)+genome(3*i);
    proteome(i)=TheGeneticCode(codon_num); % translation from the simulated genome to simulated proteome based on the standard genetic code table
end

% Calculating if\tilde{c}, namely normalized ifaa, for the simulated proteome

L=strlength(proteome);
position=zeros(26,L);
for i=1:L
    k=proteome(i)-'A'+1;
    position(k,i)=1;
end

positioin_old=0;
counter20AA=0;
distribution_20AA=zeros(20,cutoff);
for n_AA=[1,3,4,5,6,7,8,9,11,12,13,14,16,17,18,19,20,22,23,25] % the 20 amino acids out of the 26 letters
    for i=1:L
        if position(n_AA,i)==1
            positioin_old=i;
            break
        end
    end

    n=0;
    interval=zeros(1,L);
    for i=positioin_old+1:L
```

```
            if position(n_AA,i)==1
               n=n+1;
               interval(n)=i-positioin_old;
               positioin_old=i;
            end
         end

         distribution=zeros(1,cutoff);
         size_interval=size(interval,2);
         for i=1:size_interval
            interval_i=int32(interval(i));
            if interval_i≥1 && interval_i≤cutoff
               distribution(interval_i)=distribution(interval_i)+1;
            end
         end

         counter20AA=counter20AA+1;
         distribution_20AA(counter20AA,:)=distribution;
      end

      dist1=reshape(distribution_20AA,[1,size(distribution_20AA,1)*size(distribution_20AA,2)]);
      total=sqrt(dist1*dist1');
      ifaa_normal(N_GenomeProteome,:,:)=distribution_20AA/total;
   end

   CW_taxon_all=[CW_taxon_all,CW_taxon];
   ifaa_normal_all=[ifaa_normal_all;ifaa_normal];
end

% Calculating the coordinates in GenEvoSp and ProEvoSp

% Groups of codons in the amino acid biosynthetic familes
num_Glu=[22 11 24 35 9 33 23 21 26 28 27 19 25 17 1 3]; %16
num_Asp=[34 2 36 4 7 5 15 13 6 14 8 16]; %12
num_Ala=[30 32 31 29 63 61 45 46 47 48 37 38 39 40]; %14
num_Ser=[53 54 55 56 10 12 58 60 41 42 43 44]; %12
num_Phe=[62 64 50 52 59]; %5

cwdist=CW_taxon_all;
N_species=size(CW_taxon_all,2);
for spec=1:N_species
   cw0=0;
   for cod=num_Glu
      cw0=cw0+cwdist(cod,spec);
   end
   cwdist_Glu=cw0/16;
   cwdist_Glu_all(spec)=cw0/16;
   cw0=0;
   for cod=num_Asp
      cw0=cw0+cwdist(cod,spec);
   end
   cwdist_Asp=cw0/12;
   cwdist_Asp_all(spec)=cw0/12;

   cw0=0;
   for cod=num_Ala
      cw0=cw0+cwdist(cod,spec);
   end
   cwdist_Ala=cw0/14;
   cwdist_Ala_all(spec)=cw0/14;
   cw0=0;
   for cod=num_Ser
      cw0=cw0+cwdist(cod,spec);
   end
   cwdist_Ser=cw0/12;
   cwdist_Ser_all(spec)=cw0/12;
   cw0=0;
   for cod=num_Phe
      cw0=cw0+cwdist(cod,spec);
   end
   cwdist_Phe=cw0/5;
   cwdist_Phe_all(spec)=cw0/5;
end
```

```
xx=cwdist_Asp_all-cwdist_Glu_all;
yy=cwdist_Ser_all-cwdist_Ala_all;
zz=cwdist_Phe_all-cwdist_Ser_all;
position_GenEvoSp_all=[xx;yy;zz];

% Groups of amino acids in the amino acid biosynthetic familes
Family_Val=[18 1 10];
Family_Ser=[16 6 2];
Family_Glu=[4 13 15 14 9];
Family_Asp=[3 17 8 11 12];
Family_Phe=[5 19 20];
Family_His=[7];

Family_VSGAPH={Family_Val;Family_Ser;Family_Glu;Family_Asp;Family_Phe;Family_His};

N_species=size(ifaa_normal_all,1);
N_AA=size(ifaa_normal_all,2);
distdist=sum(ifaa_normal_all.^2,3);
AAW=[];
for f=1:1
   Family_any=Family_VSGAPH{f,:};
   distdist_Family=distdist(:,Family_any);
   AAW=sum(distdist_Family,2); % Amino Acid Weight
end
for f=2:6
   Family_any=Family_VSGAPH{f,:};
   distdist_Family=distdist(:,Family_any);
   AAW=[AAW,sum(distdist_Family,2)]; % Amino Acid Weight
end
num_Family=[3 3 5 5 3 1];
for f=1:6
   AAW(:,f)=AAW(:,f)/num_Family(f);
end

Family_Phe_Family_Ser=AAW(:,5)-AAW(:,2);
Family_Asp_Family_Glu=AAW(:,4)-AAW(:,3);
Family_Ser_Family_Val=AAW(:,2)-AAW(:,1);
x=Family_Asp_Family_Glu';
y=Family_Ser_Family_Val';
z=Family_Phe_Family_Ser';
position_ProEvoSp_all=[x;y;z];

disp('Saving the simulated genome and proteome results in Supplementary_Data_4_ifaa_ifc_simulation.xlsx'
    ↪ )
position_GenEvoSp_Vl=position_GenEvoSp_all(:,NumGenPro+1:5*NumGenPro);
position_GenEvoSp_Bh=position_GenEvoSp_all(:,5*NumGenPro+1:6*NumGenPro);
position_GenEvoSp_Bl=position_GenEvoSp_all(:,6*NumGenPro+1:7*NumGenPro);
position_GenEvoSp_A=position_GenEvoSp_all(:,7*NumGenPro+1:8*NumGenPro);
position_GenEvoSp_E=position_GenEvoSp_all(:,8*NumGenPro+1:9*NumGenPro);
writematrix(position_GenEvoSp_Vl','Supplementary_Data_4_ifaa_ifc_simulation.xlsx','sheet','simulation
    ↪ Fig 1d','Range','A2')
writematrix(position_GenEvoSp_Bh','Supplementary_Data_4_ifaa_ifc_simulation.xlsx','sheet','simulation
    ↪ Fig 1d','Range','D2')
writematrix(position_GenEvoSp_Bl','Supplementary_Data_4_ifaa_ifc_simulation.xlsx','sheet','simulation
    ↪ Fig 1d','Range','G2')
writematrix(position_GenEvoSp_A','Supplementary_Data_4_ifaa_ifc_simulation.xlsx','sheet','simulation Fig
    ↪  1d','Range','J2')
writematrix(position_GenEvoSp_E','Supplementary_Data_4_ifaa_ifc_simulation.xlsx','sheet','simulation Fig
    ↪  1d','Range','M2')
position_ProEvoSp_Vl=position_ProEvoSp_all(:,NumGenPro+1:5*NumGenPro);
position_ProEvoSp_Bh=position_ProEvoSp_all(:,5*NumGenPro+1:6*NumGenPro);
position_ProEvoSp_Bl=position_ProEvoSp_all(:,6*NumGenPro+1:7*NumGenPro);
position_ProEvoSp_A=position_ProEvoSp_all(:,7*NumGenPro+1:8*NumGenPro);
position_ProEvoSp_E=position_ProEvoSp_all(:,8*NumGenPro+1:9*NumGenPro);
writematrix(position_ProEvoSp_Vl','Supplementary_Data_4_ifaa_ifc_simulation.xlsx','sheet','simulation
    ↪ Fig 3d','Range','A2')
writematrix(position_ProEvoSp_Bh','Supplementary_Data_4_ifaa_ifc_simulation.xlsx','sheet','simulation
    ↪ Fig 3d','Range','D2')
writematrix(position_ProEvoSp_Bl','Supplementary_Data_4_ifaa_ifc_simulation.xlsx','sheet','simulation
    ↪ Fig 3d','Range','G2')
writematrix(position_ProEvoSp_A','Supplementary_Data_4_ifaa_ifc_simulation.xlsx','sheet','simulation Fig
    ↪  3d','Range','J2')
```

```
writematrix(position_ProEvoSp_E','Supplementary_Data_4_ifaa_ifc_simulation.xlsx','sheet','simulation Fig
    ↪  3d','Range','M2')
CW_Vl=CW_taxon_all(:,1:NumGenPro);
writematrix(CW_Vl','Supplementary_Data_4_ifaa_ifc_simulation.xlsx','sheet','simulation SFig 4d','Range',
    ↪ 'B2')

% Saving the simulation results for plotting figures (Fig. 1d, 3d; Supp. Fig. 1d, 3d)
CW_taxon_V_F_Glu_rich=CW_taxon_all(:,1:NumGenPro);
CW_taxon_V=CW_taxon_all(:,NumGenPro+1:5*NumGenPro);
CW_taxon_Bh=CW_taxon_all(:,5*NumGenPro+1:6*NumGenPro);
CW_taxon_Bl=CW_taxon_all(:,6*NumGenPro+1:7*NumGenPro);
CW_taxon_A=CW_taxon_all(:,7*NumGenPro+1:8*NumGenPro);
CW_taxon_E=CW_taxon_all(:,8*NumGenPro+1:9*NumGenPro);
% save('CW_taxon_V_F_Glu_rich.mat','CW_taxon_V_F_Glu_rich','-mat')
% save('CW_taxon_V.mat','CW_taxon_V','-mat')
% save('CW_taxon_Bh.mat','CW_taxon_Bh','-mat')
% save('CW_taxon_Bl.mat','CW_taxon_Bl','-mat')
% save('CW_taxon_A.mat','CW_taxon_A','-mat')
% save('CW_taxon_E.mat','CW_taxon_E','-mat')
ifaa_normal_V_F_Glu_rich=ifaa_normal_all(1:NumGenPro,:,:);
ifaa_normal_V=ifaa_normal_all(NumGenPro+1:5*NumGenPro,:,:);
ifaa_normal_Bh=ifaa_normal_all(5*NumGenPro+1:6*NumGenPro,:,:);
ifaa_normal_Bl=ifaa_normal_all(6*NumGenPro+1:7*NumGenPro,:,:);
ifaa_normal_A=ifaa_normal_all(7*NumGenPro+1:8*NumGenPro,:,:);
ifaa_normal_E=ifaa_normal_all(8*NumGenPro+1:9*NumGenPro,:,:);
% save('ifaa_normal_V_F_Glu_rich.mat','ifaa_normal_V_F_Glu_rich','-mat')
% save('ifaa_normal_V.mat','ifaa_normal_V','-mat')
% save('ifaa_normal_Bh.mat','ifaa_normal_Bh','-mat')
% save('ifaa_normal_Bl.mat','ifaa_normal_Bl','-mat')
% save('ifaa_normal_A.mat','ifaa_normal_A','-mat')
% save('ifaa_normal_E.mat','ifaa_normal_E','-mat')

disp('Finished')
```

### Supplementary references


[1] Shepherd, J.C.W. (1981) Periodic correlation in DNA sequences and evidence suggesting their evolutionary origin in a comma-less genetic code. J. Mol. Evol. 17:94-102.

[2] Yin, C., Yau, S.-T. (2005) A Fourier characteristic of coding sequences: origins and a non-Fourier approximation. J. Comput. Biol. 12(9):1153-1169.

[3] Yin, C., Yau, S.-T. (2007) Prediction of protein coding regions by the 3-base periodicity analysis of a DNA sequences. J. Theor. Biol. 247:687-694.

[4] Ahmad, M., Jung, L.T., Bhuiyan, A.A. (2017) From DNA to protein: why genetic code context of nucleotides for DNA signal processing? a review. Biomed. Sign. Proc. Contr. 34:44-63.

[5] Bonidia, R.P., Sampaio, L.D.H., Domingues, D.S., et al. (2021) Feature extraction approaches for biological sequences: a comparative study of mathematical features. Brief. Bioinform. 22(5):1-20.

[6] Li, D.J. (2022) Distributional features of triplet codons in genomes underlie the diversification of life. Biosystems 217: 104681.

[7] Geoffroy Saint-Hilaire, É. (1838) Notions synthétiques, historiques et physiologiques de philosophie naturelle. Dénain, Paris.

[8] Iurato, G., Igamberdiev, A.U. (2021) Étienne Geoffroy Saint-Hilaire as a predecessor of the epigenetic concept of evolution. Biosystems 210:104571.

[9] Galera, A. (2021) Étienne Geoffroy Saint-Hilaire and the first embryological evolutionary model on the origin of vertebrates. J. Hist. Biol. 54:229-245.

[10] Cottam, R., Iurato, G., Igamberdiev, A.U. (2022) Fundamentals of evolutionary transformations in biological systems. Biosystems 222:104779.

[11] Zhou, P., Yang, X.L., Wang, X.G., Hu, B., Zhang, L., Zhang, W., Si, H., Zhu, Y., Li, B., Huang, C., Chen, H., Chen, J., Luo, Y., Guo, H., Jiang, R., Liu, M., Chen, Y., Shen, X., Wang, X., Zheng, X., Zhao, K., Chen, Q., Deng, F., Liu, L., Yan, B., Zhan, F., Wang, Y., Xiao, G., Shi, Z. (2020) A pneumonia outbreak associated with a new coronavirus of probable bat origin. Nature 579:270-273.

[12] Bloch, D.P., McArthur, B., Mirrop, S. (1985) tRNA-rRNA sequence homologies: Evidence for an ancient modular format shared by tRNAs and rRNAs. Biosystems 17:209-225.

[13] Nazarea, A.D., Bloch, D.P., Semrau, A.C. (1985) Detection of a fundamental modular format common to transfer and ribosomal RNAs: second-order spectral analysis. Proc. Natl. Acad. Sci. U. S. A. 82:5337-5341.

[14] Rodin, A.S., Szathmáry, E., Rodin, S.N. (2011) On origin of genetic code and tRNA before translation. Biol. Direct. 22:6-14.

[15] Frenkel, Z.M., Trifonov, E.N. (2012) Origin and evolution of genes and genomes. Crucial role of triplet expansions. J. Biomol. Struct. Dyn. 30:201-210.

[16] Frenkel, Z.M., Barzily, Z., Volkovich, Z., Trifonov, E.N. (2013) Hidden ancient repeats in DNA: Mapping and quantification. Gene 528:282-287.

[17] Kravatskaya, G.I., Kravatsky, Y.V., Chechetkin, V.R., Tumanyan, V.G. (2011) Coexistence of different base periodicities in prokaryotic genomes as related to DNA curvature, supercoiling, and transcription. Genomics 98:223-231.

[18] Ohno, S., Epplen, J. T. (1983) The primitive code and repeats of base oligomers as the primordial protein-encoding sequence. Proc. Natl. Acad. Sci. U. S. A. 80:3391-3395.

[19] Ohno, S. (1984) Repeats of base oligomers as the primordial coding sequences of the primeval earth and their vestiges in modern genes. J. Mol. Evol. 20:313-321.

[20] Shiba, K., Takahashi, Y., Noda, T. (2002) On the role of periodism in the origin of proteins. J. Mol. Biol. 320:833-840.

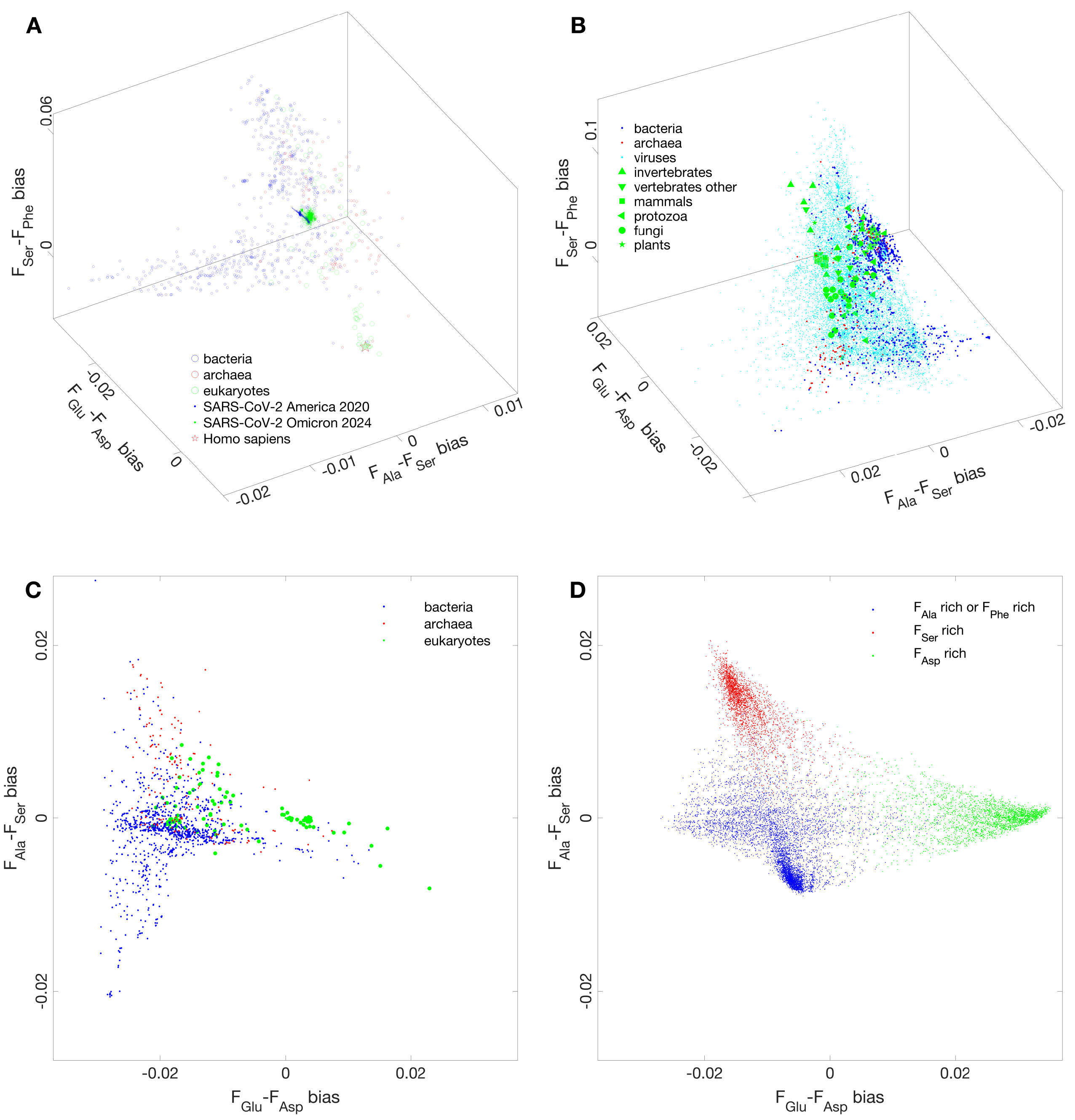


**Supp. Fig. 1a.** Supplementary figures to Fig. 1a. **A** Both the three domains of life and SARS-CoV-2 in GenEvoSp. **B** Bacteria with high GC content, bacteria with low GC content, archaea, and eukaryotes occupying the four vertices of the tetrahedron, respectively, in GenEvoSp, and viruses occupying the inner region of the tetrahedron. **C** The direction from bacteria with high GC content to archaea is parallel to the axis of $F_{Ala} - F_{Ser}$ bias. The direction from archaea to eukaryotes is approximately parallel to the axis of $F_{Glu} - F_{Asp}$ bias, with a small included angle. **D** Simulation of the distribution of life in GenEvoSp agrees approximately with the observation based on real life. There is an unexplained small angle of rotation between simulation and observation.

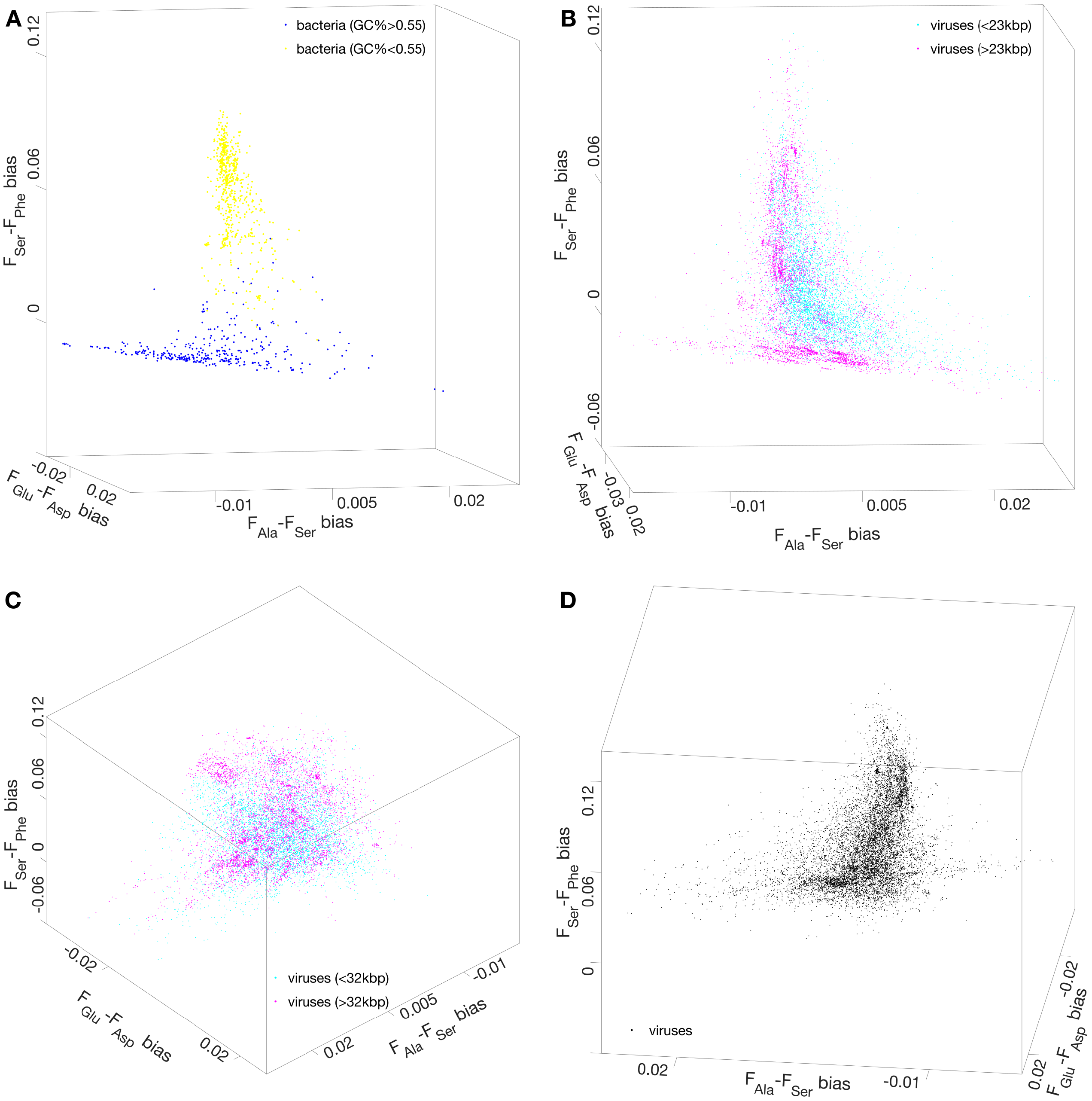


**Supp. Fig. 1b.** Supplementary figures to Fig. 1b. **A** The direction from bacteria with high GC content to bacteria with low GC content in GenEvoSp. **B** The direction from viruses with small genome size to viruses with large genome size in GenEvoSp. **C** Plaid pattern of virus distribution in GenEvoSp. **D** These high-density distribution regions of viruses are arranged in sequence.

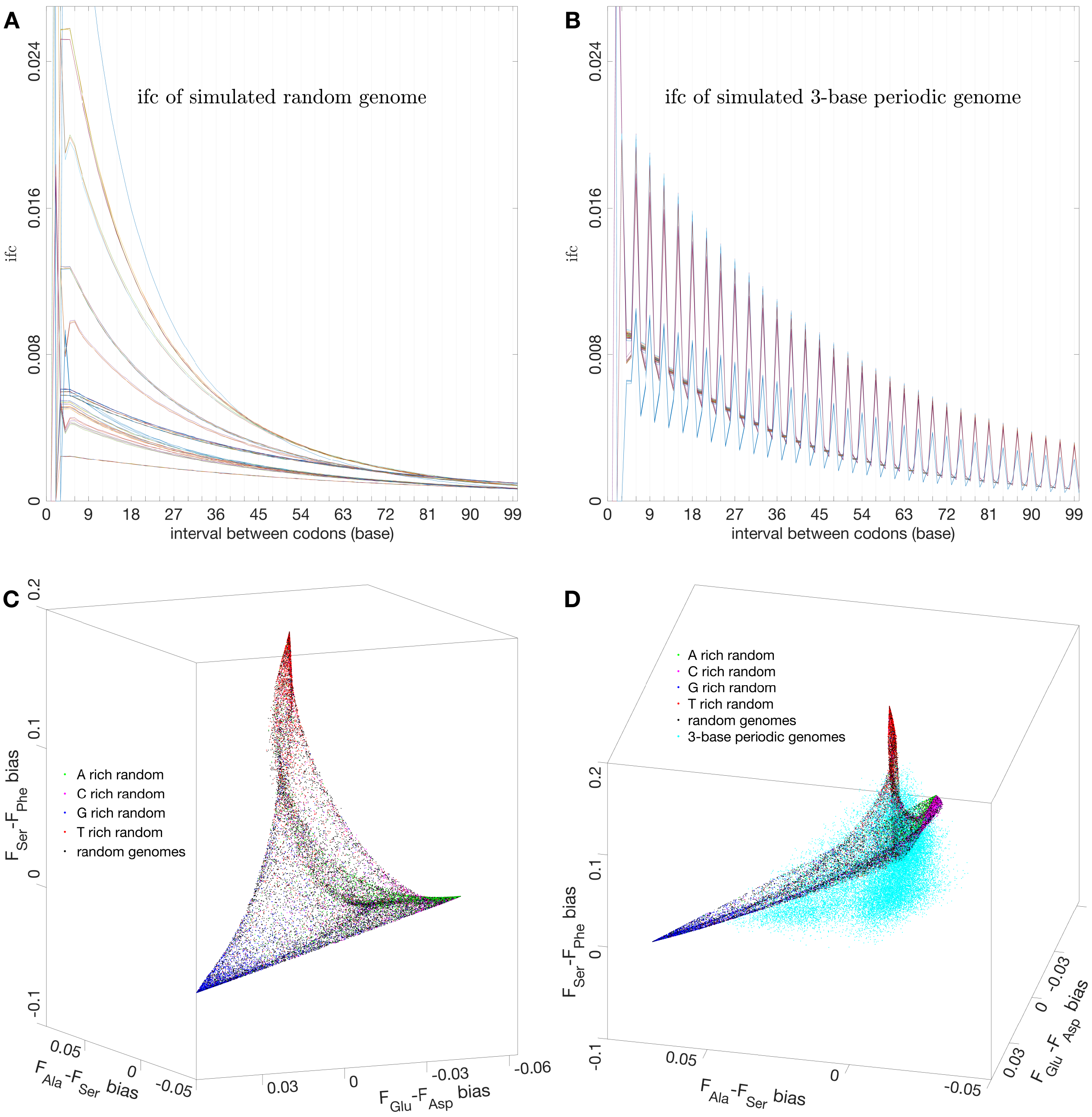


**Supp. Fig. 1c.** Supplementary figures to Fig. 1c. **A** Random sequences exhibit no periodicity, let alone three-base periodicity. **B** Simulation of TBP via concatenation of randomly selected elements from the subsets of the 64 canonical codons. **C** The distribution region of simulated random-sequence genomes forms a trident star, which differs significantly from the tetrahedron of real biological genomes. **D** The three-base periodic genomes adopt an efficient mechanism to distinguish themselves from random sequences, whose tetrahedral distribution region deviates from the curved surface of the trident star.

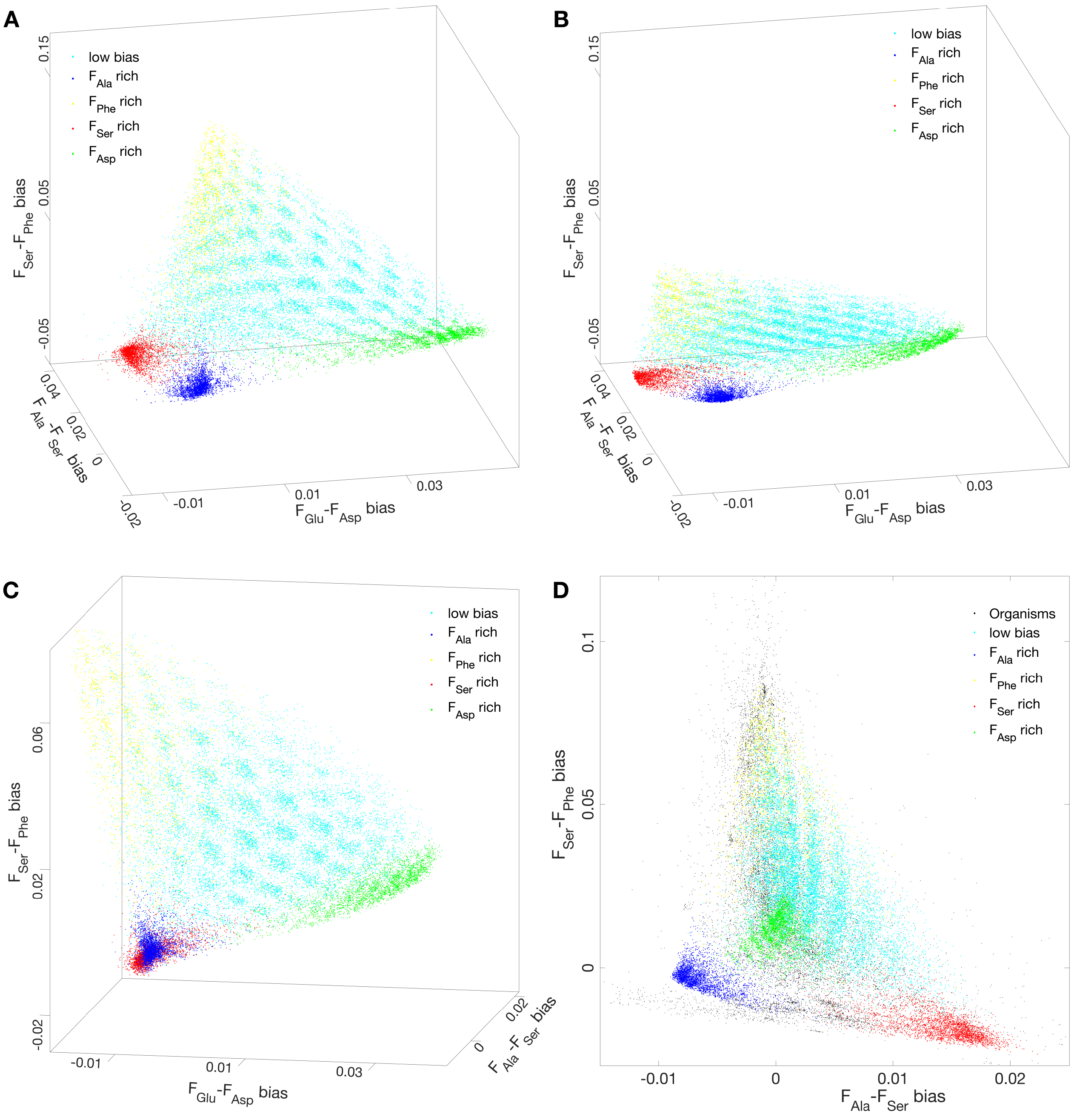


**Supp. Fig. 1d.** Supplementary figures to Fig. 1d. **A** When only a single strand of DNA is considered, the tetrahedron exhibits a positional shift between simulation and observation. **B** The mismatch described in (A) has been corrected by reading both complementary strands of DNA. **C** The plaid pattern of real life in GenEvoSp has been simulated. **D** The simulated tetrahedron approximately matches the tetrahedron for real life in terms of dimensions and spatial orientations.

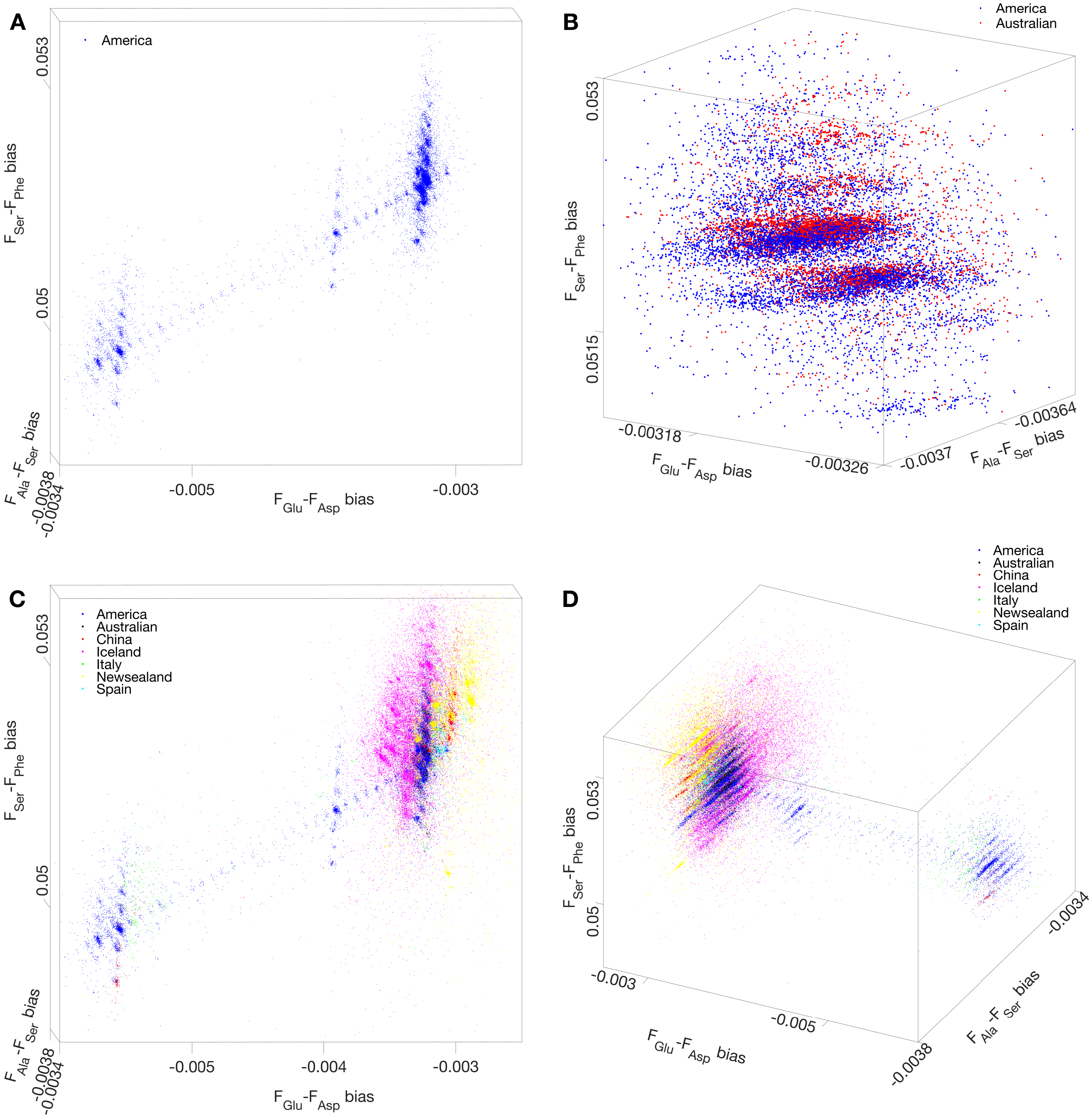


**Supp. Fig. 2a.** Supplementary figures to Fig. 2a. **A** America Kite with short-tail array, middle-tail array, and long-tail array in GenEvoSp. **B** The long-tail array of America Kite and the Australia Kite are misaligned and mutually overlapped. **C** Distribution of SARS-CoV-2 in GenEvoSp: Kites of different geographical regions. **D** Alternative perspective for the observation in (C).

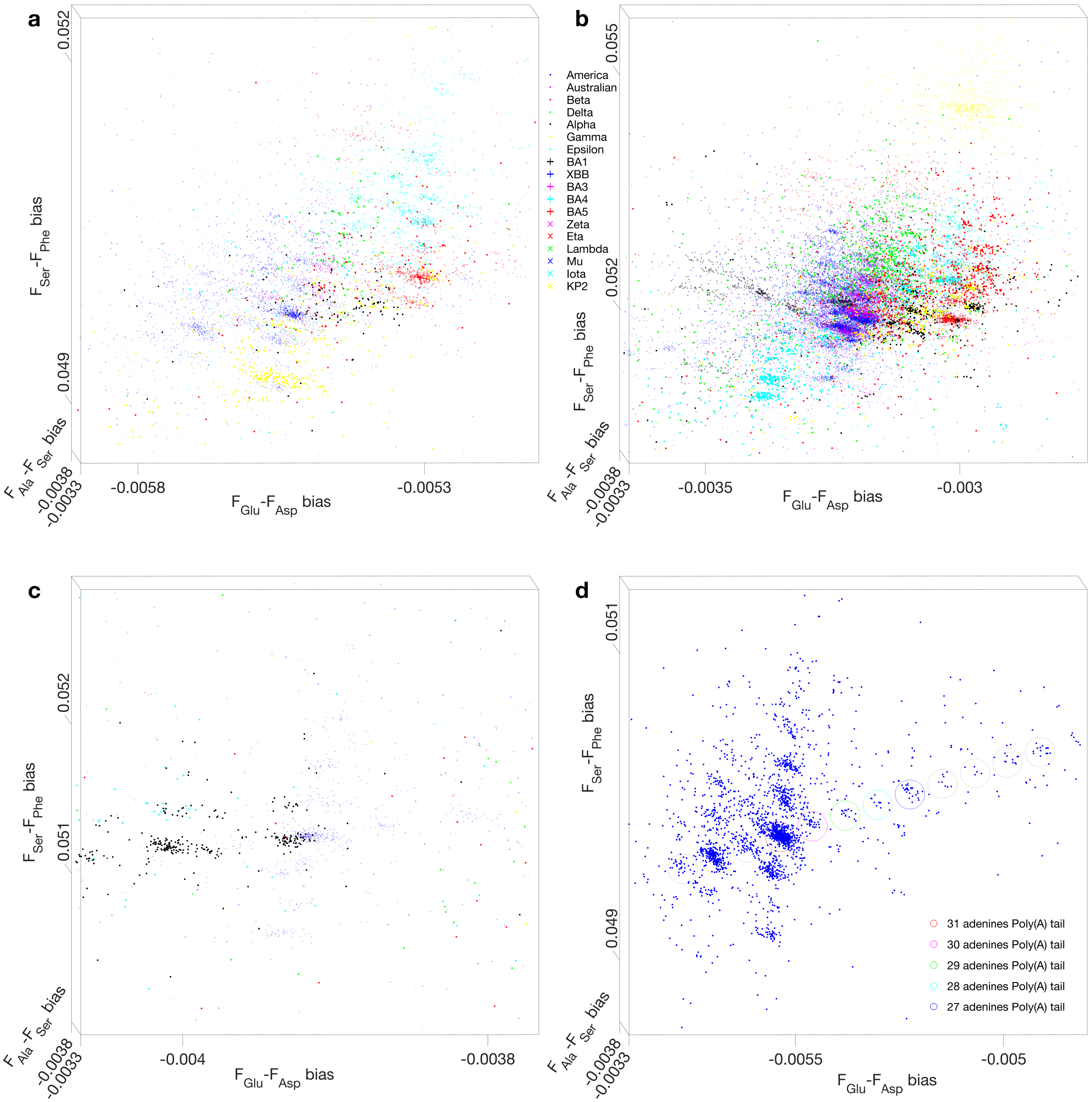


**Supp. Fig. 2b.** Supplementary figures to Fig. 2b. **A** The plaid pattern in the long-tail array of America Kite in GenEvoSp. **B** The plaid pattern in the short-tail array of America Kite in GenEvoSp. **C** The plaid pattern in the middle-tail array of America Kite in GenEvoSp. **D** Along the symmetry axis of the America Kite, adjacent subclusters differ by exactly 1 adenine residue in their poly(A) tail length.

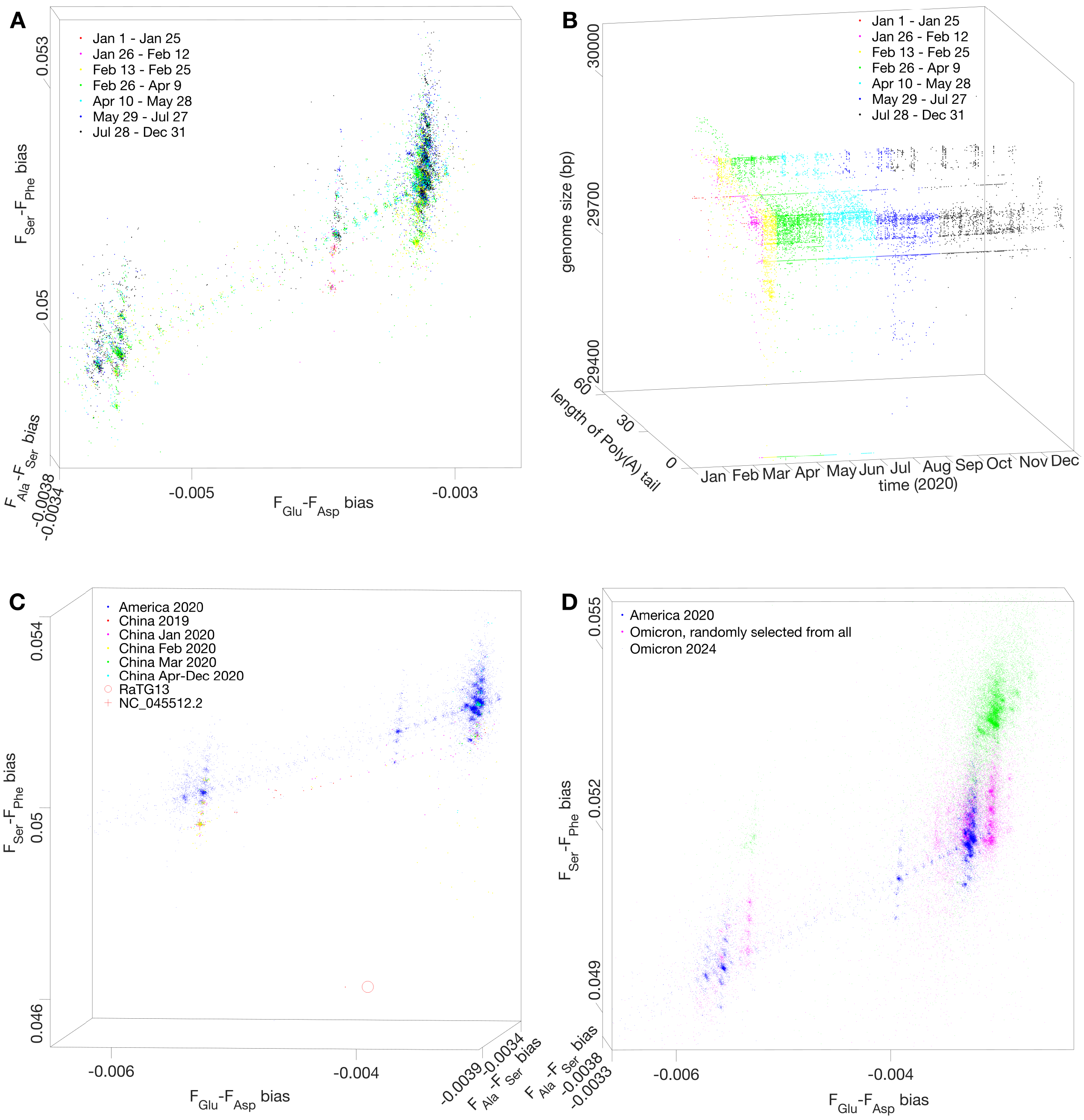


**Supp. Fig. 2c.** Supplementary figures to Fig. 2c. **A** The formation process of the America Kite, with distinct colors assigned to different sampling periods. The sampling date of each SARS-CoV-2 genome can be resolved at several days resolution. **B** The structural formation of the America Kite, where the sampling date of each SARS-CoV-2 genome is annotated at daily resolution along the dedicated time axis, while the third orthogonal axis represents the genome size of each SARS-CoV-2 genome. **C** A distinct parallel shift exists between the symmetry axis of the China Kite and that of the America Kite. **D** An evolutionary trend from America 2020 Kite to Omicron Kite.

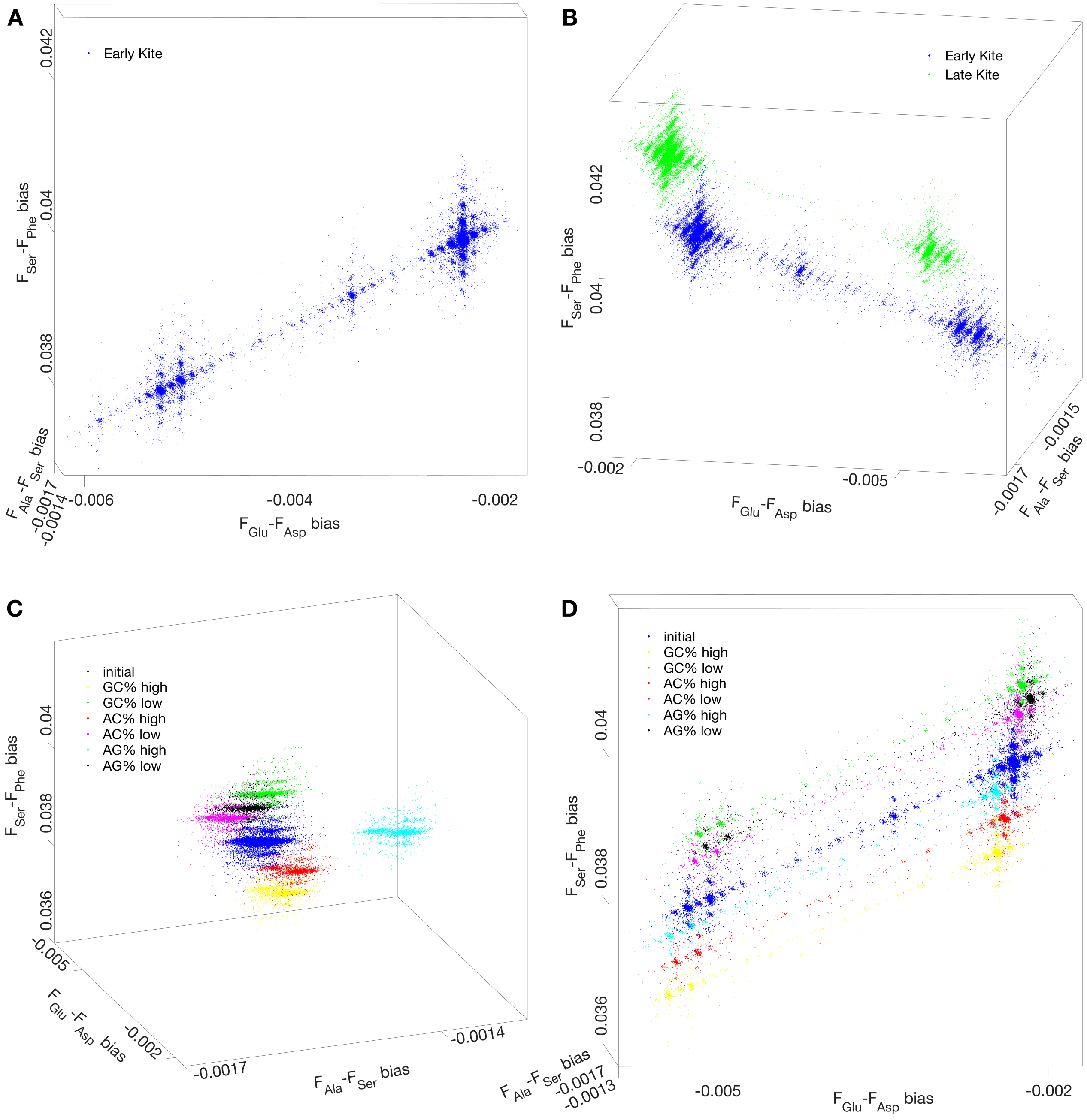


**Supp. Fig. 2d.** Supplementary figures to Fig. 2d. **A** Simulation of kite-like distribution of SARS-CoV-2 in GenEvoSp. **B** Simulation of evolutionary trend of Kites in GenEvoSp. **C** Kite drifts in GenEvoSp, due to variation of base composition, according to the simulation. **D** Alternative perspective for the observation in (C).

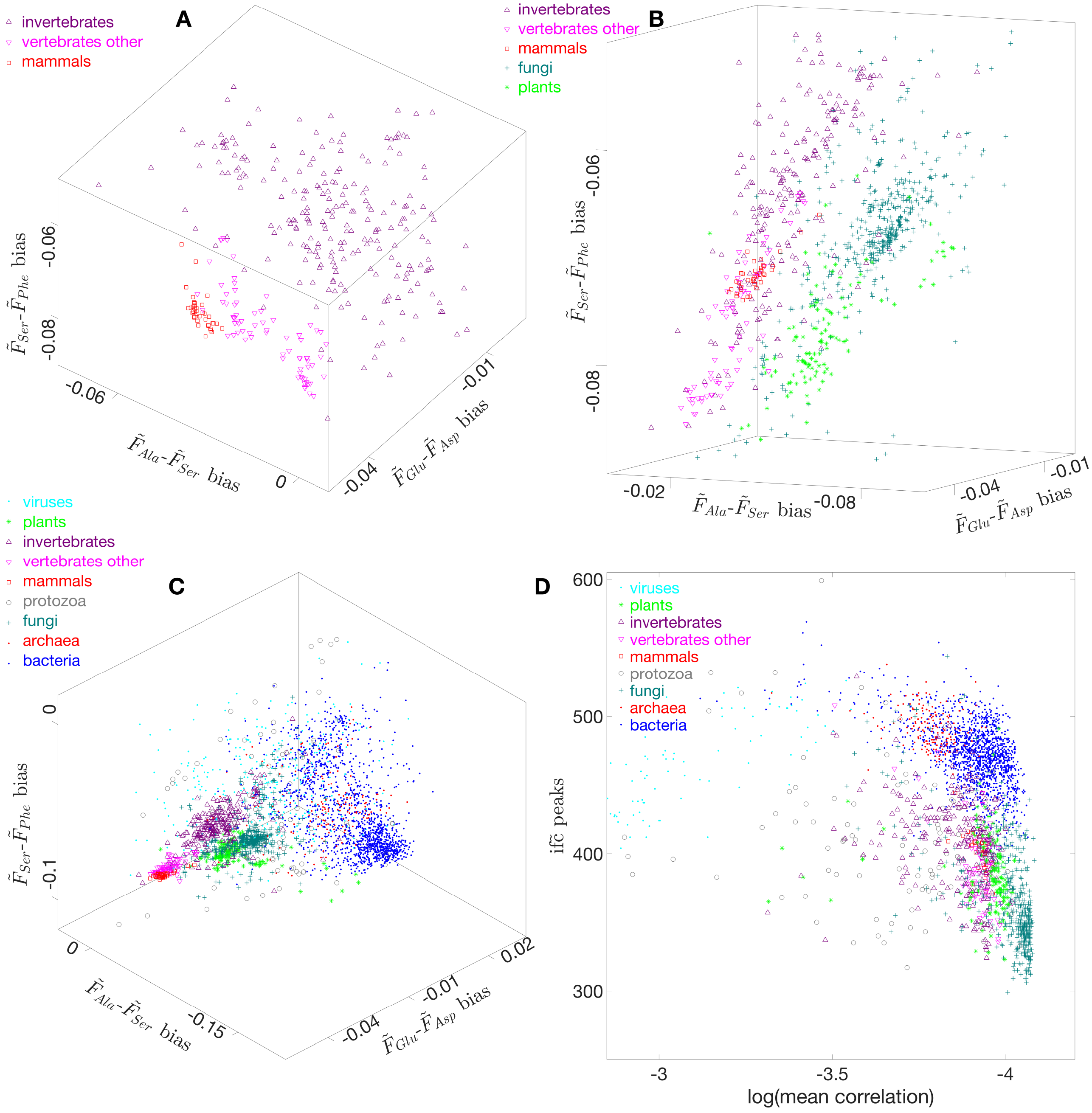


**Supp. Fig. 3a.** Supplementary figures to Fig. 3a. **A** In ProEvoSp, invertebrates, vertebrates excluding mammals, and mammals extend along the same direction. **B** Plants and animals exhibit distinct separate clustering distributions in ProEvoSp. **C** Separate clustering distribution of life in ProEvoSp. **D** Separate clustering distribution of the three domains of life based on the number of peaks in $if\tilde{c}$ and their mean correlation coefficients.

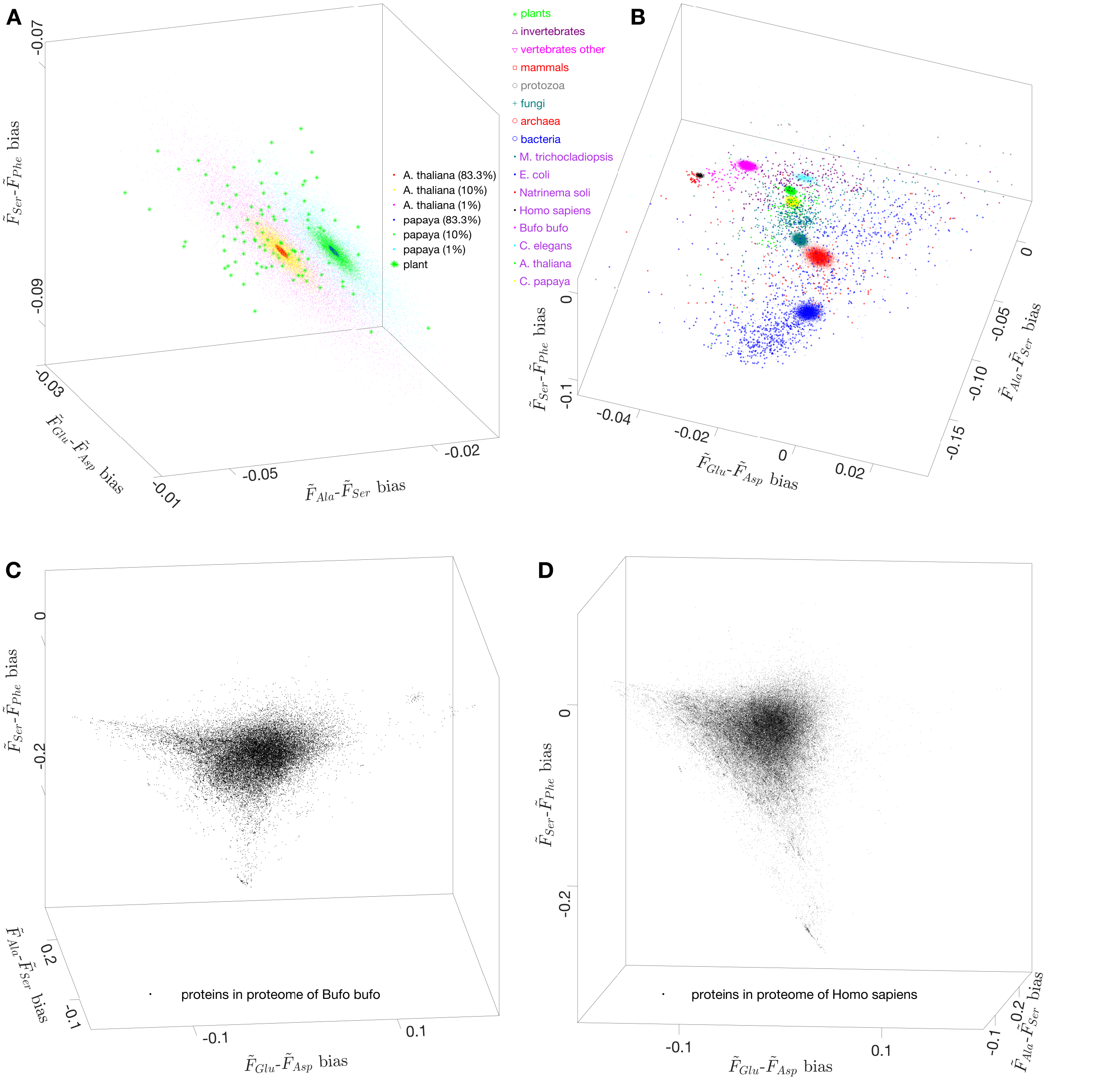


**Supp. Fig. 3b.** Supplementary figures to Fig. 3b. **A** Distribution of randomly selected proper subsets of a proteome in ProEvoSp, with colors marking subsets that account for 83.3%, 10% and 1% of the full proteome of a given species. The distribution orientation of these proteome subsets from A. thaliana or papaya agrees with the orientation of distribution of plants. **B** Distribution of randomly selected proper subsets of the proteome of a species across different taxa in ProEvoSp. **C** Distribution of all protein sequences from Bufo bufo in ProEvoSp, where the distribution density is significantly elevated in several small regions. **D** Distribution of all protein sequences from Homo sapiens in ProEvoSp, with markedly increased distribution density observed in specific small regions that are arranged in sequential rows.

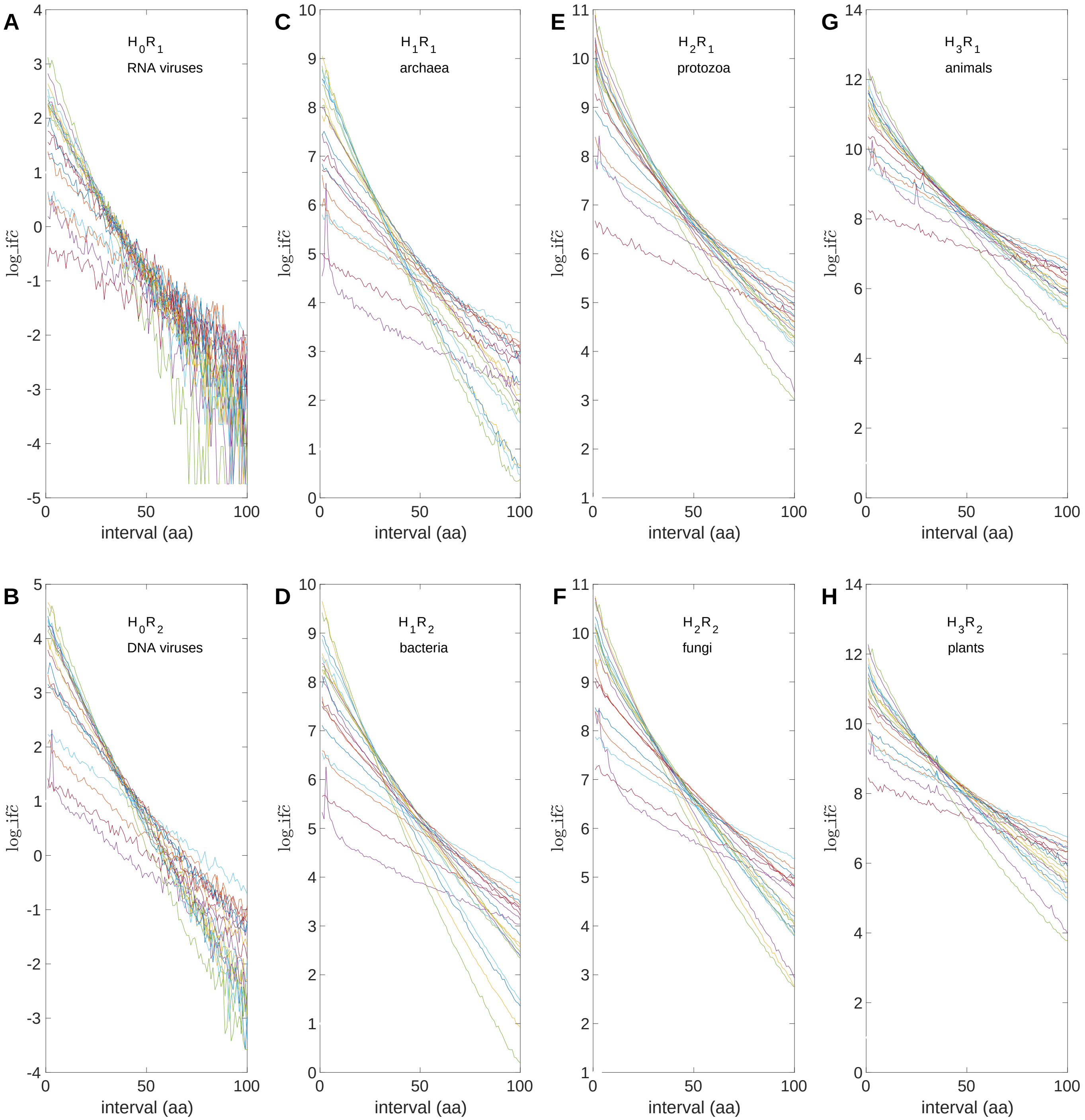


**Supp. Fig. 3c.** Supplementary figures to Fig. 3c. **A** Logarithm of $if\tilde{c}$ for RNA viruses. **B** Logarithm of $if\tilde{c}$ for DNA viruses. **C** Logarithm of $if\tilde{c}$ for archaea. **D** Logarithm of $if\tilde{c}$ for bacteria. **E** Logarithm of $if\tilde{c}$ for protozoa. **F** Logarithm of $if\tilde{c}$ for fungi. **G** Logarithm of $if\tilde{c}$ for animals. **H** Logarithm of $if\tilde{c}$ for plants.

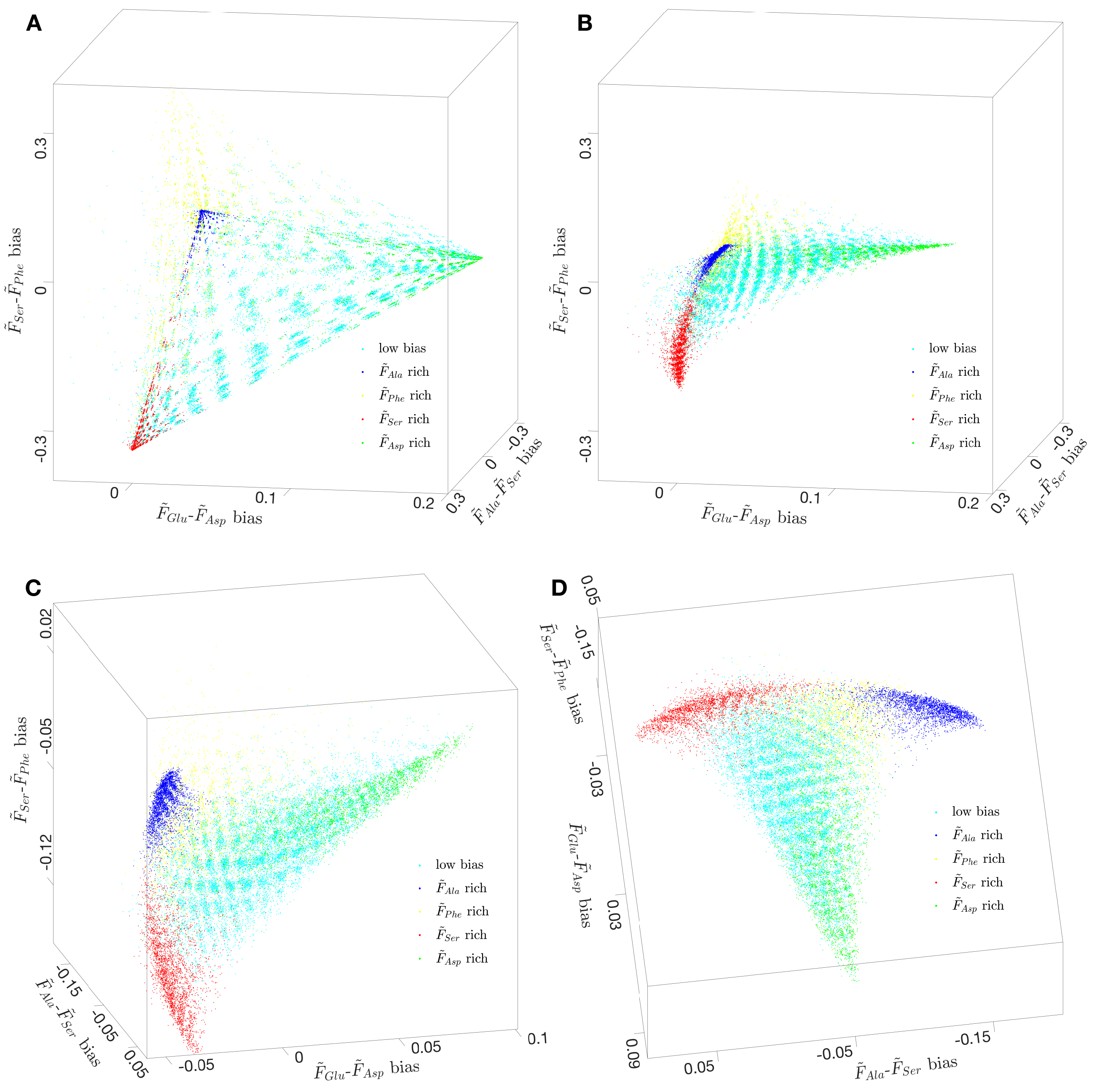


**Supp. Fig. 3d.** Supplementary figures to Fig. 3d. **A** The tetrahedron distribution in ProEvoSp of simulated proteomes translated from only one reading frame of the aforementioned simulated genomes. **B** The triangular distribution in ProEvoSp of simulated proteomes translated via three reading frames on a single DNA strand of the aforementioned simulated genomes. **C** The plaid pattern in ProEvoSp for real life has been simulated. **D** Alternative perspective for the observation in (C).

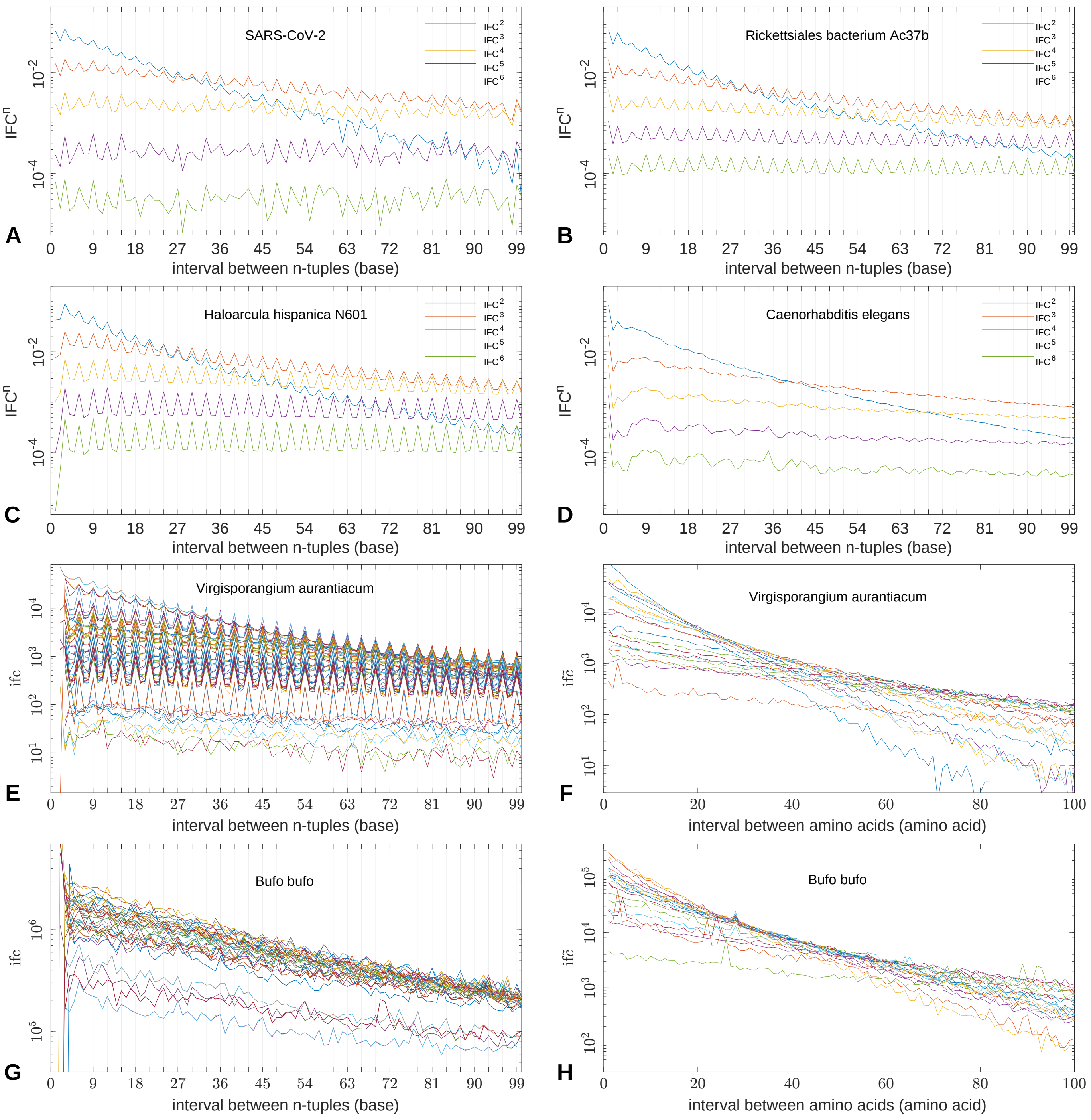


**Supp. Fig. 4a.** Supplementary figures to Fig. 4. **A** TBP for a virus. **B** TBP for a bacterium. **C** TBP for an archaeon. **D** TBP for a eukaryote. **E** $ifc$ of Virgisporangium aurantiacum. **F** $if\tilde{c}$ of Virgisporangium aurantiacum. **G** $ifc$ of Bufo. **H** $if\tilde{c}$ of Bufo.

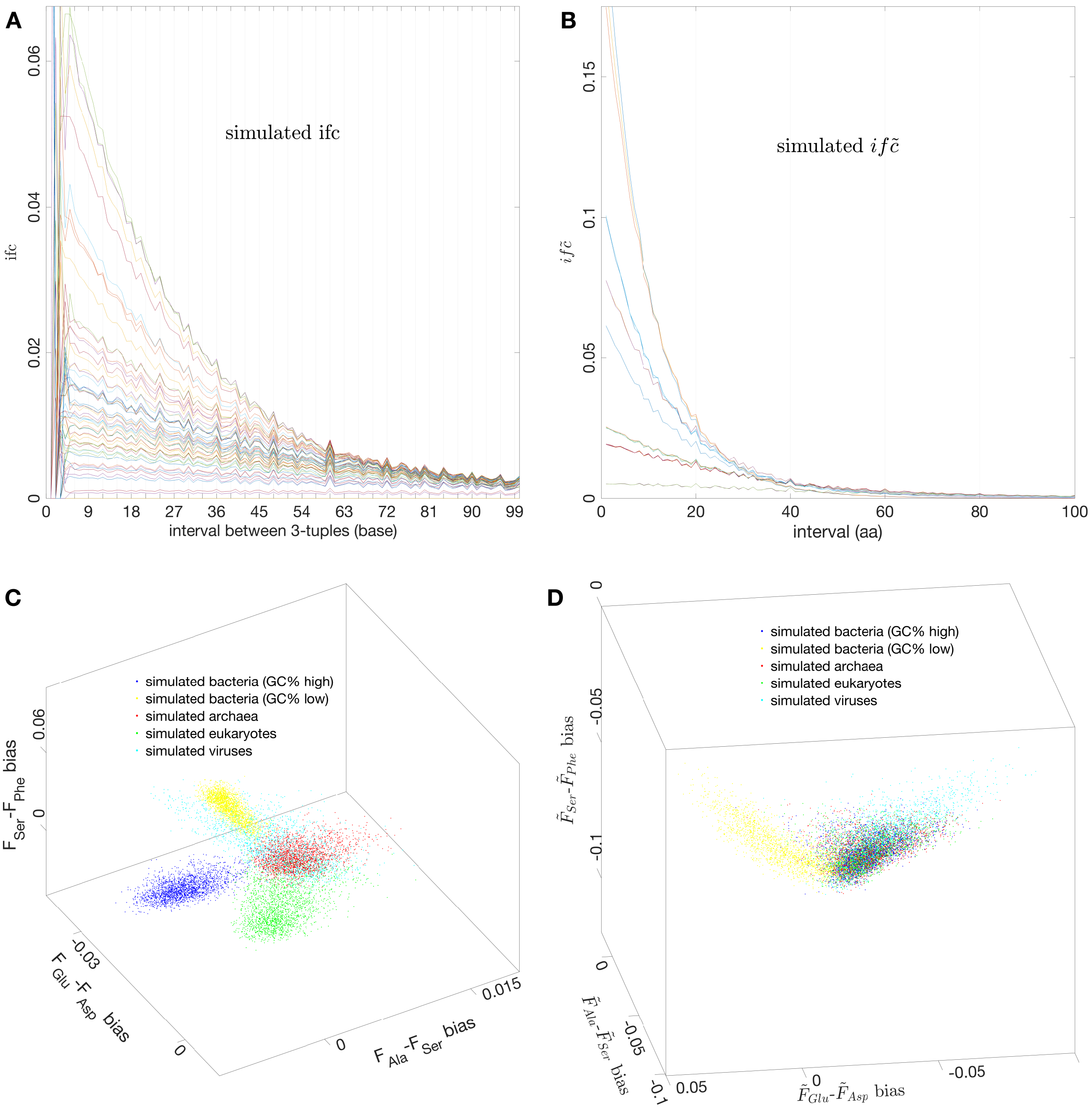


**Supp. Fig. 4b.** Supplementary figures to Fig. 4. **A** Simulation of initial TBP via a period averaging procedure. **B** The $if\tilde{c}$ based on proteome translated from the same genome used to calculate $ifc$ in (A). **C** Explanation of the diversification of the three domains of life in GenEvoSp, based on numerous genomes generated by substitutions of bases between amino acid biosynthetic families. **D** Explanation of the diversification of life in ProEvoSp based on numerous proteomes translated from the genomes in (C), which indicates that bacteria might have diverged prior to the establishment of the standard genetic code.

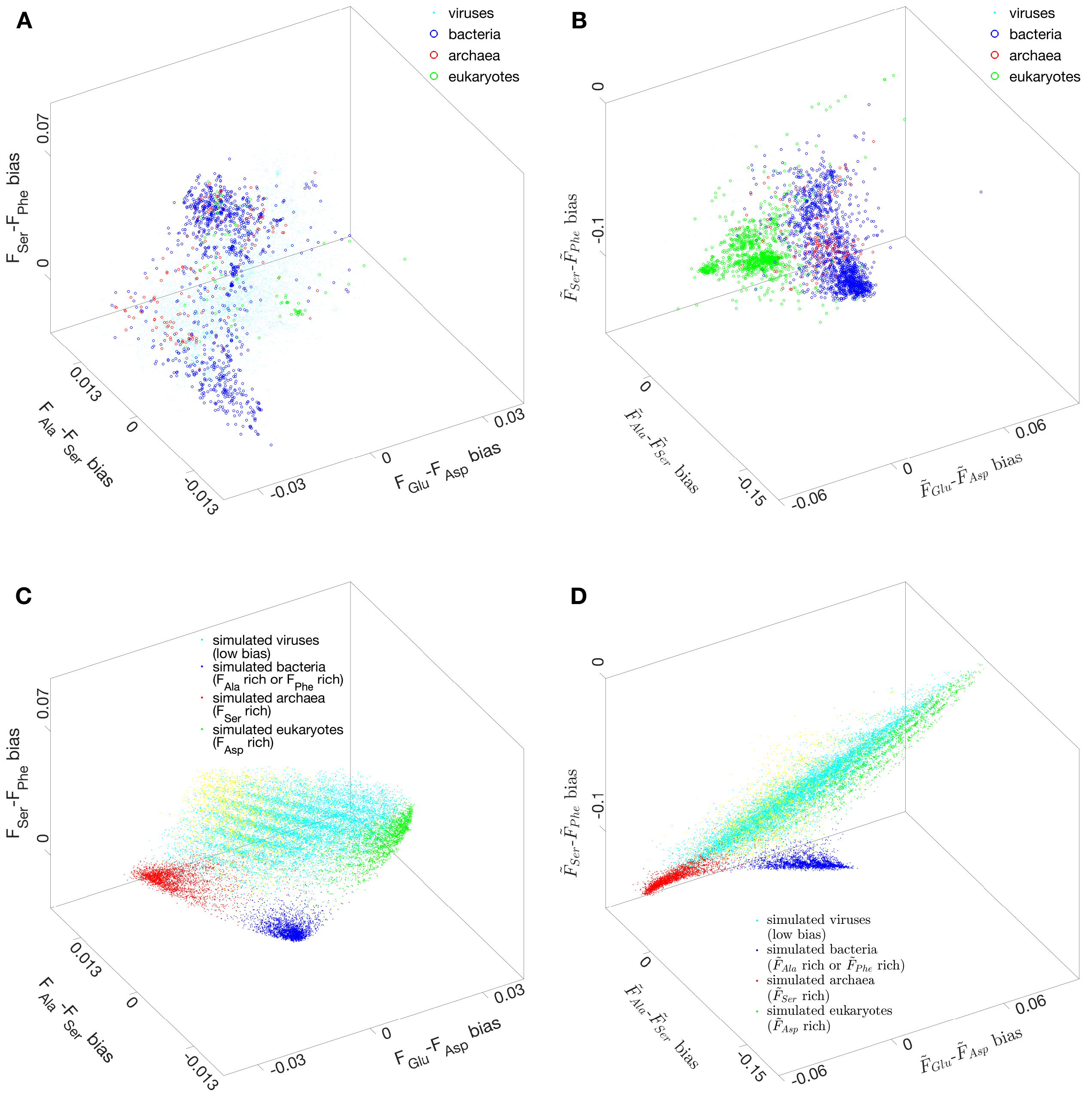


**Supp. Fig. 4c.** Comparison of the genome/proteome evolution space between observation and simulation. **A** Orientations of the three domains of life and virus in GenEvoSp for real life. **B** Orientations of the three domains of life and virus in ProEvoSp, which partially agree with the corresponding orientations in (A). **C** Orientations of the simulated three domains of life and virus in GenEvoSp, which agree with the orientations in (A). **D** Orientations of the simulated three domains of life and virus, in ProEvoSp, whose proteomes are translated from the simulated genomes in (C).

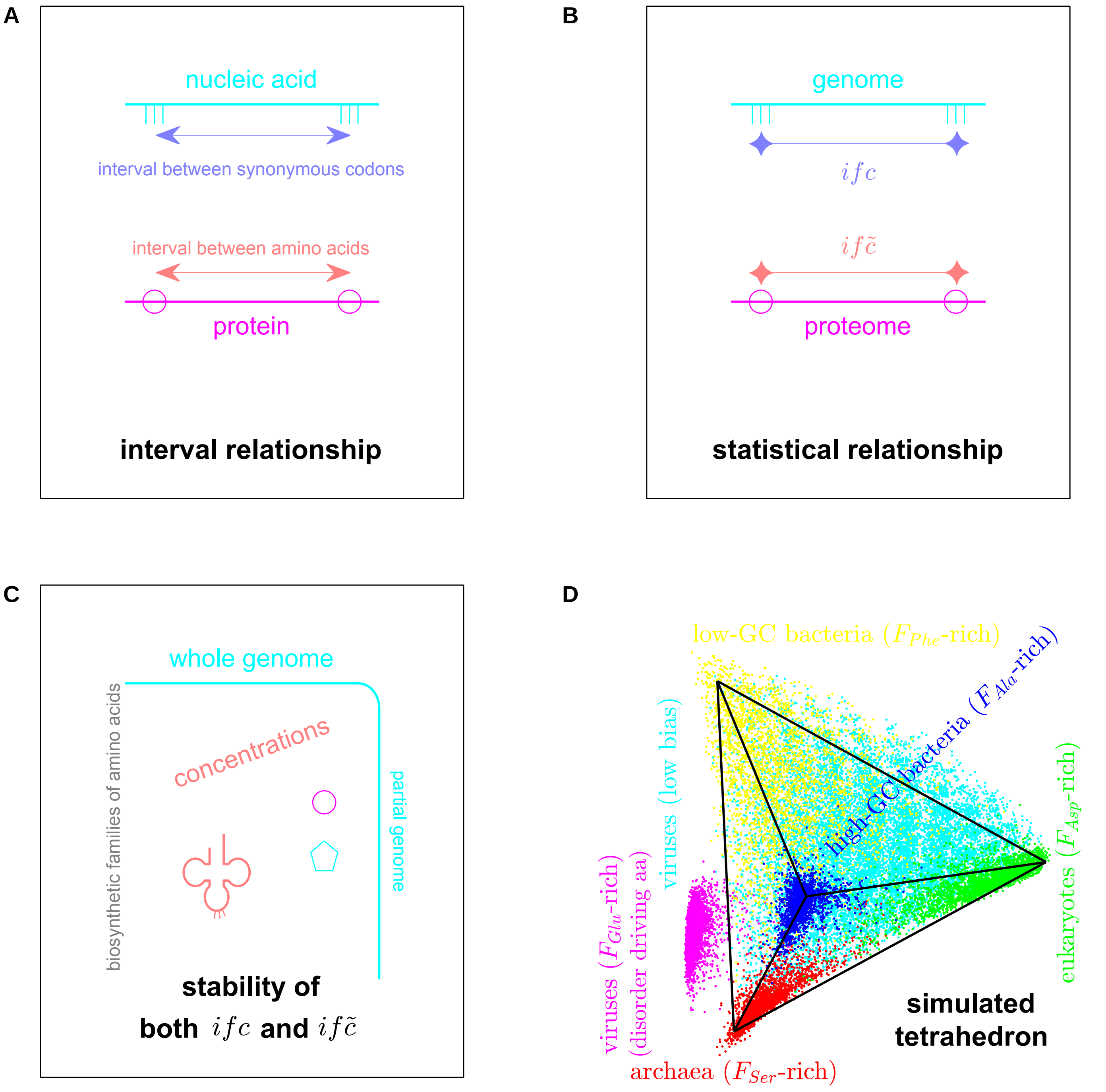


**Supp. Fig. 4d.** Statistical properties and the global picture of three-base periodic genomes. **A** There is a direct relationship between the spacing of adjacent triplet bases in a genome and the spacing of the corresponding amino acid residues in the corresponding proteome. **B** There is a statistical relationship between the distribution of triple base intervals in a genome and distribution of the corresponding amino acid residue intervals in the corresponding proteome. **C** There are interdependent relationships among $ifc$, $if\tilde{c}$, and tRNA concentrations. **D** The simulated tetrahedron agrees with the tetrahedron of real life (Fig. 1a).
